# *k*-spaces: Mixtures of Gaussian latent variable models

**DOI:** 10.1101/2025.11.24.690254

**Authors:** Nicholas Markarian, Barbara E. Engelhardt, Niles A. Pierce, Paul W. Sternberg, Lior Pachter

## Abstract

Principal component analysis (PCA) and *k*-means clustering are two seemingly different methods for dimension reduction and clustering, respectively, but can be understood as special cases of inference in a Gaussian latent variable model framework. We leverage this insight to develop a probabilistic framework and methods for simultaneous dimension reduction, clustering, and latent space learning that are efficient and interpretable, and that can replace current ad hoc combinations of PCA and clustering. The algorithm, *k*-spaces, has broad applicability, which we demonstrate in several distinct genomic settings. In particular, we show how *k*-spaces can be used to model gene expression in quantitative hybridization chain reaction (qHCR) images, for inference in epigenomics, and for dimension reduction of single-cell RNA-sequencing data.

## Introduction

Dimension reduction is an essential step in the analysis of large tabular data for exploration and visualization of patterns in the data. Two classical and useful dimension reduction techniques for matrices are principal component analysis (PCA) and *k*-means clustering, which reduce the dimensions of features and partition observations, respectively. Although seemingly unrelated methods, both PCA and *k*-means can be viewed as special cases of an algorithm that iteratively clusters samples while reducing the dimension of the feature space by fitting a set of affine subspaces (i.e., points, lines, planes, etc.) to the features in order to capture distinct linear relationships within subpopulations in the data.

PCA may be derived as a limiting case, where the variance approaches zero, of a probabilistic homoskedastic latent Gaussian factor model called probabilistic PCA (pPCA) (1, 2). This formulation leads to an interpretation of PCA as the optimal inference procedure (in the limit) to find a latent space for points perturbed by isotropic Gaussian noise. From this point of view, zero-dimensional pPCA fits a single isotropic Gaussian to the data. In a Gaussian mixture model framework, it is therefore natural to consider *k* mixtures of Gaussian latent variable models, each possibly of a different dimension, with isotropic noise in their complementary spaces. We derive an algorithm to perform inference for such models that contains Gaussian mixture models and pPCA as special cases, and we call this framework *k*-spaces. These specific models were considered in earlier work (1–5); in particular, Bishop and Tipping explored the extension of their pPCA to mixture models for piecewise local linear dimension reduction of nonlinear manifolds (6), but their model has not been widely explored or deployed for clustering to the best of our knowledge. We develop a framework for fitting subspaces of mixed dimension and shared noise parameters as well as performing model selection, which makes *k*-spaces suitable for a variety of biological settings where it is useful to simultaneously cluster and reduce dimension.

Our work was motivated by several problems in biology that require fitting multiple linear latent spaces to data. One such problem is detecting distinct gene coexpression relationships within different cell types or states. We use *k*-spaces to analyze a quantitative hybridization chain reaction (qHCR) image, where our results and *k*-spaces’s behavior can be interpreted in an anatomical and developmental context. qHCR imaging overcomes multiple shortcomings of traditional enzyme-based imaging methods to enable multiplex, quantitative, high-resolution RNA imaging, yielding subcellular voxel intensities that scale linearly with target abundance (7–9). In prior work (7), a qHCR image staining the expression levels of four mRNAs was used to identify the spatial organization of distinct circuit states active during zebrafish somitogenesis. In that study, circuit states for regions of interest (ROIs) were identified by manual clustering of their gene expression vectors across several pairwise scatter plots and then analyzed in spatial context by shading voxels of the image within ROIs by the identified circuit states. Here, we use *k*-spaces to process the entire image and automatically fit *k* lines to the entire dataset in the full 4-D voxel intensity space. We show that this automated approach to the analysis technique introduced by that study both recapitulates and extends earlier findings (7). By analyzing all cells in the image at once, we identify spatial usage of distinct gene circuit states and their spatial gradients of expression scaling.

Our *k*-spaces method has immediate application to diverse problem types, as we demonstrate with additional examples from epigenomics and RNA sequencing. The epigenomics application, namely detection of differentially methylated sites between underlying cell types in bulk data, challenges *k*-spaces through the dimension of the clusters. Clustering on features rather than observations of the methylation matrix, we project uncorrelated and correlated CpG sites (i.e., the sites of methylation) onto separate 0- and 4-dimensional linear manifolds. In particular, sub-populations in the data are not linearly separable but are separable by the dimensionality of their subspaces, and our model lends itself to tractable theoretical analysis. Similarly, we show that *k*-spaces is a powerful tool for the analysis of high-dimensional single-cell RNA-sequencing (scRNA-seq) data by virtue of unifying the dimension reduction and clustering steps, allowing us to visualize the distinct yet partially synchronized developmental trajectories of *C. elegans* hypodermis and seam cells across a series of linear dimension reductions.

## Results

### *k*-spaces overview

The *k*-spaces algorithm uses Gaussian latent variables and an expectation maximization (EM) procedure to learn a set of *k* latent spaces with dimension *d*_1_, *d*_2_, …, *d*_*k*_. Observed data points are assumed to be generated by a process with isotropic Gaussian noise in the complementary space, with either a distinct variance parameter for noise for each latent space or one that is shared for the entire model. Observations are iteratively assigned to subspaces (grouping rows of the data matrix) and an optimal linear dimension reduction is computed to represent each submatrix (reducing columns of the matrix; Fig. 1). *k*-spaces produces affine subspaces that cluster the observations along with projections of the clusters onto their subspaces. Depending on the application, different aspects of the *k*-spaces method may be of interest. For example, we show that the clustering of epigenomics data produces a classification of sites. In the gene coexpression analysis, the affine subspaces provide insights into gene expression scaling in qHCR. In single-cell RNA-sequencing data analyses, the latent spaces reveal complex structure in qHCR and single-cell relationships.

**Fig. 1.**
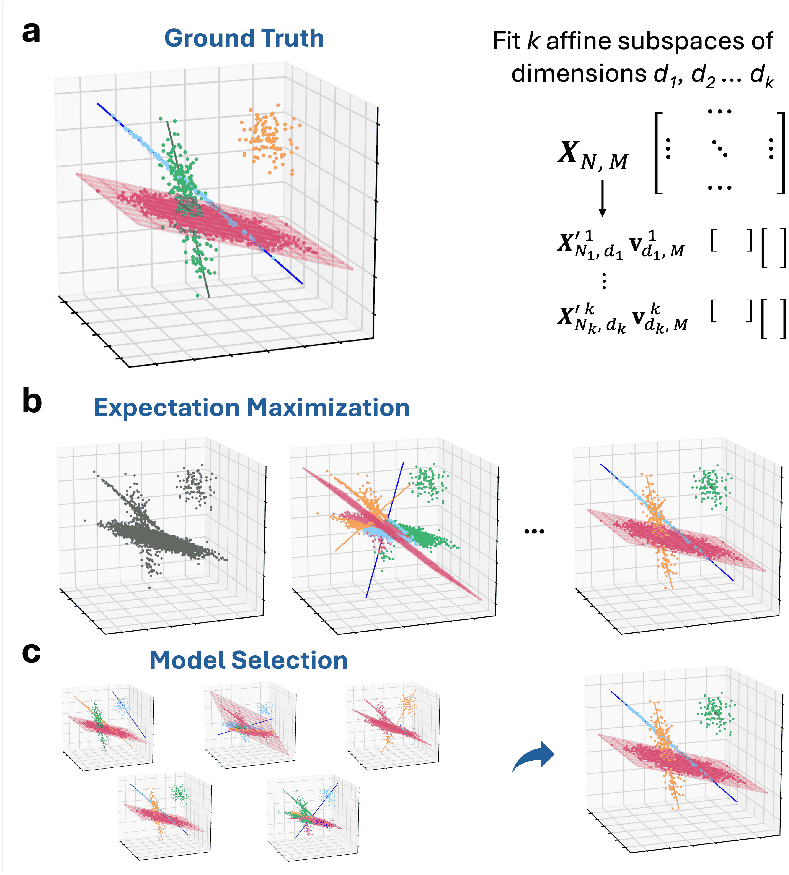
Overview of *k*-spaces clustering on simple synthetic data. **a**, Data are generated with isotropic noise from 1 plane, 2 lines, and 1 point. The *k*-spaces clustering algorithm will simultaneously partition and reduce the dimension of the data. **b**, Once a set of spaces is specified to cluster with (here, 1 plane, 2 lines, and 1 point), an expectation maximization algorithm is used to maximize the data likelihood. **c**, Model selection with the Integrated Completed Likelihood (ICL) or Bayesian information criterion (BIC) could be used to select the best model, both between models with spaces of the same dimension or models with different dimensions. Here, BIC was used to select between competing models. From left to right: 1 plane and 3 lines (yielding a correct classification of points but incorrect dimension of clusters); the correct solution from (b); a local minimum with 1 plane, 2 lines, and 1 point; 2 lines and 3 points; and another local minimum with 1 plane, 2 lines, and 1 point.

A self-contained description of our model and derivations of the inference procedures are provided in the Methods and Supplement.

### *k*-spaces clustering applied to qHCR data identifies distinct gene circuit states and their oscillating expression levels in the developing zebrafish embryo

We first evaluated the behavior of *k*-spaces on systems data reflecting heterogeneous cell types and gene circuit states using spatial embryogenesis data produced with qHCR. The read-out/read-in analysis framework we build upon enables bi-directional quantitative discovery (7): read-out from anatomical space to expression space to discover co-expression relationships; read-in from expression space to anatomical space to discover the anatomical regions in which selected gene co-expression relationships occur. Trivedi and coworkers did the following: in a *read-out* step, to identify circuit states for regions of interest (ROI) within the image, voxel intensities were displayed as six 2-D scatter plots (one plot for each pairwise combination of the four mRNAs assayed in their 4-plex qHCR image) and gene expression clusters of interest were identified manually. Second, in a *read-in* step, to identify the anatomical location of each circuit state within the embryo, the voxels within the image were shaded by their expression cluster index. Here, we revisit their analysis of four mRNAs active during zebrafish somitogenesis.

We started with the 4-plex qHCR image (Fig. 2a). Following normalization of the signal for each channel, we used *k*-spaces to read-out all 17,340 subcellular (2 × 2 × 6 µm) voxels to 4-D expression scatter plots and fit 9 lines to the voxel intensities directly within the 4-D space (Fig. 2b); each line is an automatically generated expression cluster corresponding to a circuit state with different mRNA stoichiometry. We then used *k*-spaces to read these expression clusters back into the anatomical context of the image, coloring voxels according to cluster index to automatically segment the anatomy based on circuit state (Fig. 2c).

**Fig. 2.**
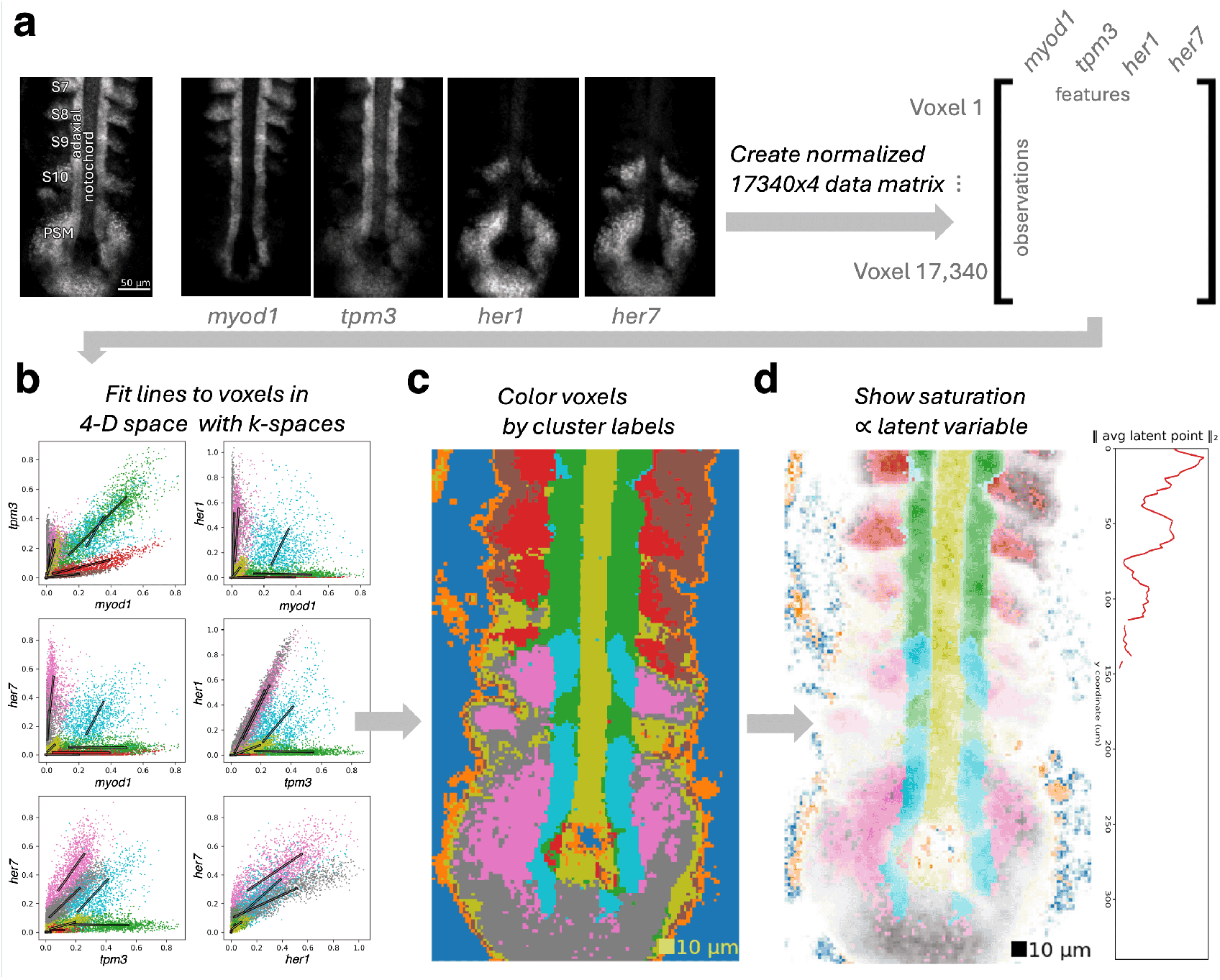
Automated read-out/read-in analysis of a qHCR image of mRNA expression during zebrafish somitogenesis. **a**, Anatomical features labeled in a grayscale merge (left) of a 4-channel quantitative image of four target mRNAs (middle); 4-channel data was extracted and normalized (right) using the Read-out/Read-in software with default settings (7) to produce a 4-D scatter plot of 17,340 voxel intensities. **b**, Clusters were then automatically identified in 4-D using *k*-spaces to fit lines (*k* = 9, *d*_*i*_ = 1∀*i*); these 4-D clusters can be conveniently visualized as six 2-D scatter plots, corresponding to projections of the 4-D data and spaces onto 2-D planes. **c**, Each voxel in the qHCR image is then shaded with its *k*-spaces expression cluster index to reveal the anatomical distribution of each circuit state (e.g., red program, brown program, etc). 10 µm is a typical cell size. **d**, Visualization of spatial distribution in expression level for gene circuit states using pixel saturation (left; all programs) or average *L*_2_ norm of latent points versus *y* coordinate (right; red program). See Figs. S1 and S3 for additional data.

Reconstruction of anatomical features based on circuit state provides a powerful method to study the spatial usage of gene expression programs. Reconstruction also enables assessment of under-/over-clustering for different values of *k* based on visual inspection of expression space (Fig. S2) and anatomical space (Fig. S3). We varied *k* from 1 to 12 and determined that *k* = 9 was most appropriate for our data (Fig. S2a). Two of the nine clusters (blue and orange in Fig. 2c) correspond to voxels outside the embryo or voxels with a low basal expression level of the four mRNAs, leaving seven expression clusters to represent somitogenesis circuit states. Anatomical features emerge as *k* increases (Fig. S3a). The first feature identified was non-sample versus sample (*k* = 2). As *k* increases, the notochord, somites (S7 - S10), presomitic mesoderm (PSM), and adaxial cells are identifiable in the segmented image; eventually these anatomical structures become subdivided (Fig. S3a). Lines fitting multiple clusters (i.e., bimodality) in the scatter plots indicated underfitting, and overfitting was apparent in the image when voxels were partitioned on subcellular spatial scales (Figs. S2a, S3a).

Beyond segmenting the image into discrete circuit states, we use voxel saturation to depict the relative position of a voxel’s projection onto its assigned subspace (Fig. 2d), enabling visualization of co-expression changes produced by varying developmental programs in the embryo. Notably, this representation reveals that the red program (the mRNA stoichiometry in the expression cluster for which voxels are shaded red in Figs. 2b-d) is turning on with increasing strength as somites mature (increasing maturity S9→S8→S7; Fig. 2d).

Automated cluster identification and image segmentation of the whole image with *k*-spaces replicated and earlier extended findings (7) (Fig. 2c; see Fig. S4 for a side-by-side annotated comparison of segmented images). Replicated findings include: 1) usage of distinct gene expression programs in the maturing somites (S9, S8, S7; red program) versus presomitic mesoderm (PSM; pink program); 2) increasing expression of the red program as somites mature (Fig. 2d); 3) anterior-posterior regionalization of the nascent somite (S10), with the posterior region featuring the pink program found in the adjacent PSM and the anterior region featuring the red program found in adjacent maturing somites (S9, S8, S7); 4) corresponding anterior/posterior regionalization of the adaxial cells adjacent to the nascent somite (green program anterior, blue program posterior). New findings include: 5) additional segmentation of the adaxial cells between the nascent somite (S10) and the PSM (alternating between green and blue programs); 6) additional expression clusters corresponding to possible detection of distinct cell types within the somites (red versus brown) — discussed below; 7) a structure between the PSM and the left nascent somite (S10) that deviates from bilateral symmetry and features the same pink program featured in the PSM and the posterior region of S10.

Interestingly, *k*-spaces subdivides each somite, identifying distinct programs in the interior voxels versus an anterior and lateral border. Early in somite maturation, non-adaxial cells form transcriptionally-distinct anterior, medial, and posterior compartments, with the adaxial and posterior cells committing early to formation of slow muscle types (10–12) and medial fast fibers (11, 13), respectively, before migration within the somites as the somite undergoes rotation (12). Within the anterior compartment, there are anterior border cells (13), also called row one cells (12), which are located on the anterior and lateral edge of the somite (Fig. 1E of (13); Figs. 3 and 4 of (12)) and give rise late to lateral fast fibers (11, 13). In the maturing somites that we analyzed (S9, S8, S7), *tpm3* and *myod1* are expressed proportionally for both the red program (inner) and brown program (anterior and lateral), but with a distinctly lower expression level for *tpm3* in the brown program. Human *TPM3*.*12* is the muscle-specific isoform of *TPM3* found in sarcomeres, and studies have shown it is further specific to slow muscle in humans, mice, and zebrafish (14–17). Consistent with their slow muscle fates (18), the green adaxial cells express a substantially higher level of *tpm3*. The width of the brown region is approximately the expected ≈ 10 µm width of a cell in the embryo (approximated as the cube root of cell volume in Fig. S2 of (19)). Based on location and low *tpm3* expression, the brown region could specifically be anterior somitic compartment cells, but it may additionally reflect non myogenic lineages in the somite (20).

**Fig. 3.**
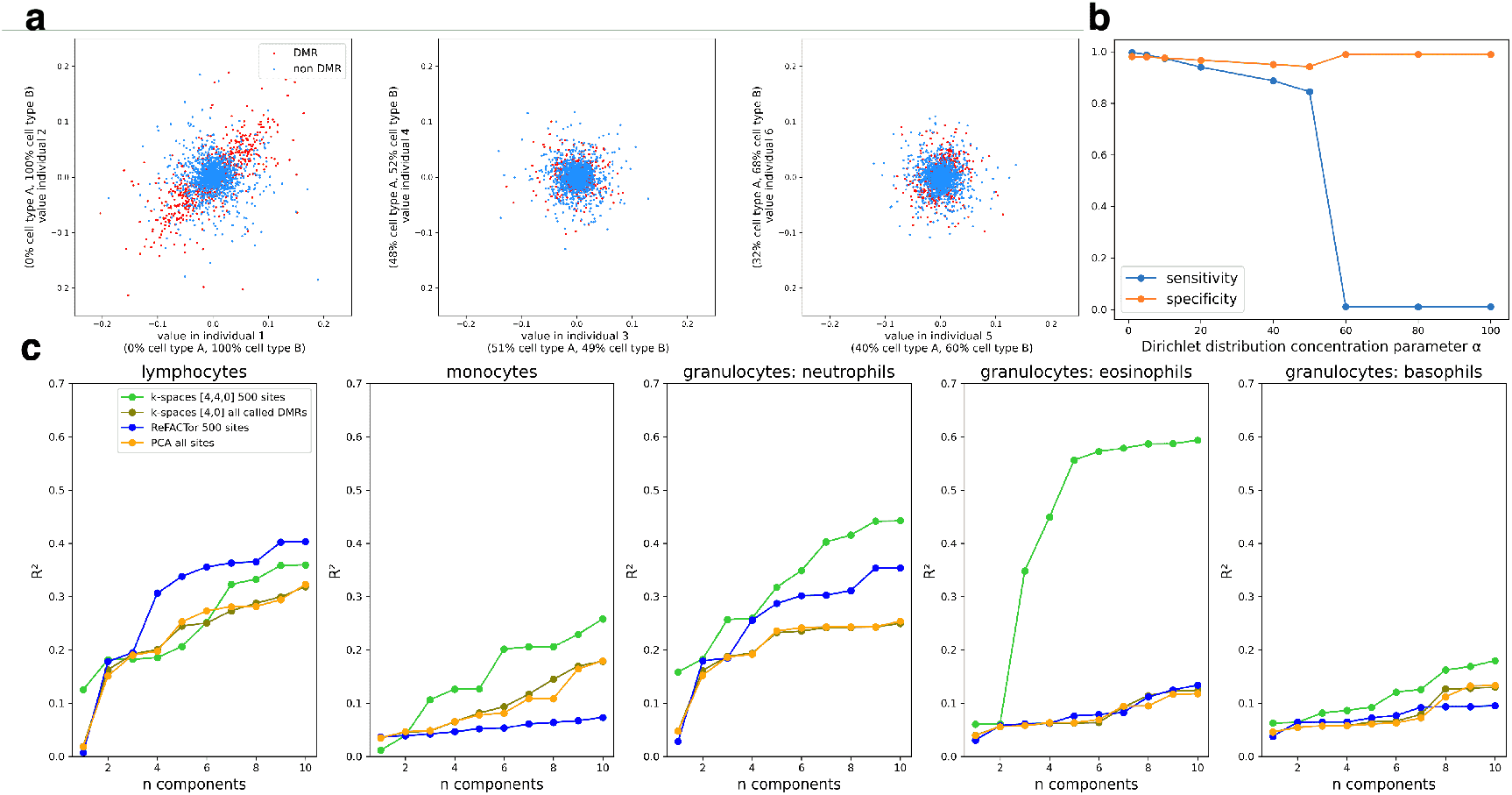
DMR detection in simulated and real data. **a**, Visualization of centered and transposed data of a toy example with only two cell types in six individuals. In the transposed data space, each axis is an individual while each point is a methylation site. DMRs are correlated between individuals with overlapping extreme cell-type proportions (left) while no cell-type proportion signal is observed in dimensions corresponding to individuals with prototypical cell-type proportions, defined as 50% cell type A and 50% cell type B in this simulation (center). In individuals with mild variation from the average proportions (right), a subtle signal is visible to distinguish DMRs from non DMRs, which is used by *k*-spaces to infer DMRs by examining sites across many individuals in high dimensional space **b**, Sensitivity and specificity of DMR detection in the simulation of five cell types using a symmetric Dirichlet distribution. When cell-type proportions varied by *>* ∼ 1-2%, DMRs were detectable, but when the variation in cell-type proportions becomes too low, the model failed catastrophically. **c**, Regression performance on cell-type proportions in the GALA II dataset.

**Fig. 4.**
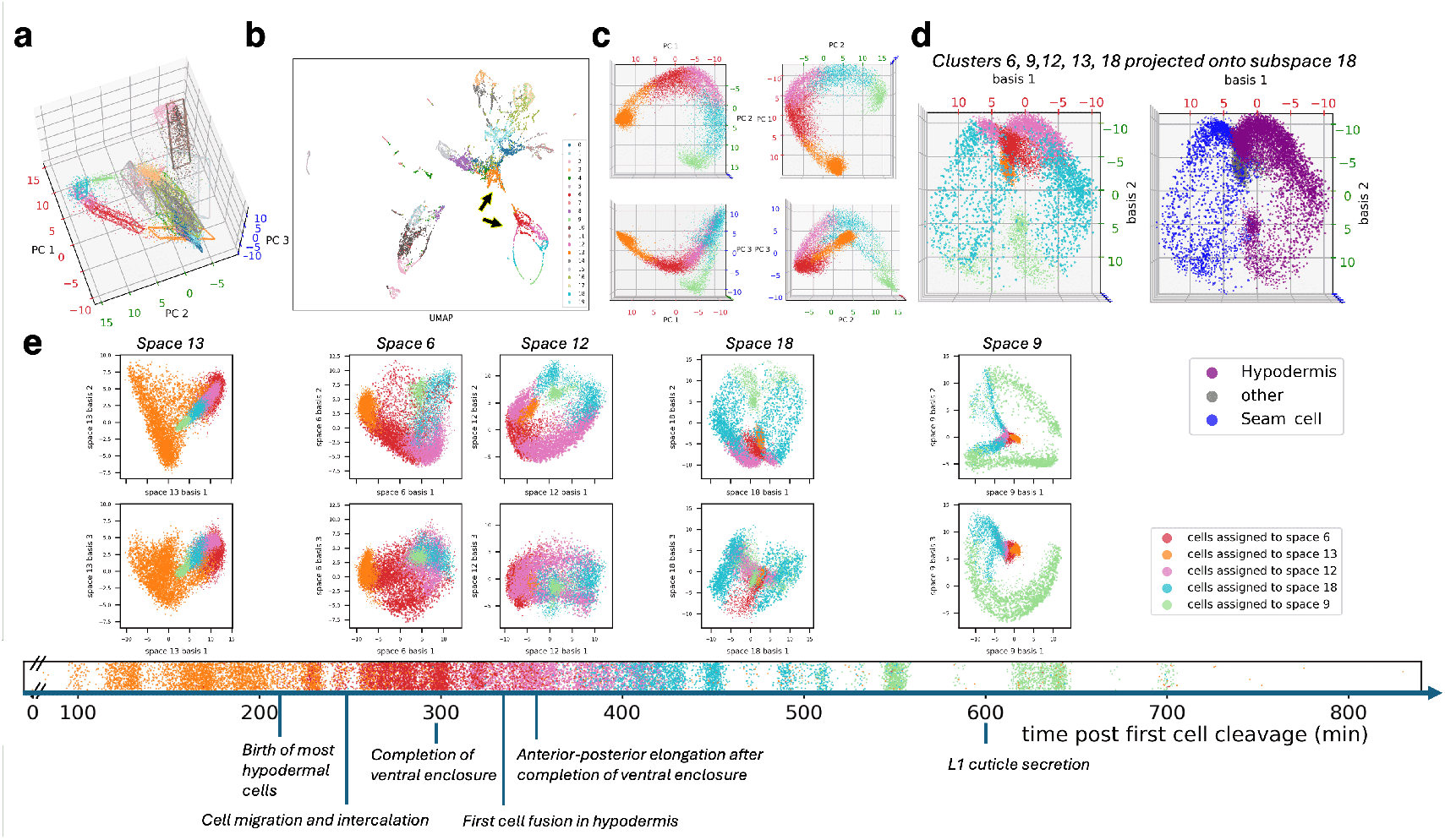
Projection onto multiple subspaces allows visualization of the hypodermis and seam cell trajectories. **a**, 3-D PCA visualization of developing cells with projections of *k*-spaces’s affine subspaces. **b**, UMAP visualization of the same cells from (a), colored by *k*-spaces cluster index. **c**, 3-D PCA of the subset of cells in the orange, red, pink, blue, and green clusters showing a different topology from the UMAP in (b). **d**, Projection of the same cells from (c) onto subspace 18, optimized for viewing gene expression changes in the blue cluster. Alternate coloring by cell type (right) shows that space 18 captures the bifurcation between the seam cell and hypodermis trajectories. **e**, Timeline of key events in epidermis and general embryonic development along with visualization of the cells from (c) and (d) using the dimension reduction yielded by each subspace. Cells are colored by *k*-spaces cluster index, but all cells are projected onto each subspace to visualize the full trajectory. The horizontal strip plot displays the time elapsed from first cell cleavage for cells from each cluster. Points are “jittered” by Gaussian random noise with a standard deviation of 3 minutes for visibility for illustrative purposes only. Sizes of points in (d) and (e) decrease with distance from the 3-D subspace they are projected onto. Thus, points with nearby projections but different sizes are separated in other dimensions of gene expression space. The “unjittered” timeline plot is shown in Fig. S21. Plots are reproduced with alternative coloring by cell type and embryo time in Figs. S16 and S17. Both cell types trace the same path over time in basis 2 vs 3 of space 9 (see Fig. S17 for coloring by time), suggesting parallel gene expression changes in both seam cell and hypodermis cell trajectories. There are also seam cell specific (space 9 basis 1 versus 2) and hypodermis cell specific (space 12 basis 1 versus 2) expression changes over time.

In summary, with *k*-spaces we use an interpretable framework to i) identify seven gene circuit states, ii) automatically segment the image into anatomical regions featuring those circuit states, iii) visualize oscillations of the developmental gene expression program, iv) observe predominantly symmetrical but also asymmetrical circuit states in the developing embryo, and v) potentially identify distinct cell types within maturing somites. This analysis suggests that *k*-spaces is able to recapitulate patterns found using other techniques but also capture novel patterns reflecting known biology about a system.

### *k*-spaces clustering can model DNA methylation probes in bulk epigenomics data driven by cell composition variation

Epigenome-wide association studies (EWAS) can be confounded by varying percentages of blood cell types in whole blood (21–23). This issue does not arise in classical genome-wide association studies, but cell types have unique DNA methylation profiles, and disease is often associated with increases or decreases in certain cell types (24). Reference-based methods use known methylation profiles of cell types to estimate cell-type proportions to control false positives (21). On the other hand, reference-free methods may use heuristics to identify a small number of sites in differentially methylated regions, and use PCA on those sites (hereafter referred to as differentially methylated regions, or “DMRs,” for notational consistency with prior work) as a proxy for cell-type composition (23).

A popular reference-free method, ReFACTor (23), uses heuristics to identify a preset number (500 by default) of differentially methylated sites in the GALA II dataset (25) and in simulations with a reference dataset (26) to capture information on the proportions of the underlying cell types. The study posed the following model: for each DMR, global cell-type means are sampled from a normal distribution, and samples’ underlying cell-type methylation profiles are sampled from another normal distribution with those cell-type means and a site-specific variance. These sample-specific underlying cell-type profiles combine with sample-specific underlying cell-type proportions to produce observed data. The study used extensive preprocessing of the GALA II data (see “GALA II analysis” in Methods, Fig. S8) prior to running the ReFACTor algorithm to identify the preset number of DMRs informative of cell-type composition. We set out to classify sites as DMRs or non-DMRs through probabilistic modeling and without preprocessing or heuristics.

We first demonstrate that *k*-spaces can solve the mathematical problem of classifying DMRs (23) using theory (see Supplement Section 2) and simulations. In our analyses, we centered the data so correlated deviations from global means may be modeled while retaining information about the variance. Simulated data was generated using cell-type proportions (25) and cell-type methylation profiles (26) from prior work. We ran *k*-spaces with a 4-D subspace (dimension one less than the number of cell types) and a 0-D subspace, respectively clustering and fitting sites with variation driven by cell-type composition (DMRs) or pure noise (non-DMRs). As subspace-specific noise parameters led to an unwanted partitioning of sites by high versus low site-specific variance (Fig. S9), and theory showed that a shared variance parameter between the spaces (see Supplement Section 3.4) may compensate for variation in the scale of noise in each methylation site, we exploited the constraint of sharing the variance parameter across the *k*-spaces.

When we sampled the site-specific standard deviations for noise from a half-normal distribution and bootstrapped the cell proportions from the GALA II data (*N* = 95), sensitivity was 90% and specificity was 94%. When we boot-strapped the variances from the reference data (26) used by the ReFACTor study (*N* = 6 individuals, *k* = 5 cell types, *M* = 485, 577 sites) sensitivity and specificity were 79% and 97%, respectively. In ideal conditions with high variance in cell-type proportions, near perfect performance was achieved with sensitivity *>* 99.5% and specificity *>* 98% (Fig. 3b). Using our probabilistic model, we attained not only a high positive predictive value, like ReFACTor did on a small preset number of sites, but we also produced a classifier with high sensitivity and specificity for DMRs in theory and simulation in the absence of batch effects.

Following prior work (23), we turned to the GALA II dataset to evaluate its suitability out-of-the-box for correcting confounding by cell composition variation in EWAS; here, the dataset does not contain purified cell type profiles from which to determine a ground truth for DMRs versus non-DMRs. In prior work (23), PCA of selected DMRs was correlated with cell-type composition measurements to support the claim that informative DMRs were identified by ReFACTor. However, in that study, both samples and sites were arbitrarily removed prior to testing the software, and we found that using the full dataset led to a failure of ReFACTor’s heuristic (Fig. 3c; Fig. S11a, b) and performance no better than PCA (Figs. S8, S11b).

We chose not to preprocess the data for *k*-spaces. An initial attempt at clustering with 4-D and 0-D spaces and a shared noise parameter predicted cell-type composition poorly in a downstream DMR-PCA versus cell composition correlation analysis (Fig. S11c). We hypothesized this may be because many of the called DMRs could be misclassified non-DMRs, or they could be *uninformative* DMRs because, while subtle effects of cell type composition variation may be detectable in a large sample, for any one individual, DMRs with low cell-type information may be dominated by technical noise, batch effects, or artifacts introduced by batch correction. Using the ground truth data from our prior simulation, we confirmed that classification of sites as DMRs or non-DMRs and selection of informative DMRs are not equivalent problems— PCA of all DMRs produced performance no better than using all sites for PCA (Fig. S12).

Because of the non-equivalency between the DMR classification problem and the cell proportion variation problem, we followed prior work (23) in selecting the 500 sites closest to the subspace for our analysis of the GALA II dataset. As we did not remove low-variance sites prior to clustering, we needed to remove sites close to the subspace solely by virtue of having low variance overall to avoid using DMRs that were more likely to be dominated by noise. To do this, we filtered out called DMRs using the estimated variance parameter of the 0-D space modeling sites with pure isotropic noise (Fig. S13). In addition, we incidentally found that not using the shared variance constraint led to better results on the full GALA II data (Fig. S11). We suspect that, unlike in the simulation, there were unaccounted latent factors (either biological or batch effects) in the GALA II data resulting in a subset of sites with “structured noise” rather than either isotropic noise or a signal reflecting cell-type composition variation. These latent factors were visually evident, showing that at least a portion of noise was likely non-isotropic through persistent streaks in the visual representation of the data (Fig. S13). The free variance parameter may have enabled the 0-D space to capture the “structured noise” sites with additional variance from latent factors along with the expected non-DMRs with isotropic noise.

To account for these latent effects, we ran *k*-spaces with an additional 4-D space to capture structured noise. Surprisingly, PCA of the sites captured by the 4-D space representing structured noise revealed distinct subpopulations in the data (Fig. S14) that did not correlate with sex or ancestry labels (Fig. S15). With the additional space to model structured noise in the full GALA II dataset and the post-hoc selection of informative DMRs, *k*-spaces achieved the high performance in predicting cell type composition variation seen with ReFACTor applied to the preprocessed data (Figs. 3c, S11). Comparisons with ReFACTor and simple PCA of all sites on the preprocessed data are shown in Fig. S11. While the unsupervised application of *k*-spaces to this type of data in practice is complicated by unexpected and unknown sources of variation, we found that *k*-spaces was suitable for the mathematical problem of manifold separation and DMR classification, and we showed it had the capability to model batch effects or unwanted variation.

### *k*-spaces offers subspaces for distance-preserving dimension reduction for exploratory data analysis in single-cell sequencing

Next, we used the dual functions of clustering and dimension reduction in *k*-spaces to resolve gene expression similarities and differences in the diverging developmental trajectories of two classes of epidermal cells—seam cells and hypodermis cells—in the *C. elegans* embryo. The embryogenesis atlas includes 86,024 single cells collected from bleach-synchronized embryos at three time points (estimated time post first cell cleavage was provided as metadata) of which 2,748 cells were seam cells and 7,093 were hypodermis cells (27).

Traditional linear dimension reduction methods such as PCA have been replaced with embedding methods such as t-SNE and UMAP for single-cell RNA sequencing data, based on claims that the complexity of structures in the data cannot be represented with linear maps (28–30). In turn, these nonlinear maps to two dimensions are known to produce artifactual structures and to heavily distort global distances (31). Here, we fit 20 3-D spaces to the 86,024 cells in the *C. elegans* embryo dataset to gain insight into relationships between “structures” in the UMAP visualization. We do this by coloring cells according to their *k*-spaces clustering (Fig. 4b) while also studying *k*-spaces’s informative and interpretable set of linear dimension reductions on these complex data (Fig. 4d, e).

After clustering, we examined the near-loop (indicated by lower black arrowhead) of the UMAP as the colors of several spaces were split across its arms, suggesting proximity between different sides of the loop, and additionally, the split orange cluster suggested proximity between the loop and the upper structure in the UMAP. PCA of those cells (the orange, red, pink, blue, and green clusters) revealed a broken loop as well (Fig. 4c), which was similar to that seen in the global PCA in Fig. 4a; however, the arrangement of colors along the structure revealed a different topology than was implied by UMAP. Therefore, we examined the clusters in each of their *k*-spaces subspaces, as each space is optimized for best representing the variance in its assigned cells in 3-D.

Unlike t-SNE and UMAP, which are not only nonlinear but also do not produce functions defined on the entire input space, the 3-D projections produced with *k*-spaces can be used to project any cell onto any space such as those from other clusters or from held out data. This allows us to observe the entire trajectory by projecting the cells across the set of linear subspaces (Fig. 4d) to reveal a bifurcation between hypodermis and seam cell trajectories, as suggested by the UMAP. In particular, the basis vectors of two of the subspaces - spaces 12 and 18 - represent directions in gene expression space along which seam cell and hypodermis cell developmental trajectories diverge, making these subspaces useful as dimension reductions to visualize and analyze the bifurcation event.

In addition, this application of *k*-spaces also revealed both lineage-specific changes (space 9 and space 12) and parallel gene expression changes shared by the two lineages (space 9) that cannot be inferred from UMAP as UMAP axes do not correspond to gene expression. For the shared changes as well as the axis of bifurcation (space 18 basis 1), we further observed that more mature cells returned to the same state as early progenitors with respect to these basis vectors. We also observed the divergence of the seam cells and hypodermis cells from developing neurons (space 13), properly connecting the ectoderm lineages. These results together suggest that *k*-spaces yields more interpretable linear projections and clustering of complex, high-dimensional data than the 2-D nonlinear alternatives.

Although *k*-spaces was only applied to gene expression data, cluster membership aligned with intervals of embryo-time reported in the atlas (Fig. 4e). Thus, each subspace was optimized for coordinated gene expression changes during a different developmental period. A second benefit of plot axes corresponding to meaningful directions in gene expression space is the ability to extract the genes contributing most to each basis vector. We compared the genes best capturing each subspace to ground truth knowledge of embryonic events (Fig. S18, Tables S20, S19). For example, we interpreted two basis vectors in one space (1 and 3 in space 9) as potential shared gene expression programs for cuticle synthesis, which both cell types participate in. Examining bases of other spaces revealed several signaling genes, such as warthog and groundhog-like gene family members, genes involved in molting, and dumpy family members (named for mutants producing a shortened worm), among others. Collectively, these results suggest that *k*-spaces structures recapitulate known biological signals more accurately from unsupervised data analyses than their nonlinear counterparts.

## Discussion

In this study, we presented a probabilistic model that we call *k*-spaces for joint clustering and dimension reduction of data generated by mixtures of linear subspaces defined by latent Gaussian variables. We described how the *k*-spaces model relates to other methods in this space, and showed its applications in biological data analyses. One of the primary motivations for this study was to develop a framework to study the latent space of gene expression and how correlated modules of genes scale their expression across time or space in response to external stimuli or innate physiological demand. Our *k*-spaces framework may generally be used to study systems that may be approximated as linear latent variables modulated by categorical latent variables such as cell type, tissue type, batch, sex, disease status, or genetic background. Though inspired and initially derived from a *k*-means framework (32–37), we have connected *k*-spaces to PCA, GMMs, and probabilistic PCA, reiterating closely related results and interpretations in prior work (2, 6), while also generalizing the model to subspaces of mixed dimension and shared noise parameters.

In retrospect, it is unsurprising that this more general model is connected with these specific models, as the *L*_2_ norm and the Gaussian distribution are fundamentally linked (38, 39). Methods that find the mean of data (such as total least squares (40, 41) or *k*-means clustering), which corresponds to minimizing the *L*_2_ norm or sum of squared residuals, implicitly fit Gaussian distributions, and PCA has Gaussian interpretations both from the perspective of minimizing the *L*_2_ norm and maximizing the likelihood of Gaussian data (see Supplement Section 1). Thus, we can approach this *k*-spaces framework both as an interpretable clustering and dimension reduction method or as a linear latent space approximation where a theoretical analysis of failure modes is tractable. We term our method *k*-spaces to emphasize the focus on probabilistic learning of subspaces rather than the specific distribution, and future work can include generalizations to other data-relevant distributions (42, 43).

Our application to qHCR imaging of zebrafish somitogenesis shows the potential for *k*-spaces to be used to scale and automate current methods of analyzing biological data that are done by hand, especially where the learned subspaces are themselves valuable for study and can be assessed with human expertise but manual clustering is intractable as direct visualization of the complete data is not possible.

Given the exploratory power of quantitative analyses moving back and forth between anatomical space and expression space, it is of interest to increase the scale of readout/read-in analyses (more genes, more ROIs, more expression clusters). However, as the problem scale increases, a manual approach becomes unwieldy, both because the number of 2-D scatter plots increases quadratically with the number of multiplexed targets (e.g., 10 targets yields 45 2-D scatter plots), and because the number of regions of interest (in anatomical space) and expression clusters of interest (in expression space) can grow out of hand as the spatial extent of the study increases, as in whole-embryo analyses. qHCR spectral imaging now enables robust 10-plex quantitative imaging in highly autofluorescent samples including whole-mounts and delicate specimens that are not amenable to repeated imaging (9), opening the door for 10-dimensional automated read-out/read-in using *k*-spaces on whole-mount embryos to examine more and larger circuits. qHCR imaging has also been extended to enable compatible quantitative imaging of RNA (7, 8), protein (44), and protein:protein complex targets (45) in highly autofluorescent samples, creating the opportunity for automated read-out/read-in using *k*-spaces to examine circuits involving selected targets from the transcriptome, proteome, and interactome.

Application of *k*-spaces to qHCR data revealed another benefit in terms of its inductive bias. Bishop and Tipping demonstrated the usefulness of pPCA over full rank Gaussian models in the setting of high dimensional, sparse data, where fewer free parameters makes pPCA more robust to overfitting (1). Here, the inductive bias of *k*-spaces—that gene expression is scaled along lines—allows it to identify clusters that would be merged by full rank GMMs, and indeed, cluster labeling of voxels appears to be more anatomically correct with *k*-spaces than with GMMs or *k*-means (Figs. S3 and S7). Furthermore, full rank GMMs lack the notion of a latent space, while *k*-means has a 0-D latent space and no continuous latent variable, but only discrete clusters and noise (though we should properly refer to this as a spherical GMM with equal covariance matrices in a hard assignment setting). In contrast, *k*-spaces has an explicit notion of a latent space and allows us to visualize oscillations of gene expression programs in complex systems such as a developing embryo. *k*-spaces could also be applied to qHCR images of other systems such as bacterial biofilms (46) or to spatial RNA sequencing data, though application to unnormalized sequencing data would likely require an extension of the *k*-spaces software using a probabilistic model for count data (43).

*k*-spaces also strikes a balance between being sophisticated enough to model complex data while being simple enough to theoretically analyze. In the context of bulk epigenomics, we analytically showed that the *k*-spaces model is appropriate for this data (even when the variation in noise across methylation sites produces a non-Gaussian distribution of noise) by (mis)using a shared noise parameter between the spaces. Furthermore, application to epigenomics allowed us to identify a sudden failure of the model at *α* between 40 and 50 (Fig. 3c). Our prior theoretical analysis suggested a possible cause, and we support this with simulation evidence by removing the top 0.03% noisiest sites to restore robustness to low cell proportion variation (Fig. S26; see Supplement Section 3.4).

The inspiration for applying *k*-spaces to bulk epigenomics data, ReFACTor, was tested on a subset of the GALA II data, both in terms of samples and sites, and used heuristics (23). While we showed that the DMR classification problem as set up by the simulation in (23) could be solved by *k*-spaces, we learned that selection of informative DMRs, whether in real or simulated data, was a distinct problem (Fig. S12). Theoretical justification or empirical validation of our heuristic selection of “informative” DMRs after clustering would be needed for wider use. Furthermore, while visualization of the data (Fig. S14) suggested which of the two 4-D spaces represented the desired signal and which was structured noise, a robust selection strategy of the correct dimension for the second space is unclear; we merely chose 4 as it seemed plausible that a subset of sites were behaving systematically differently yet also affected by underlying cell proportion variation. In the future, the out-of-the-box *k*-spaces framework could be tailored to this epigenomics problem. However, with the wider availability of single-cell epigenomics assays (47), development of a robust reference-free computational method to identify DMRs may be less of a pressing need and is outside of the scope this study, though we used this example to challenge *k*-spaces with a problem requiring manifold separation through dimension.

We also explored other applications in transcriptomics where *k*-spaces may be used. In addition to visualizations for single-cell data (Fig. 4), we also examined species mixture “barnyard” plots and gene correlation in single-cell sequencing. In droplet-based RNA-sequencing, there is some low level of background contamination in the sample. *k*-spaces offers a principled way to select the threshold for when nonzero UMI counts are too high and the droplet likely captured more than one cell (See Supplement Section 6, Figs. S22a, S23).

We next assessed whether distinct relationships between pairs or groups of genes can be identified and characterized with *k*-spaces clustering in single-cell data, analogous to our qHCR analysis (see Figs. S22, S24, S25). This is important because experiments that identify co-regulated modules of genes using techniques, such as weighted gene co-expression network analysis (48), often begin by constructing a gene-gene correlation matrix, but this assumes the linear correlation between a pair of genes is representative of their relationship.

While *k*-spaces was able to detect distinct relationships, it overclustered the data when used in an unsupervised fashion with model selection via the Bayesian information criterion (BIC) or the integrated completed likelihood (ICL), which adds an entropy penalty to BIC to approximate the joint likelihood of the data and the clustering solution (49, 50). One particularly challenging artifact contributing to this overclustering is a linear structure in pairwise scatter plots – especially in genes with low counts – introduced by depth normalization because angular position is determined by pairs of discrete counts, producing streaks in the data with quantized slopes but pseudo-continuous scaling along the lines. In the extreme case, *k*-spaces with Gaussian distributions will fit a separate noiseless line to each streak, and even outside of this extreme setting, it is difficult to determine whether the “distinct” spaces are artifacts rather than biology. To address this and other challenges, we envision an extension of our *k*-spaces framework with a generalization of the negative binomial distribution to mixture models, where the multidimensional gamma priors in the gamma-Poisson formulation capture the latent spaces to be inferred. We anticipate this will be particularly useful in handling the artifactual linear structures as well as overdispersion. In genes with low counts, it is not necessarily true that *no* shared latent variable controls gene expression and that all such gene abundances are independent; rather, we lack the resolution to make any inference with *certainty*. Modeling this within a Bayesian framework should make this evident with wide, uncertain posteriors.

Finally, we believe one of the most important contributions of *k*-spaces is the expansion of the biologist’s exploratory data analysis (EDA) toolkit to provide insights in to global relationships and better visualize finer structures with *k*-spaces. Most applications discussed in this study involve directly modeling latent biological processes, but we show the ability to tease apart these finer structures with the single-cell RNA-sequencing data for *C. elegans* embryogenesis. Importantly, this neither requires proper selection of *k* nor finding the “correct” clustering solution, as underclustering to capture multiple structures in one subspace allows their comparison, and visualization with different *k*-spaces results can yield different insights. Bishop and Tipping used a mixture of pPCAs to partition a dataset and view the relative maximum a posteriori positions of data points within their own assigned pPCAs (6), and they used a hierarchical technique that performed mutual projection to only show ambiguously assigned points (51). We extend this idea by projecting points onto each other’s subspaces to view entire structures in the context of gene expression space. The hierarchical technique for pPCA mixtures proposed by Su and Dy (5) could also be useful in guiding EDA, though the nonhierarchical clustering we use may be more helpful for detecting global similarities in distantly related cells.

One limitation of our approach is that *k*-spaces visualizations can plot points in proximity to each other that are well-separated in another axis of variation. This limitation echoes the same behavior in PCA in obscuring fine structures when the data are underclustered. We addressed the former by decreasing the sizes of points to reflect increasing distance from the 3-D space of projection. Thus, larger points that are plotted in proximity are likely to be close in the ambient gene expression space, points of drastically different sizes are guaranteed to be separated in unseen dimensions, and small points are understood to be far away but may diverge from the subspace in different directions. The Eckart-Young theorem (52) states that PCA is the optimal linear dimension reduction for visualizing variance in data. By definition of being nonlinear, other methods will distort distances or, worse, provide embeddings with features that do not correspond to continuous axes – linear or nonlinear – in gene expression space. Visualizing the data with piecewise linear dimensionality reductions trades off the global optimality of PCA for local optimality to offer more fine grained visualizations of structure while still providing interpretable axes. Critically, the shortcomings of *k*-spaces are tractable to analyze for a specific application. In this particular case each space can be viewed as a 3-D PCA for a subset of the data. By using this framework to visualize single-cell data, we can not only see high-resolution structures in data but also quantify the reliability of our inferences.

## Data and code availability

The *k*-spaces software package is available at https://github.com/pachterlab/k-spaces.

The purified human blood cell reference epigenomics data was accessed on September 11, 2024 at https://www.ncbi.nlm.nih.gov/geo/query/acc.cgi?acc=GSE35069. The GALA II epigenomics data was accessed on September 17, 2024 at https://www.ncbi.nlm.nih.gov/geo/query/acc.cgi?acc=GSE77716. Code to preprocess data is included in our notebooks in our repository but sample and probe IDs for the full and the filtered datasets are additionally available at https://github.com/pachterlab/MEPSP_2024/data_paper.

HCR data was downloaded as part of the Readout/Read-in Software Package on November 15, 2024 from https://www.moleculartechnologies.org/info/software. Both the raw data and preprocessed matrices are available in our repository at https://github.com/pachterlab/MEPSP_2024/data_paper.

The *C. elegans* embryogenesis data was accessed on August 25, 2025 at https://www.ncbi.nlm.nih.gov/geo/query/acc.cgi?acc=GSE126954.

The human-mouse mixture, 10K PBMC, and 5K PMBC datasets were downloaded from the 10X Genomics website at https://www.10xgenomics.com/datasets/10k-hgmm-3p-gemx, https://www.10xgenomics.com/datasets/10k-human-pbmcs-3-v3-1-chromium-x-without-introns-3-1-high, and https://www.10xgenomics.com/datasets/5k_Human_Donor4_PBMC_3p_gem-x (filtered matrix), respectively.

We used anndata v0.9.2, MATLAB v24.1.0.2653294 (R2024a) Update 5, Numpy v1.23.5, Pandas v2.1.4, Python v3.10.9, ReadoutReadin v1.0, ReFACTor v1.0, v1.0scanpy v1.10.2, scipy v1.11.4, and sklearn v1.3.0. The full environment is available at https://github.com/pachterlab/MEPSP_2024. Code for reproducing the results and figures in the manuscript is available at https://github.com/pachterlab/MEPSP_2024 under Notebooks.

## Author contributions

The *k*-spaces clustering algorithm was conceived by BE and LP, and developed and implemented by NM. NM, BE, NP, PS, and LP conceived the applications, and NM generated the results. NP provided qHCR data and directed its analysis. NM, BE, NP, PS and LP analyzed results. The manuscript was drafted by NM, and was edited and approved by all authors. PS and LP supervised the project.

## Acknowledgments

We thank the Banff International Research Station for supporting and hosting the workshop on challenges and synergies in the analysis of large-scale population based biomedical data in Oaxaca, Mexico in 2017. The workshop led to discussions between BE and LP that led to this project. We thank Caleb Ghione, Ulrich Herget, and Peter Currie for their kind assistance in interpreting HCR data and discussions about zebrafish embryology and anatomy. This work was supported by NIH F30 1F30GM156092-01 to NM and by NIH-NHGRI grant U24HG010859 to PWS, a Bren Professor of Biology, and by the Beckman Institute at Caltech (PMTC). BEE was funded in part by grants from the Parker Institute for Cancer Immunology (PICI), the Chan-Zuckerberg Institute (CZI), the Biswas Family Foundation, NIH NHGRI R01 HG012967, and NIH NHGRI R01 HG013736. BEE is a CI-FAR Fellow in the Multiscale Human Program.

## Methods

### Overview

Our model consists of a mixture of latent Gaussian variables defining distributions on *k* affine subspaces *s*_1_ … *s*_*k*_ of dimensions *d*_1_ … *d*_*k*_ with the latent points perturbed by isotropic Gaussian noise in the complementary space to generate observed data. We consider the cases where the variance for noise can vary by latent space and where it is the same for all subspaces. For conciseness, we refer them to the individual latent spaces as “spaces,” and we refer to our mixture model as *k*-spaces.

We model each observation **x** ∈ ℝ^*D*^ as the sum of a structured component in a latent subspace and an independent noise component in the complementary subspace such that:

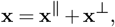

where **x**^∥^ ∈ ℝ^*d*^ lies in a latent affine subspace associated with a mixture component (signal), and **x**^⊥^ ∈ ℝ^*D*−*d*^ lies in the orthogonal complementary space (noise). Under this model, the density of **x** conditioned on mixture component space *s* factorizes as

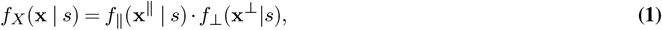

where *f*_∥_ (**x**^∥^ | *s*) is the multivariate Gaussian density of the latent component under the mixture model, and *f*_⊥_ (**x**^⊥^| *s*) is the isotropic Gaussian density for the independent noise component of **x** that lies in the complementary space.

We denote by *f*_*X*_ (**x**_*j*_; *s*_*i*_) the probability density function of observation **x**_*j*_ under space *s*_*i*_. Let *n*_*i*_ = *D* − *d*_*i*_ for convenience. The probability density function for each space is therefore

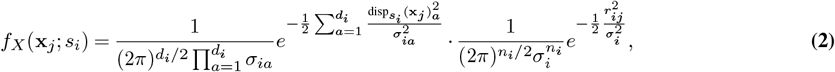

where 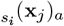 and 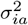 correspond to the displacement of the projection along principal axis *a* of subspace *i* and that axis’s respective variance, 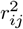 refers to the length of the orthogonal residual from **x**_*j*_ to *s*_*i*_, and 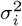 refers to the variance of the noise in each complementary dimension for space *i*. Note that the resulting distribution is Gaussian and the density function can be written concisely with a single covariance matrix. However, this form, which emphasizes the spectral decomposition, allows for separation into the first term corresponding to the latent space and the second corresponding to the complementary space. It is of interest in biological applications where the position relative to other latent points or to the mean within the latent subspace rather than position in the ambient space (or conversely the distance from the subspace) is relevant.

Each component has a mixing proportion *π*_*i*_, where 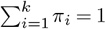. The marginal likelihood of **x**_*j*_ under the mixture model is then given by

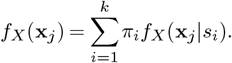

Unless noted otherwise, throughout the Methods and Supplement our derivations relate to this probabilistic model. We estimate the parameters by maximizing the model likelihood with an expectation-maximization (EM) approach, in which points are soft-assigned to spaces based on their probability density functions and mixing proportions, and then spaces are fit in parallel based on eigenvalue decomposition of their points’ weighted covariance matrices. The noise is determined as a function of the residuals (see Supplement Section 2). In the case of hard-assignment, where each point is fully assigned to its most likely space of origin, the M-step is performed with a singular value decomposition (SVD) solver. Lastly, also implemented in our software package is a direct generalization for *k*-means, where points are assigned to the closest space and spaces are fit to their assigned points by SVD. This is distinct from our probabilistic model as it assumes no distribution over the latent space and is equivalent to assuming the noise is equal for all spaces 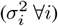, i.e., this approach directly minimizes reconstruction error and is not a generative model.

### Derivation

We can generalize the objective function for *k*-means clustering,

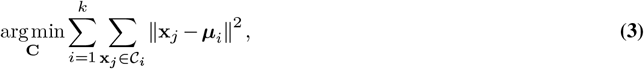

to an objective for clustering onto a set of *k* affine subspaces *s*_1_ … *s*_*k*_ of dimensions *d*_1_ … *d*_*k*_:

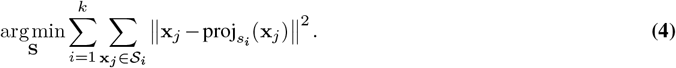

In *k*-means, *C*_*i*_ is the set of points closer to the mean of cluster *i* than any other cluster. In the most immediate generalization, points can be assigned to their nearest space; this was done in the image compression and computer vision communities (32, 33, 35). However, we assume a latent distribution and therefore for hard assignment we use **x**_*j*_ ∈ *S*_*i*_ if *i* = argmax *f*_*X*_ (*x*_*j*_; *s*_*i*_)*π*_*i*_. Relaxing the hard-assignment requirement, we obtain a maximization of the expected complete log likelihood

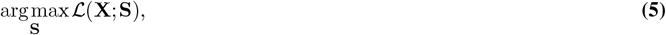

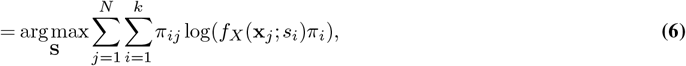

which we optimize by EM. Setting *d*_*i*_ = 0 and *σ*_*i*_ = *σ* ∀*i* yields *k*-means clustering in the hard-assignment setting or its probabilistic analog: spherical Gaussians of equal variance. Setting *k* = 1 yields PCA. In the M step of the EM procedure for this algorithm, PCA can be performed to fit spaces to their assigned points, as the optimal decomposition into **x**^∥^ and **x**^⊥^ minimizes the distances from points, **x**, to their projections, **x**^∥^, as in equation (4), making this a PCA mixture model (see Supplement Section 1.1). In many contexts, it is desirable to perform a probabilistic assignment of points to spaces as well as to assume a probability distribution over the latent space, which is critical to proper clustering when the dimensions of the subspaces varies, as will be discussed shortly. Without a latent distribution, higher dimensional spaces are favored over lower dimensional ones for clustering. When we assume latent Gaussian distributions with isotropic Gaussian noise in the complementary space, we obtain a constrained Gaussian Mixture Model (GMM), which is approximately equivalent to a mixture of pPCAs with varying dimension.

#### Model selection reveals the need for a latent distribution

An approach based on likelihoods of residuals alone fails when comparing spaces of different dimension, as subspaces of higher dimension observe out-of-subspace variance in fewer dimensions of the data. Consider a sufficiently large set of points sampled from a bivariate standard normal *N*_2_(**0, I**). Our objective in equation (4) minimizes out-of-subspace variance, the per-point mean squared distance to the subspace. Fitting a single point (*k* = 1, *d* = 0) will yield a space with basis = ∅ and translation = **0**. Because the bivariate standard Normal is rotationally symmetric, any line that passes through **0** is an equally good fit, so without loss of generality, assume fitting a single line (*k* = 1, *d* = 1) yields the basis [1,0] and translation **0**. Thus, it is apparent that the mean squared residual length 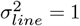 as the residuals’ lengths correspond to the y-values of the points in the sample. However, for the point,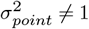. The mean squared residual is 2 — the distances from points to the origin are 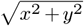, and each dimension has a variance of 1. Note that the distances do not follow a Half-normal distribution; they follow a Rayleigh distribution, but regardless, the likelihood of residuals when fitting a point must account for variance in both the *x* and *y* dimensions, while the likelihood of residuals when fitting a line only needs to account for variance in the *y* dimension. Although the data was generated from a 0-dimensional subspace with isotropic noise, if we were choosing between models with *d* = 1 and *d* = 0, the likelihood of the residuals under the *d* = 1 model would be better, and if we had a model with both spaces and were assigning points in the expectation step, the *d* = 1 space would always be favored except exactly at the origin. For these reason, we assume a Gaussian distribution over the latent space, akin to Bishop and Tipping’s pPCA.

### qHCR Image Analysis

#### Preprocessing

Data for the analysis in Fig. 2 is included in the Read-out/Read-in v1.0 software package, which was retrieved from www.moleculartechnologies.org/supp/ReadoutReadin1.0.zip. A single 304 x 510 pixel ROI was placed at the x, y coordinates (1,3) in the image, and the voxel size was 3×3×1 pixels as in the default setting. Voxel intensities are normalized per-channel by default in the Read-out/Read-in workflow. Rather than perform scatterplot analysis in the user interface, the data was exported and used in our *k*-spaces analysis.

#### k-spaces clustering

*k*-spaces was fit with *d*_*i*_ = 1 ∀*i* for *k* ranging from 1 to 12, with 100 random initializations each and a tolerance setting of tol =.0005. This was repeated with *d*_*i*_ = 3 ∀*i* (a standard GMM in 4-D space) and with *d*_*i*_ = 0 ∀*i* and assignment = ‘closest’ (standard *k*-means) rather than the default of ‘soft’, as in all other portions of this study. For coloring the voxels by cluster in the figures as well as coloring points on the scatterplots, final soft cluster assignment was converted to a hard assignment by selecting the space with the maximum responsibility for each point.

#### Scatter plots

In the plots, line segments (the basis vector for each 1-D affine subspace) were centered at the cluster mean and extended for one standard deviation in each direction along the latent space before projection onto the plane of the plot. Line lengths therefore reflect absolute variation in expression for each cluster and each gene, and relative lengths for the same cluster across different plots reflect relative influence of each gene on the direction of the space.

#### Visualization of spatial gradients in Fig. 2d

In the plot of *y* coordinate vs gene circuit expression, for each cluster, the mean *L*_2_ norm per *y* coordinate 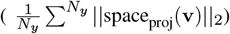 was plotted if there were at least 5 voxels with that label at that *y* coordinate. 5 voxels was chosen because it corresponds to 10 *µm* (the approximate cell dimension). In the image, pixel intensity was determined by 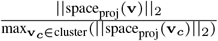, where voxels of each cluster were normalized to a maximum of 1 separately. Without this normalization, clusters with overall low gene expression would otherwise appear dim, and both hue and gradients would be difficult to discern. An alternative would be to use the displacement along the cluster latent space (the line) from the cluster mean and to scale values from 0 to 1 for saturation. However, this would result in a saturation of 0 for some voxels despite the presence of nonzero gene expression and may be a misleading visualization.

### Differential Methylation

#### Simulation

Simulation was performed as described in the ReFACTor study (23). Let 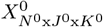 be a matrix of observed methylation values for each individual at each site and for each cell type,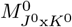 cell type at each site, averaged across the 6 individuals, and 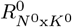 be a matrix of methylation values for each be a matrix of each individual’s cell type proportions in the reference data (*N* ^0^ = 6, GSE35069) (26). Notation here is chosen for consistency with (23) and does not refer to variables of the same name in our presentation of the *k*-spaces model. The simulation procedure used is as follows:

1. Determine a unique variance for each site, 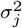: An empirical distribution of site-specific variances was determined via 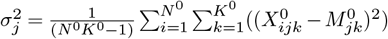. However, *σ*_*j*_ ∼ Half-normal(0, *σ*^∗^ = 0.03) was used in Fig. S26 and prior to bootstrapping variances from the above empirical distribution for the final simulation as noted in the text.
2. For each site, sample *µ*_*j*_ from some distribution: This step is not particularly relevant as we center the data, but we used Norm(0.5, 0.15).
3A. Randomly select 15% of sites to be DMRs: For each of these sites, sample *K* values for the row *M*_*j*_ via *M*_*jk*_∼ Norm(*µ*_*j*_, *τ* ^2^ = 0.07). *τ* was determined in the ReFACTor study.
3B. For non DMR sites, set *M*_*jk*_ = *µ*_*j*_ for all *k*.
4. For each individual and each cell type, at each site *X*_*ijk*_ ∼ Norm(*M*_*jk*_, *σ*_*j*_).
5. Sample each individual’s cell type proportions **R**_*i*_ from a Dirichlet distribution or bootstrap from reference data.
6. Obtain each *O*_*ij*_ by taking 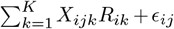 where *ϵ*_*ij*_ ∼ Norm(0, 0.01) and truncate values to [0,1].

500 individuals and 100,000 sites were simulated with 5 cell types. *k*-spaces with a 4-D and 0-D spaces was fit to the data using a shared noise parameter. Sites were called as DMRs if their assignment responsibility to the 4-D space was greater than 0.5.

#### GALA II analysis

For the analyses in supplementary Figs. S11, S27, and S28, filtering of samples and sites prior to using the ReFACTor software was done as described in the ReFACTor study (23). Normalized and batch-corrected data in the series GSE77716 were downloaded from GEO. First, samples were projected onto the top two PCs. Any sample greater than two standard deviations away from the mean on either PC was removed. Then, all sites with means greater than 0.8 or less than 0.2 were removed. Prior to running ReFACTor, we did not manually remove sites with standard deviations less than 0.02, nor did we center and scale the data to unit variance, as these steps are part of the ReFACTor algorithm and are done by its software. ReFACTor v1.0 was run with k = 6 as described in its paper with default settings, t = 500 and standard deviation threshold = 0.02. For the *k*-spaces clustering in the analyses in Figs. 3 and S13, the data from GEO was used directly and only centered. In neither case did we scale the data, and all samples and sites were included. Clustering was performed with two 4-D subspaces (to model signal and structured noise) and a 0-D subspace (to model unstructured noise) with a shared noise parameter unless noted otherwise. Ancestry labels used in Fig. S15 were taken from the “ethnicity” column of the GALA II metadata.

#### Preprocessed subsets of GALA II data for Fig. S11

The ReFACTor software itself removes low-variance sites; the numbers below reflect the dataset sizes that were inputted to *k*-spaces and ReFACTor. Note that the numbers in A will not match (23) exactly as only 525 samples were available at the time of their study. For example, the number of individuals with cell proportion measurements after preprocessing in (A) was 80, while in (23) there were 78. Similarly, there were 102,503 probes used in their analysis to our 102,470 in (A).

A. the data produced by the preprocessing procedure described in (23) (subset of 546 individuals and 102,470 sites).ReFACTor removed 1,142 low-variance sites downstream.
B. filtering out sites on the basis of mean *>* 0.8 or *<* 0.2 (all 573 individuals, subset of 103,024 sites). ReFACTor removed 640 low-variance sites downstream.
C. filtering invdividuals on basis of PCs (subset of 546 individuals, all 473878 sites). ReFACTor removed 263,620 low-variance sites downstream.

Full: all data (all 573 individuals and 473,838 sites). ReFACTor removed 226,847 low-variance sites downstream.

### Transcriptomics

#### C. elegans embryogenesis

A preprocessed single-cell RNA sequencing atlas of C. elegans embryogenesis was from GEO (GSE 126954), comprised of 86,024 cells from gastrulation to terminal cell differentiation that passed quality control (27). We multiplied counts by the size factors determined by Packer and coworkers to depth normalize and log1p transformed the data. We then removed 1,172 genes with 0 variance and used SVD for a linear dimensionality reduction to 100 dimensions, a typical step performed before nonlinear dimensionality reduction methods like t-SNE (28) and UMAP (29). We then separately produced a 2-D UMAP visualization of the data with default parameters and ran *k*-spaces clustering with 20 3-D spaces to obtain a piecewise set of linear dimensionality reductions for the data. *k* = 20 was chosen simply because the tab20 color palette has 20 colors, and the goal was not necessarily to produce an anatomically- or developmentally-coherent clustering to match a ground truth but rather to partition the data into a manageable number “viewing windows.” Timing for annotated embryogenesis events in the timeline of Fig. 4 were obtained from WormAtlas (53, 54).

After choosing spaces 6, 9, 12,13, and 18 due to the discrepancy in their clustering with the visible near-loop in the UMAP, we projected cells assigned to all 5 spaces onto each of the 5 spaces to confirm a continuous trajectory and view gene expression changes represented by each space. As projections can obscure separation between points in other dimensions and each space maximizes variance within only its own cells and is not necessarily a good dimensionality reduction to show separation of cells assigned to other spaces, we encoded the distance for each cell from each subspace by the size of points. We scaled the size of the points by the cube of their distance in the ambient space from the subspace. We chose to raise them to the power of 3 because it provided a better dynamic range.

After selecting basis vectors (100-dimensional) that appeared to represent key parts of the trajectory, we mapped them back to gene expression space (19,050-dimensional) by multiplication with the original SVD components. We retrieved a list of genes to represent each PC based on a cutoff of +/-0.05 for loadings and performed complete gene ontology enrichment analysis for biological process, molecular function, and cellular component with PANTHER release 19.0 (overrepresentation test/Fisher’s exact test with FDR multiple testing correction) (55, 56). The cutoff of 0.05 was chosen based on the empirical cumulative distribution functions of the basis loadings (coefficients denoting the direction of the basis vector in gene-expression space).

#### Barnyard Plot

The Cell-ranger preprocessed 10k 1:1 Mixture of Human HEK293T and Mouse NIH3T3 Cells, Chromium GEM-X Single Cell 3’ dataset was downloaded from the 10X Genomics website. The number of UMIs mapping to the GRCh38 and GRCm39 genome assemblies were summed for each cell. No further preprocessing was performed.

#### Gene-gene relationships

The 10K PBMC 3’ nextgem Chromium X filtered feature by cell matrix and 5K Human PBMCs (Donor 4) were downloaded from the 10X Genomics website and preprocessed with scanpy (57). Cells with counts in less than 100 genes and less than 3000 total counts were removed, as well as genes expressed in less than 200 cells. Doublets were then removed with scrublet (58). Counts were then depth normalized to the median number of counts and log1p (natural logarithm as per the default in scanpy.pp.log1p) transformed.

After this, further filtering of cells and genes were performed to avoid clustering on genes with apparent coexpression due to normalization artifacts such as those in perceptible SRSF5 vs ATP5MG in Fig. S24, as structure in the data for genes with low counts is dominated by artifacts of depth normalization for count data. This effect is much more pronounced in non highly expressed genes. To produce random examples of clustering in Fig. S24 with the 10K PBMC dataset, genes with less than 10,000 total counts were excluded from further analysis, as well as cells with less than 5,000 or greater than 15,000 total counts, resulting in 8,219 cells and 213 genes. 100 pairs of genes were then chosen randomly.

### Limitations

Like GMMs and *k*-means clustering, selection of *k* is challenging in the case of non-Gaussian distributed data without a Bayesian prior. We found ICL performed better than BIC, but it was still insufficient on its own, although combination with an automated elbow method (59) improves performance. There is also the problem of selecting the proper dimension *d*_*i*_ for each cluster, a known challenge in PCA. Scree plots or cross validation are typically used for PCA, but a proposed Bayesian method using a prior may be a viable approach to implement (60). Until methods for selection of *k* or subspace dimension are validated, we emphasize caution in correlating the latent variable(s) of a cluster with observed data points or using the explained variance in a Scree plot as a method for validating clustering. For example, data generated on the surface of a 2-D square could be (mis)partitioned into clusters from fitting the data with 2 lines. Unlike BIC, the ICL criterion discourages this, but should it occur, each cluster will indeed appear more line-like when viewed in isolation as the clustering method itself implies a single latent variable per cluster. This is reminiscent of a statistical “double dipping” problem with differential expression analysis after clustering addressed by Gao, Bien, and Witten in (61). Viewing clusters together will be of benefit in this scenario.

## References

1. Michael E. Tipping and Christopher M. Bishop. Probabilistic Principal Component Analysis. Journal of the Royal Statistical Society Series B: Statistical Methodology, 61(3):611–622, September 1999. ISSN 1369-7412, 1467-9868. doi: 10.1111/1467-9868.00196.

2. Sam Roweis. EM Algorithms for PCA and SPCA. pages 626–632, Denver, Colorado, USA, July 1998.

3. Geoffrey E Hinton, Michael Revow, and Peter Dayan. Recognizing Handwritten Digits Using Mixtures of Linear Models. 1995.

4. Zoubin Ghahramani and Geoffrey E. Hinton. The EM Algorithm for Mixtures of Factor Analyzers. Technical Report CRG-TR-96-1, May 1996.

5. Ting Su and Jennifer G. Dy. Automated hierarchical mixtures of probabilistic principal component analyzers. In Twenty-first international conference on Machine learning - ICML ‘04, page 98, Banff, Alberta, Canada, 2004. ACM Press. doi: 10.1145/1015330.1015393.

6. Michael E Tipping and Christopher M Bishop. Mixtures of Probabilistic Principal Component Analyzers. Neural Computation, 11(2):443–482, February 1999. doi: 10.1162/089976699300016728.

7. Vikas Trivedi, Harry M. T. Choi, Scott E. Fraser, and Niles A. Pierce. Multidimensional quantitative analysis of mRNA expression within intact vertebrate embryos. Development, 145(1):dev156869, January 2018. ISSN 1477-9129, 0950-1991. doi: 10.1242/dev.156869.

8. Harry M. T. Choi, Maayan Schwarzkopf, Mark E. Fornace, Aneesh Acharya, Georgios Artavanis, Johannes Stegmaier, Alexandre Cunha, and Niles A. Pierce. Third-generation in situ hybridization chain reaction: Multiplexed, quantitative, sensitive, versatile, robust. Development, 145:dev165753, 2018. doi: 10.1242/dev.165753.

9. Samuel J. Schulte, Mark E. Fornace, John K. Hall, Grace J. Shin, and Niles A. Pierce. HCR spectral imaging: 10-plex, quantitative, high-resolution RNA and protein imaging in highly autofluorescent samples. Development, 151:dev202307, 2024.

10. Stephen H. Devoto, Ellie Melançon, Judith S. Eisen, and Monte Westerfield. Identification of separate slow and fast muscle precursor cells in vivo, prior to somite formation. Development, 122(11):3371–3380, November 1996. ISSN 0950-1991, 1477-9129. doi: 10.1242/dev.122.11.3371.

11. Samuel R. Keenan and Peter D. Currie. The Developmental Phases of Zebrafish Myogenesis. Journal of Developmental Biology, 7(2):12, June 2019. ISSN 2221-3759. doi: 10.3390/jdb7020012.

12. Georgina E. Hollway, Robert J. Bryson-Richardson, Silke Berger, Nicholas J. Cole, Thomas E. Hall, and Peter D. Currie. Whole-Somite Rotation Generates Muscle Progenitor Cell Compartments in the Developing Zebrafish Embryo. Developmental Cell, 12(2): 207–219, February 2007. ISSN 15345807. doi: 10.1016/j.devcel.2007.01.001.

13. Frank Stellabotte, Betsy Dobbs-McAuliffe, Daniel A. Fernández, Xuesong Feng, and Stephen H. Devoto. Dynamic somite cell rearrangements lead to distinct waves of myotome growth. Development, 134(7):1253–1257, April 2007. ISSN 1477-9129, 0950-1991. doi: 10.1242/dev.000067.

14. Michaela Yuen, Sandra T. Cooper, Steve B. Marston, Kristen J. Nowak, Elyshia McNamara, Nancy Mokbel, Biljana Ilkovski, Gianina Ravenscroft, John Rendu, Josine M. De Winter, Lars Klinge, Alan H. Beggs, Kathryn N. North, Coen A.C. Ottenheijm, and Nigel F. Clarke. Muscle weakness in TPM3-myopathy is due to reduced Ca2+-sensitivity and impaired actomyosin cross-bridge cycling in slow fibres. Human Molecular Genetics, 24(22):6278–6292, November 2015. ISSN 0964-6906, 1460-2083. doi: 10.1093/hmg/ddv334.

15. Michael A. Geeves, Sarah E. Hitchcock-DeGregori, and Peter W. Gunning. A systematic nomenclature for mammalian tropomyosin isoforms. Journal of Muscle Research and Cell Motility, 36(2):147–153, April 2015. ISSN 0142-4319, 1573-2657. doi: 10.1007/s10974-014-9389-6.

16. Kathy Pieples and David F. Wieczorek. Tropomyosin 3 Increases Striated Muscle Isoform Diversity. Biochemistry, 39(28):8291–8297, July 2000. ISSN 0006-2960, 1520-4995. doi: 10.1021/bi000047x.

17. Aline Bonnet, Guillaume Lambert, Sylvain Ernest, François Xavier Dutrieux, Fanny Coulpier, Sophie Lemoine, Riadh Lobbardi, and Frédéric Marc Rosa. Quaking RNA-Binding Proteins Control Early Myofibril Formation by Modulating Tropomyosin. Developmental Cell, 42(5): 527–541.e4, September 2017. ISSN 15345807. doi: 10.1016/j.devcel.2017.08.004.

18. Mai E. Nguyen-Chi, Robert Bryson-Richardson, Carmen Sonntag, Thomas E. Hall, Abigail Gibson, Tamar Sztal, Wendy Chua, Thomas F. Schilling, and Peter D. Currie. Morphogenesis and Cell Fate Determination within the Adaxial Cell Equivalence Group of the Zebrafish Myotome. PLoS Genetics, 8(10):e1003014, October 2012. ISSN 1553-7404. doi:10.1371/journal.pgen.1003014.

19. Harunobu Kametani, Yue Tong, Atsuko Shimada, Hiroyuki Takeda, Takamichi Sushida, Masakazu Akiyama, and Toru Kawanishi. Twisted cell flow facilitates three-dimensional somite morphogenesis in zebrafish. Cells & Development, 180:203969, December 2024. ISSN 26672901. doi: 10.1016/j.cdev.2024.203969.

20. Phong Dang Nguyen, Georgina Elizabeth Hollway, Carmen Sonntag, Lee Barry Miles, Thomas Edward Hall, Silke Berger, Kristine Joy Fernandez, David Baruch Gurevich, Nicholas James Cole, Sara Alaei, Mirana Ramialison, Robert Lyndsay Sutherland, Jose Maria Polo, Graham John Lieschke, and Peter David Currie. Haematopoietic stem cell induction by somite-derived endothelial cells controlled by meox1. Nature, 512(7514): 314–318, August 2014. ISSN 0028-0836, 1476-4687. doi: 10.1038/nature13678.

21. Eugene Andres Houseman, William P Accomando, Devin C Koestler, Brock C Christensen, Carmen J Marsit, Heather H Nelson, John K Wiencke, and Karl T Kelsey. DNA methylation arrays as surrogate measures of cell mixture distribution. BMC Bioinformatics, 13(1):86, December 2012. ISSN 1471-2105. doi: 10.1186/1471-2105-13-86.

22. Jennifer Listgarten, Christoph Lippert, Carl M Kadie, Robert I Davidson, Eleazar Eskin, and David Heckerman. Improved linear mixed models for genome-wide association studies. Nature Methods, 9(6):525–526, June 2012. ISSN 1548-7091, 1548-7105. doi: 10.1038/nmeth.2037.

23. Elior Rahmani, Noah Zaitlen, Yael Baran, Celeste Eng, Donglei Hu, Joshua Galanter, Sam Oh, Esteban G Burchard, Eleazar Eskin, James Zou, and Eran Halperin. Sparse PCA corrects for cell type heterogeneity in epigenome-wide association studies. Nature Methods, 13(5):443–445, May 2016. ISSN 1548-7091, 1548-7105. doi: 10.1038/nmeth.3809.

24. Alkes L. Price, Noah A. Zaitlen, David Reich, and Nick Patterson. New approaches to population stratification in genome-wide association studies. Nature Reviews Genetics, 11 (7):459–463, July 2010. ISSN 1471-0056, 1471-0064. doi: 10.1038/nrg2813.

25. Esteban Burchard. Differential dna methylation in latino population. https://www.ncbi.nlm.nih.gov/geo/query/acc.cgi?acc=GSE77716, 2 2016.

26. Lovisa E. Reinius, Nathalie Acevedo, Maaike Joerink, Göran Pershagen, Sven-Erik Dahlén, Dario Greco, Cilla Söderhäll, Annika Scheynius, and Juha Kere. Differential DNA Methylation in Purified Human Blood Cells: Implications for Cell Lineage and Studies on Disease Susceptibility. PLoS ONE, 7(7):e41361, July 2012. ISSN 1932-6203. doi: 10.1371/journal.pone.0041361.

27. Jonathan S. Packer, Qin Zhu, Chau Huynh, Priya Sivaramakrishnan, Elicia Preston, Hannah Dueck, Derek Stefanik, Kai Tan, Cole Trapnell, Junhyong Kim, Robert H. Waterston, and John I. Murray. A lineage-resolved molecular atlas of C. elegans embryogenesis at single-cell resolution. Science, 365(6459):eaax1971, September 2019. ISSN 0036-8075, 1095-9203. doi: 10.1126/science.aax1971.

28. Laurens van der Maaten and Geoffrey Hinton. Visualizing Data using t-SNE. Journal of Machine Learning Research, 9:2579–2605, 2008.

29. Leland McInnes, John Healy, and James Melville. UMAP: Uniform Manifold Approximation and Projection for Dimension Reduction, September 2020. 1802.03426 [stat].

30. Dmitry Kobak and Philipp Berens. The art of using t-SNE for single-cell transcriptomics. Nature Communications, 10(1):5416, November 2019. ISSN 2041-1723. doi: 10.1038/s41467-019-13056-x.

31. Tara Chari and Lior Pachter. The specious art of single-cell genomics. PLOS Computational Biology, 19(8):e1011288, August 2023. ISSN 1553-7358. doi: 10.1371/journal.pcbi.1011288.

32. PS Bradley and OL Mangasarian. k-Plane Clustering. Journal of Global Optimization, 16: 23–32, 2000. doi: 10.1023/A:1008324625522.

33. Pankaj K Agarwal and Nabil H Mustafa. k-Means Projective Clustering. In PODS ‘04: Proceedings of the twenty-third ACM SIGMOD-SIGACT-SIGART symposium on Principles of database systems, pages 155–165, Paris, France, June 2004. Association for Computing Machinery. ISBN 978-1-58113-858-0.

34. Teng Zhang, Arthur Szlam, and Gilad Lerman. Median K-flats for hybrid linear modeling with many outliers. In Proceedings of 2nd IEEE International Workshop on Subspace Methods, pages 234–241, Kyoto, Japan, September 2009. IEEE. doi: 10.1109/ICCVW.2009.5457695.

35. René Vidal. A tutorial on subspace clustering. 2010.

36. Andrew Gitlin, Biaoshuai Tao, Laura Balzano, and John Lipor. Improving k-subspaces via coherence pursuit. IEEE Journal of Selected Topics in Signal Processing, 12(6):1575–1588, December 2018. ISSN 1932-4553, 1941-0484. doi: 10.1109/JSTSP.2018.2869363.

37. John Lipor, David Hong, Yan Shuo Tan, and Laura Balzano. Subspace clustering using ensembles of K-subspaces. Information and Inference: A Journal of the IMA, 10(1):73– 107, March 2021. ISSN 2049-8772. doi: 10.1093/imaiai/iaaa031.

38. John Aldrich. Doing Least Squares: Perspectives from Gauss and Yule. International Statistical Review, 66(1):61–81, April 1998. doi: 10.2307/1403657.

39. Robin L. Plackett. A Historical Note on the Method of Least Squares. Biometrika, 36(3/4): 458–460, December 1949. doi: 10.2307/2332682.

40. Robert J. Adcock. A Problem in Least Squares. The Analyst, 5(2):53–54, March 1978. doi: 10.2307/2635758.

41. W. Edwards Deming. The General Problem In Least Squares. In Statistical adjustment of data, pages 49–51. John Wiley & Sons, Inc, New York, 1943.

42. Cédric Archambeau, Nicolas Delannay, and Michel Verleysen. Mixtures of robust probabilistic principal component analyzers. Neurocomputing, 71(7-9):1274–1282, March 2008. ISSN 09252312. doi: 10.1016/j.neucom.2007.11.029.

43. Julien Chiquet, Mahendra Mariadassou, and Stéphane Robin. Variational inference for probabilistic Poisson PCA. The Annals of Applied Statistics, 12(4), December 2018. ISSN 1932-6157. doi: 10.1214/18-AOAS1177.

44. Maayan Schwarzkopf, Mike C. Liu, Samuel J. Schulte, Rachel Ives, Naeem Husain, Harry M. T. Choi, and N. A. Pierce. Hybridization chain reaction enables a unified approach to multiplexed, quantitative, high-resolution immunohistochemistry and in situ hybridization. Development, 148(22):dev199847, 2021. ISSN 0950-1991, 1477-9129. doi: 10.1242/dev.199847.

45. Samuel J. Schulte, Boyoung Shin, Ellen V. Rothenberg, and Niles A. Pierce. Multiplex, quantitative, high-resolution imaging of protein:protein complexes via hybridization chain reaction. ACS Chem. Biol., 19(2):280–288, 2024.

46. Jadzia Livingston, Melanie A. Spero, Zachery R. Lonergan, and Dianne K. Newman. Visualization of mRNA Expression in Pseudomonas aeruginosa Aggregates Reveals Spatial Patterns of Fermentative and Denitrifying Metabolism. Applied and Environmental Microbiology, 88(11):e00439–22, June 2022. ISSN 0099-2240, 1098-5336. doi: 10.1128/aem.00439-22.

47. Sebastian Preissl, Kyle J. Gaulton, and Bing Ren. Characterizing cis-regulatory elements using single-cell epigenomics. Nature Reviews Genetics, 24(1):21–43, January 2023. ISSN 1471-0056, 1471-0064. doi: 10.1038/s41576-022-00509-1.

48. Peter Langfelder and Steve Horvath. WGCNA: an R package for weighted correlation network analysis. BMC Bioinformatics, 9(1):559, December 2008. ISSN 1471-2105. doi: 10.1186/1471-2105-9-559.

49. Christophe Biernacki, Gilles Celeux, and Gérard Govaert. Assessing a mixture model for clustering with the integrated completed likelihood. IEEE Transactions on Pattern Analysis and Machine Intelligence, 22(7):719–725, July 2000. ISSN 01628828. doi: 10.1109/34.865189.

50. Marco Bertoletti, Nial Friel, and Riccardo Rastelli. Choosing the number of clusters in a finite mixture model using an exact Integrated Completed Likelihood criterion, May 2015. 1411.4257 [stat].

51. Christopher M. Bishop and Michael E. Tipping. A hierarchical latent variable model for data visualization. IEEE Transactions on Pattern Analysis and Machine Intelligence, 20(3): 281–293, March 1998. ISSN 01628828. doi: 10.1109/34.667885.

52. Carl Eckart and Gale Young. The approximation of one matrix by another of lower rank. Psychometrika, 1:211–218, September 1939. doi: 10.1007/BF02288367.

53. Zeynep F Altun and David H Hall. Epithelial system, hypodermis. In WormAtlas. 2009.

54. Zeynep F Altun and David H Hall. Epithelial system, seam cells. In WormAtlas. 2009.

55. Paul D. Thomas, Michael J. Campbell, Anish Kejariwal, Huaiyu Mi, Brian Karlak, Robin Daverman, Karen Diemer, Anushya Muruganujan, and Apurva Narechania. PANTHER: A Library of Protein Families and Subfamilies Indexed by Function. Genome Research, 13(9): 2129–2141, September 2003. ISSN 1088-9051. doi: 10.1101/gr.772403.

56. Huaiyu Mi, Anushya Muruganujan, Xiaosong Huang, Dustin Ebert, Caitlin Mills, Xinyu Guo, and Paul D. Thomas. Protocol Update for large-scale genome and gene function analysis with the PANTHER classification system (v.14.0). Nature Protocols, 14(3):703–721, March 2019. ISSN 1754-2189, 1750-2799. doi: 10.1038/s41596-019-0128-8.

57. F. Alexander Wolf, Philipp Angerer, and Fabian J. Theis. SCANPY: large-scale single-cell gene expression data analysis. Genome Biology, 19(1):15, December 2018. ISSN 1474-760X. doi: 10.1186/s13059-017-1382-0.

58. Samuel L. Wolock, Romain Lopez, and Allon M. Klein. Scrublet: Computational Identification of Cell Doublets in Single-Cell Transcriptomic Data. Cell Systems, 8(4):281–291.e9, April 2019. ISSN 24054712. doi: 10.1016/j.cels.2018.11.005.

59. Ville Satopaa, Jeannie Albrecht, David Irwin, and Barath Raghavan. Finding a “Kneedle” in a Haystack: Detecting Knee Points in System Behavior. In ICDCSW ‘11: Proceedings of the 2011 31st International Conference on Distributed Computing Systems Workshops, pages 166–171, Minneapolis, MN, USA, June 2011. doi: 10.1109/ICDCSW.2011.20.

60. Thomas Minka. Automatic choice of dimensionality for PCA. MIT Media Laboratory Perceptual Compution Section Technical Report No. 514, December 2000.

61. Lucy L. Gao, Jacob Bien, and Daniela Witten. Selective Inference for Hierarchical Clustering. October 2022. doi: 10.48550/arXiv.2012.02936. 2012.02936 [stat].

