## Supplement for "*k*-spaces: Mixtures of Gaussian latent variable models"

#### 1: Exposition on PCA and pPCA.

***1.1: Review of PCA.*** Principal component analysis, or PCA, is a common dimensionality reduction method used in biology, and is often specifically used to visualize data in 2-D plots. These visualizations are the projections of the data onto principal components— typically the first two. These principal components themselves are linear combinations of features in the data (i.e. directions in feature space). They are perpendicular directions in the feature space chosen such that the first component is the direction of maximum variance, the second is that with the most variance after removing the first component, and so on. Essentially, PCA defines a new coordinate system in our feature space, and it could be viewed as a rotation of our starting axes (maintaining the orthogonality) followed by dropping "less important" axes that contain less variance from our visualization. As an analogy for 2-D PCA of 3-D data, one could consider entering a rectangular room and seeking the best angle to take a photograph where all subjects can be seen. The walls of the room and the doorway provide a natural set of axes and "origin" from which to frame the photograph, but some subjects may block each other from view so finding a new vantage point may be better. Depth perception is lost in the photograph, and this is akin to discarding variance from our 3-D space to produce the 2-D projection.

Mathematically, determining this coordinate system is equivalent to finding a subspace that is "closest" to the data, meaning it minimizes the Frobenius norm or sum of squared distances ($L_2$ norms) from the data points to the subspace. (Note the similarity to linear regression, which uses residuals in a dependent variable while this PCA procedure uses orthogonal residuals, implying errors in all variables.) It is the optimal linear dimensionality reduction method in the sense that it retains the maximum variance in the data, or "spaces out" the plotted points as much as is possible without a nonlinear distortion (1). Unlike tSNE and UMAP, both local and global distances are meaningful, though points that appear next to each other in a plot may be further apart in another, discarded dimension (2–4).

The process of minimizing the sum of squared residuals and the goal of retaining the maximum variance (implying that we are minimizing the discarded variance) suggest a relationship with the Normal or Gaussian distribution and the covariance matrix. Indeed, one way to obtain PCA is to construct the covariance matrix of the data and then use singular value decomposition (SVD) to decompose it into its eigenvalues and eigenvectors, also called the spectral decomposition. The eigenvectors with the largest corresponding eigenvalues (or variances of projections of the data along the eigenvectors) are the principal components. Dropping lower PCs is equivalent to ignoring the components of our data along the directions of our lower eigenvectors. Thus, we can also arrive at PCA by fitting a Gaussian to the data first and then determining how the distribution is "stretched" in different directions to find those with the greatest variance.

***1.2: The relationship between PCA and pPCA.*** Despite the above, PCA is not a probabilistic model for subtle reasons, and this is hinted by the slight discrepancy between our first definition- the optimal linear dimensionality reduction that maximizes retained variance- and our second definition- the basis vectors of the affine subspace closest to the data point. The second definition is subtly wrong as it is too restrictive. Finding the closest subspace produces PCA, but PCA does not necessarily find the closest subspace. This is because the notion of a "closest" affine subspace implies a specific translation of our subspace. In our photograph analogy, if we already have determined the optimal angle to take the snapshot, moving the camera forwards or backwards while maintaining the angle of view will not change the relative positions of subjects in our photograph. (This analogy neglects the fact that a camera in real life takes an image of subjects facing the focal point of the curved lens, and moving the camera into the midst of the subjects will change their relative positioning.) Likewise, any affine subspace that shares the same basis vectors (or relative orientation) with the closest affine subspace will produce the same projection. Translation of the subspace along the discarded coordinates affects the projection in neither an absolute nor relative way. There are infinitely many affine subspaces that we can find to obtain PCA, though despite this degeneracy they all share the same basis. Of note, if the data is already centered prior to PCA, then we can amend our second definition to state it is the vector subspace (as opposed to an affine subspace) that is closest to the data, and our two definitions will be equivalent.

However, despite resolving the degeneracy of translation with a stricter definition, it is still not a probabilistic model despite the following probabilistic interpretations for PCA. Definition 3 is that PCA is the result of maximizing the likelihood of residuals for a set of points with isotropic noise (i.e., a generalization of linear regression to errors-in-variables that directly follows from Definition 2) (5). Definition 4 for PCA is that it is the result of fitting a Gaussian and dropping lower dimensions. First, Definition 3 does not consider the latent space distribution; therefore, it is not fully probabilistic. It may seem this can be resolved with Definition 4, with the reasoning that as Gaussians are probabilistic models and PCA results from the process of fitting a Gaussian, PCA should be probabilistic. However, one can show that PCA can be produced by finding the correct rotation matrix to apply to any arbitrary diagonal covariance matrix with distinct entries (meaning the eigenvalues are fixed beforehand) to fit a Gaussian to the data and then obtaining the top eigenvectors of the resulting covariance matrix. Therefore, PCA does not actually describe the process for generating the data, as fitting many Gaussians can produce the same principal components. This emphasizes a key point of confusion about PCA. The axes are indeed interpretable unlike UMAP and tSNE (2), and it can be viewed as the result of a probabilistic inference procedure for a subspace. However, they are the result of infinitely many probabilistic models and thus PCA does not define any one probabilistic model. Of the many Gaussian probabilistic models, there are a subset that produce the optimal marginal distribution for maximizing the data likelihood.

One such optimal model is probabilistic PCA (pPCA), which models isotropic noise added to Gaussian latent variables and produces a posterior distribution for inference of the latent points. Another is ours, which assumes isotropic noise in the complementary dimension, in which the maximum likelihood latent points are the projections. The marginal distributions for the observed data are the same (adding Gaussian noise to a Gaussian variable produces a new variable that is also Gaussian-distributed) and thus the clustering of data and fitting of subspaces is the same in the mixture case.

***1.3: The generative model for pPCA.*** Of note, there may be some confusion as well regarding the isotropic noise in pPCA, and it is particularly important to understanding how the noise is modeled in *k*-spaces as well, as the marginal distribution is the same. pPCA, as posed by Bishop and Tipping (6), has Gaussian-distributed latent points perturbed by isotropic Gaussian noise, meaning equal, normally-distributed noise in all directions. However, the data is *not* generated by sampling a latent point from a Gaussian distribution, picking a direction uniformly at random, and then sampling a distance from a Half-normal distribution to perturb the latent point to obtain an observed data point (we will refer to this as the radial Gaussian). The correct generative process is to sample a latent point from a Gaussian distribution, and then for each of the D dimensions of the observed data, perturb the data by distances independently sampled from the same Gaussian (or equivalently from an isotropic D-dimensional multivariate Gaussian). The D-dimensional multivariate Gaussian is rotationally symmetric, as is the radial Gaussian by construction. Both *are* isotropic. However, they produce data with distinct distributions. Consider the 2-D case of each, centered at the origin. Start with the radial Gaussian. If we generate data and want to know the distribution of points lying exactly along the $x$ axis, this would be the subset of points with exactly $\theta = 0$ or 180°. As $y = 0$ for this set of points, their $x$ coordinate is exactly equal to the distance from the origin, or the orthogonal residual length from our 0-D subspace. The conditional distribution is Gaussian. Rotational symmetry and our described generative process generalizes this argument to any angle of $\theta$, so since $\theta$ is sampled uniformly at random, the marginal distribution of residual lengths would be the same Gaussian. Now consider the bivariate isotropic Gaussian, and again examine the set of points lying exactly on the $x$ axis. The residuals of these points have a Gaussian conditional distribution as well. Their residuals are merely the first of the two independently sampled Gaussian variables that together give us each point. This distribution is rotationally symmetric as well. It would seem that this would produce a Gaussian marginal distribution of residual lengths. However, it does not. If we forgo using a conditional distribution and integrating and instead directly compute our marginal distribution based on independence of the $x$ and $y$ values of the data, we see that the orthogonal residuals are $\sqrt{x^2+y^2}$, where $X$ and $Y$ are Gaussian. This is a Rayleigh distribution.

The distinction in generative models (picking a direction at random and then perturbing vs. perturbing independently in orthogonal directions) is subtle. The key issue is the limiting process for obtaining the set of points along the $x$ axis in each scenario. This is known as the Borel-Kolmogorov paradox (7), and it arose because we must ensure that our conditional density is derived in the same coordinate system as we intend to integrate in to obtain the marginal distribution. In our bivariate Gaussian from pPCA, we picked an $x$ coordinate and a $y$ coordinate separately, simultaneously determining angle and distance. Our conditional density of points along the $x$ axis was obtained as $y \to 0$, i.e., as we squished a rectangle onto the $x$ axis. In contrast, squishing a wedge onto the $x$ axis as $\theta \to 0$ will produce a different conditional density- the Rayleigh distribution. Mentally "integrating" out the angle by using rotational symmetry mixed a conditional density from Cartesian coordinates with an integration in polar coordinates, producing our paradox.

**2: Derivation of variance as a function of residuals.** In ordinary least squares, minimizing the residual sum of squares is equivalent to maximizing the likelihood of a model with normally distributed residuals. PCA corresponds to multidimensional orthogonal linear regression, meaning that it minimizes the sum of squared orthogonal residuals, or squared distances of points to their projections on the subspace. PCA could also be viewed as maximizing $f_{X_\perp}$ for observing a set of $N$ points in $\mathbb{R}^D$, where the observed points are generated from some latent distribution and then given noise from multivariate Gaussians $\mathcal{N}(\text{proj}_s(\mathbf{x}), \sigma^2\mathbf{I}_D)$ (5). One can rotate the covariance matrix of these Gaussians such that $d$ axes of our new coordinate system match the $d$ principal components. Thus, the observed point's complementary component of its likelihood is given by an isotropic multivariate Gaussian in the complementary space $\mathcal{N}(\text{proj}_s(\mathbf{x}), \sigma^2 \cdot \mathbf{I}_{D-d})$:

$$f_{X_\perp}(\mathbf{x}|s) = \prod_{a=d+1}^{D} \frac{1}{\sqrt{2\pi}\sigma} e^{-\frac{1}{2}\frac{(x_a - proj_s(\mathbf{x})_a)^2}{\sigma^2}}. \tag{1}$$

Let $n_i = D - d_i$ for convenience, and define $r$ as the length of the orthogonal residual from $\mathbf{x}$ to its projection on space $s$. Let $j$ index individual points $\mathbf{x}_j$ and $a$ index dimensions for a space, such that $x_a$ refers to the value of point $\mathbf{x}$ in the $a$th dimension of a space's coordinate system. Rotational symmetry allows the likelihood of a point to be written as a function of its residual's length:

$$f_{X_\perp}(r|s) = \frac{1}{(2\pi)^{n_i/2}\sigma^{n_i}} e^{-\frac{1}{2}\frac{r^2}{\sigma^2}}. \tag{2}$$

$\sigma_i^2$ is the mean variance over the complementary dimensions, as noted in (6), and we show this below for completeness. Once the spaces have been fitted, the expected variance for each space, $\sigma_i^2$, over its complementary dimensions is given by:

$$\begin{aligned}\sigma_i^2 &= \mathbb{E}_{a\in\perp}[(x_{ja} - \text{proj}_{s_i}(\mathbf{x}_j)_a)^2] - \mathbb{E}_{a\in\perp}[(x_{ja} - \text{proj}_{s_i}(\mathbf{x}_j)_a)]^2, \qquad (3)\\ &= \sum_{j=1}^{N} P(s_i|\mathbf{x}_j)\frac{1}{n_i}\sum_{a=1}^{n_i}(x_{ja} - \text{proj}_{s_i}(\mathbf{x}_j)_a)^2 - \left(\sum_{j=1}^{N_i} P(s_i|\mathbf{x}_j)\frac{1}{n_i}\sum_{a=1}^{n_i}(x_{ja} - \text{proj}_{s_i}(\mathbf{x}_j)_a)\right)^2.\end{aligned}$$

In the case of hard assignment where $N_i$ indicates the number of points assigned to space $i$, this becomes:

$$\sigma_i^2 = \frac{1}{N_i}\sum_{i=1}^{N_i}\frac{1}{n}\sum_{a=1}^{n}(x_{ja} - \text{proj}_{s_i}(\mathbf{x}_j)_a)^2 - \left(\frac{1}{N_i}\sum_{i=1}^{N_i}\frac{1}{n}\sum_{a=1}^{n}(x_{ja} - \text{proj}_{s_i}(\mathbf{x}_j)_a)\right)^2.$$

As minimizing the residuals means the second term is 0,

$$\begin{aligned}\sigma_i^2 &= \sum_{j=1}^{N_i} P(s_i|\mathbf{x}_j)\frac{1}{n_i}\sum_{a=1}^{n_i}(x_{ja} - \text{proj}_{s_i}(\mathbf{x}_j)_a)^2 - 0,\\ &= \frac{1}{N_i n_i}\sum_{j=1}^{N_i} r_{ij}^2. \qquad (4)\end{aligned}$$

In soft, probabilistic assignment, this becomes

$$= \frac{1}{N\pi_i n_i}\sum_{j=1}^{N} P(s_i|\mathbf{x}_j) r_{ij}^2. \tag{5}$$

In some cases, we may also want to assume noise-generating process is the same for all spaces; that is, $\sigma_i^2$ is equal for all spaces. While noise within the latent spaces is not explicitly modeled, one can simply subtract the corresponding variance from the covariance matrix in $f_{X_\parallel}(\mathbf{x}|s)$ to make an isotropic $f_{X_{\text{noise}}}(\mathbf{x}|s)$ that includes both the latent and complementary dimensions. This relies on the assumption that the per-dimension estimated shared noise parameter is not greater than the per-dimension variances within any of the latent spaces, and in our software we enforce this constraint. In their paper introducing probabilistic PCA, where Bishop and Tipping note that the noise parameter corresponds to the average per-dimension variance discarded by a pPCA model, it is known how many dimensions are discarded (6). Here, where we have a single shared $\sigma^2$ rather than a $\sigma_i^2$ unique to each space and additionally allow $d_i$ to vary between mixture components, this quantity is unknown because $d_i$ varies between mixture components. $\sigma^2$ therefore corresponds to the *expected* variance discarded, and the denominator is the *expected* number of direct observations of our noise process. Let $\pi_{ij}$ be a shorthand denoting the responsibility of space $s_i$ for point $\mathbf{x}_j$ ($\pi_{ij} = P(s_i|\mathbf{x}_j)$), and use the identity $\sum_{j=1}^{N}\pi_{ij} = N\pi_i$.

$$\sigma^2 = \frac{1}{N\sum_{i=1}^{k}\pi_i n_i}\sum_{j=1}^{N}\sum_{i=1}^{k}\pi_{ij} r_{ij}^2. \tag{6}$$

Intuitively, we arrive at this formula because our model specifies each complementary dimension of each space contains a conditionally independent observation of the same noise process controlled by parameter $\sigma^2$, so rather than jointly fitting the dimensions of a $\sum_{i=1}^{k} d_i$-dimensional isotropic multivariate distribution, we can treat the problem as a univariate distribution of a single random variable for noise, $Z$, where each entry $x_{ja}^\perp$ in each $\mathbf{x}_j^\perp$ from each space is an independent sampling of $Z$. If we knew which space generated each point or performed hard assignment, we would have $\sum_{j=1}^{N}\sum_{i=1}^{k} n_i 1_{\mathbf{x}_j \in \mathcal{S}_i}$ observations of $Z$; however, the number of total observations of $Z$ depends on which of the spaces generated each point $\mathbf{x}_j$, as potentially each space generates a vector $\mathbf{x}^\perp$ with different dimension. This can also be shown with a maximum likelihood approach for the optimal choice of $\sigma^2$ in the M step. We begin with the expected complete data log-likelihood:

$$
\begin{aligned}
\mathcal{L} &= \sum_{j=1}^{N}\sum_{i=1}^{k}\pi_{ij}\log\left[f_{\|}(\mathbf{x}_j^{\|}\mid s_i)\cdot f_{\perp}(\mathbf{x}_j^{\perp}\mid s_i)\right], \\
&= \sum_{j=1}^{N}\sum_{i=1}^{k}\pi_{ij}\log\left[f_{\|}(\mathbf{x}_j^{\|}\mid s_i)\right] + \sum_{j=1}^{N}\sum_{i=1}^{k}\pi_{ij}\log\left[f_{\perp}(\mathbf{x}_j^{\perp}\mid s_i)\right].
\end{aligned} \tag{7}
$$

We can discard the first term, as it is independent of the second during optimization and will be 0 when we later take the derivative of $\mathcal{L}$ with respect to $\sigma$.

$$
\begin{aligned}
\mathcal{L}^{\perp} &= \sum_{i=1}^{k}\sum_{j=1}^{N}\pi_{ij}\log\Big(\prod_{a=1}^{n_i}\frac{1}{\sqrt{2\pi}\sigma}\exp\big(-\frac{1}{2}\frac{(\mathbf{x}_j-\text{proj}_{s_i}(\mathbf{x}_j))_a^2}{\sigma^2}\big)\Big), \\
&= \sum_{i=1}^{k}\sum_{j=1}^{N}\left[\pi_{ij}n_i\log\Big(\frac{1}{\sqrt{2\pi}\sigma}\Big)-\frac{\pi_{ij}}{2}\sum_{a=1}^{n_i}\frac{(\mathbf{x}_j-\text{proj}_{s_i}(\mathbf{x}_j))_a^2}{\sigma^2}\right], \\
&= \sum_{i=1}^{k}\sum_{j=1}^{N}-\pi_{ij}n_i\frac{1}{2}\log(2\pi\sigma^2)-\frac{1}{2}\sum_{i=1}^{k}\sum_{j=1}^{N}\pi_{ij}\frac{r_{ij}^2}{\sigma^2}, \\
&= \sum_{i=1}^{k}\sum_{j=1}^{N}-\pi_{ij}n_i\frac{1}{2}\log(2\pi\sigma^2)-\frac{1}{2}\frac{1}{\sigma^2}\sum_{i=1}^{k}\sum_{j=1}^{N}\pi_{ij}r_{ij}^2, \\
&= \sum_{i=1}^{k}\sum_{j=1}^{N}-\pi_{ij}n_i\Big(\frac{1}{2}\log(2\pi)+\log(\sigma)\Big)-\frac{1}{2}\frac{1}{\sigma^2}\sum_{i=1}^{k}\sum_{j=1}^{N}\pi_{ij}r_{ij}^2.
\end{aligned}
$$

Setting the derivative to zero,

$$
\frac{d\mathcal{L}^{\perp}}{d\sigma} = -\frac{1}{\sigma}\sum_{i=1}^{k}\sum_{j=1}^{N}\pi_{ij}n_i + \frac{1}{\sigma^3}\sum_{i=1}^{k}\sum_{j=1}^{N}\pi_{ij}r_{ij}^2 = 0,
$$

and solving yields equation 12.

**3: Analytical consideration of the methylation simulation.** The hierarchical generative model for simulating methylation data is considerably more complex than the $k$-spaces model used to fit the data. Here, we study the realized distribution of the observed data. First, we show how centering allows for a convenient decomposition of the data matrix into the sum of ($k$-1)-D and 0-D signal and noise matrices by eliminating the effects of variation in the underlying site-specific means from the hierarchical model. Both matrices contribute to DMRs, while only the noise matrix contributes to non DMRs, and $k$-spaces fits these sites separately with different dimension subspaces.

Then, we examine the noise for each site across individuals, which are our features once the matrix is transposed, and find that it is Gaussian and approximately isotropic with regards to individuals. We next turn to the distribution over all entries in our noise matrix where varying $\sigma_j^2$ means that it is not Gaussian. However, under the assumption that the noise is isotropic, this distribution is monotonically decreasing with residual length. This result is our justification for approximating it in the E-step with Gaussians of equal variance for all spaces. Finally, we discuss some conditions under which EM with the $k$-spaces model will begin to fail.

Let $i$ always index rows corresponding to individuals, $j$ always index columns representing sites, and $k$ index cell types. There are $N$ individuals, $J$ sites, and $K$ cell types. In the simulation, $N = 500$, $J = 100{,}000$, $K = 5$. Our $N$ by $J$ methylation matrix is $\mathbf{O}$, and the centered matrix is $\mathbf{O}'$, but we will cluster on its transpose $\mathbf{O}'^{\mathrm{T}}$.

***3.1: Discussion of the generative process.*** Before examining the effect of centering in the decomposition, we discuss the generative process to set up our analysis. The hierarchical generative model, described in Methods, is reproduced below for convenience.

$$
\begin{aligned}
\mu_j &\sim \text{some distribution},\\
M_{jk} &\sim \text{Norm}(\mu_j, \tau^2),\\
\sigma_j &\sim \text{Halfnorm}(0, \sigma_*^2),\\
X_{ijk} &\sim \text{Norm}(M_{jk}, \sigma_j^2),\\
\mathbf{R_i} &\sim \text{Dir}(\alpha, K),\\
O_{ij} &= \sum_{k=1}^{K} X_{ijk} R_{ik} + \epsilon \text{ where } \epsilon \sim \text{Norm}(0, 0.01), \text{ truncated to } [0,1].
\end{aligned}
$$

We will show that the precise distributions of $\mu_j$ and $M_{jk}$ do not matter. The scale of variation in $M_{jk}$ does matter, however, because technical and biological noise are added. Some distribution on the simplex is needed to acquire individual cell type proportions $R_{ik}$. The precise distribution is again not critical, but as the variation in cell type proportions from one individual to another decreases, the signal-to-noise ratio shrinks. With no variation in proportions, DMRs will not be detected regardless of $\tau$. We will use a symmetric Dirichlet distribution here for this analysis. In simulation where site-specific variances $\sigma_1^2, ..., \sigma_M^2$ are sampled from $\text{Halfnormal}(0, 0.03^2)$, even as the concentration parameter, $\alpha$, becomes quite large ($\sim 100$), *k*-spaces clustering still works, although accuracy decreases (Fig. S26a). However, it fails around $\alpha = 40$ (Fig. S26b) when using $\sigma_j^2$ sampled from the empirical distribution in the reference data (8), which is slightly heavier tailed than the Half-normal and contains a very small fraction of extreme variances (Fig. S26). We note that the reference data is comprised of only 6 individuals and 5 measured cell types, and that the small subset of sites may be highly variable due to processes different from those governing methylation noise at the vast majority of sites. Eliminating all $\sigma_j > 0.2$ from the empirical distribution (the top 0.29% most variable sites) restores the robustness (Fig. S26c). Potential reasons for this will be discussed under "Complementary space likelihood of each point $\mathbf{O}_j^{(\mathbf{n})}$ decreases monotonically with residual length and classification is biased depending on distance from origin."

There is one more factor to consider, truncation of final methylation values to the range [0,1]. Sites with $\mu_j$ close to 0 or 1 could be problematic in two ways: 1) sites that are supposed to be DMRs may in practice not be DMRs if $\tau$ is too small or if by chance all 5 sampled $M_{jk}$ for that site are either out of bounds or within bounds but still very close to $\mu_j$ 2) biological and technical noise for these sites will not be distributed in the same manner as the rest due to the truncation. Moving forward, we will assume that $\mu_j$ is distributed in such a way that most sites have $\mu_j$ far enough away from the boundaries to avoid the second problem with the distribution of noise, and the first problem is merely a semantic issue of whether a site is a DMR because $M_{jk}$ "could" vary in principle or because the realizations of $M_{jk}$ vary "enough." Note that in this simulation, $\mu_j$ and $\sigma_j^2$ are independent; if $\sigma_j^2$ were smaller near the bounds in real data, that could reduce this issue. We also ignore $\epsilon$ for analytical simplicity but acknowledge it could be important in some regimes.

***3.2: Centering the data eliminates the effects of variation in site-specific means to allow decomposition into (k-1)-D and 0-D signal and noise matrices.*** Broadly, we want to examine the covariance structure of sites away from the vector $< 1, 1, 1, ..., 1 >$, or, in other words, correlation in "errors" off the value in the average individual. DMRs (features in $\mathbf{O}^{\mathrm{T}}$) will exhibit a covariance across individuals (dimensions in $\mathbf{O}^{\mathrm{T}}$), while non DMRs will have random noise. However, the underlying mean $\mu_j$ varies from site to site, interfering with this, so we center.

Our centered matrix is $\mathbf{O'}$. It is helpful to view $\mathbf{O'}$ as a sum of cell type proportion-driven variation and biological noise. To get there, we begin by decomposing $\mathbf{O'}$:

$$
\begin{aligned}
O'_{ij} &= O_{ij} - \frac{1}{N}\sum_{a=1}^{N} O_{aj},\\
&= \sum_{k=1}^{K} X_{ijk} R_{ik} - \frac{1}{N}\sum_{a=1}^{N}\sum_{k=1}^{K} X_{ajk} R_{ak},\\
&= \sum_{k=1}^{K} [X_{ijk} R_{ik} - \frac{1}{N}\sum_{a=1}^{N} X_{ajk} R_{ak}].
\end{aligned}
$$

Define $X'_{ijk} = X_{ijk} - M_{jk}$ noting that $X'_{ijk} \sim \mathrm{Norm}(0, \sigma_j^2)$. Then we obtain the following:

$$
\begin{aligned}
O'_{ij} &= \sum_{k=1}^{K} [R_{ik}(X_{ijk} - M_{jk} + M_{jk}) - \frac{1}{N}\sum_{a=1}^{N} R_{ak}(X_{ajk} - M_{jk} + M_{jk})], \\
&= \sum_{k=1}^{K} [R_{ik}X'_{ijk} + R_{ik}M_{jk} - \frac{1}{N}\sum_{a=1}^{N}(R_{ak}X'_{ajk} + R_{ak}M_{jk})], \\
&= \sum_{k=1}^{K} [R_{ik}X'_{ijk} + R_{ik}M_{jk} - \frac{1}{N}\sum_{a=1}^{N} R_{ak}X'_{ajk} - \frac{1}{N}\sum_{a=1}^{N} R_{ak}M_{jk}].
\end{aligned}
$$

$\mathbb{E}[\frac{1}{N}\sum_{a=1}^{N} R_{ak}X'_{ajk}] = 0$ because $\mathbb{E}[X'_{ajk}] = 0$ and $X'_{ajk}$ and $R_{ak}$ are independent, and in practice, $\frac{1}{N}\sum_{a=1}^{N} R_{ak}X'_{ajk}$ will be close to 0. It is a (randomly weighted) sample mean, and as $N \to \infty$, this converges to 0. Thus,

$$
O'_{ij} \approx \sum_{k=1}^{K} R_{ik}X'_{ijk} + \frac{1}{N}\sum_{a=1}^{N}\sum_{k=1}^{K}[R_{ik}M_{jk} - R_{ak}M_{jk}].
$$

We can then define a noise matrix $\mathbf{O^{(n)}}$, denoting biological noise resulting from variation in cell type methylation profiles between individuals, and a signal matrix $\mathbf{O^{(s)}}$, denoting the signal driven by the variation in cell type proportions:

$$
O_{ij}^{(\mathbf{n})} = \sum_{k=1}^{K} R_{ik}X'_{ijk},
$$

$$
\text{and } O_{ij}^{(\mathbf{s})} = \frac{1}{N}\sum_{a=1}^{N}\sum_{k=1}^{K}[(R_{ik} - R_{ak})M_{jk}].
$$

$$
\mathbf{O}' \approx \mathbf{O^{(n)}} + \mathbf{O^{(s)}}.
$$

It is important to note that our *k*-spaces model instead considers the decomposition $\mathbf{x}' = \mathbf{x}^{\perp} + \mathbf{x}^{\parallel}$ and does not explicitly model noise within the latent subspace, while our generative model for the DMR problem has noise in all directions as modeled explicitly by pPCA. However, this difference between *k*-spaces components and pPCA components is relevant only for inference of the latent points' positions and not for classification of points, as the resulting marginal distribution is the same Gaussian (and a *k*-spaces model can be converted to a pPCA model with isotropic noise in both the latent space and complementary space by subtracting the noise from the latent variable's covariance matrix), meaning that classification of points by the spaces will be the same. Our task is then to partition the rows of $\mathbf{O}'^{T}$ into a set of sites that are differentially methylated and therefore are modeled by a (*k*-1)-dimensional space (signal + noise) and a second set of sites that are not differentially methylated and are better modeled by a 0-D space (pure noise). This could be extended to model other signals, such as a disease status or a batch effect that correlates with another subset of sites.

***3.3: The distribution of noise at each site is approximately isotropic.*** Here, we show columns of our noise matrix $\mathbf{O}_j^{(\mathbf{n})} \sim \mathcal{N}_N(\mathbf{0}, \sigma_j^2 \cdot \mathrm{diag}(||\mathbf{R}_i||_2, ..., ||\mathbf{R}_i||_2, ..., ||\mathbf{R}_N||_2))$ and the degree of anisotropy is therefore bounded by the constraints $\sum_{k=1}^{K} R_{ik} = 1, R_{ik} \geq 0$. Each $O_{ij}^{(\mathbf{n})}$ is the sum of normal random variables, making it normally distributed. Specifically, each $O_{ij}^{(\mathbf{n})}$ is the weighted mean of $K$ independent samples $X'_{ij1}...X'_{ijK}$ from the same normal distribution.

First we consider our rows to understand where the anisotropy arises. Entries within a row $\mathbf{O}_i^{(\mathbf{n})}$ (values for different sites but the same individual) are uncorrelated but not independent. While all $X'_{ijk}$ are i.i.d. for a given $j$, the realizations of $\mathbf{R}$ for each individual are shared by entries within rows of $\mathbf{O^{(n)}}$, so there are systematic differences between dimensions (individuals) when we cluster on $\mathbf{O^{(n)}}^{\mathrm{T}}$. Consider a simple case examining site $j$ in individual A, who has uniform proportions of cell types, and individual B, whose blood is composed of only one cell type. Individual B's variance for noise at site $j$ is $\sigma_j^2$, while individual A's is the variance of the sample mean of $K$ realizations with variance $\sigma_j^2$: $\frac{\sigma_j^2}{K}$. Individuals with more imbalanced cell proportions will have greater magnitude for entries in their rows of the noise matrix $\mathbf{O^{(n)}}$ because $\mathrm{var}(O_{i1}^{(\mathbf{n})}), ..., \mathrm{var}(O_{ij}^{(\mathbf{n})}), ..., \mathrm{var}(O_{iJ}^{(\mathbf{n})})$ are not independent.

In contrast, entries within a column $\mathbf{O}_j^{(\mathbf{n})}$ (noise values for the same site across individuals), are independent as no information is shared. We can then treat a column as a realization of a multivariate normal distribution with a diagonal covariance matrix. $O_{ij}^{(\mathbf{n})}$ is the sum of random variables $R_{i1}X'_{ij1}...R_{iK}X'_{ijK}$. $\text{cov}(R_{ia}X'_{ija}, R_{ib}X'_{ijb}) = 0$ for $a \neq b$, because $R_{ik}$ and $X'_{ijk}$ are independent, and $\mathbb{E}[X'_{ijk}] = 0$. (Alternatively, one could observe that while $R_{ia}X'_{ija}$ and $R_{ib}X'_{ijb}$ are not independent— their magnitudes are related, because as $R_{ia}$ increases, $R_{ib}$ is more likely to be smaller— they are uncorrelated, as the sign of $X'_{ijk}$ is independent for each entry and it is symmetric about 0.) Thus, the variance for $O_{ij}^{(\mathbf{n})}$ is the sum of variances for the underlying $R_{i1}X'_{ij1}...R_{iK}X'_{ijK}$. Once the cell proportion vectors $\mathbf{R}_i$ have been realized, $\text{var}(O_{ij}^{(\mathbf{n})}) = \sum_{k=1}^{K} R_{ik}^2 \text{var}(X'_{ijk}) = \sigma_j^2 \sum_{k=1}^{K} R_{ik}^2$ because each $R_{ij}$ is fixed in $R_{i1}X'_{ij1}...R_{iK}X'_{ijK}$. This can be simplified to $\text{var}_{ij} = \sigma_j^2 ||\mathbf{R}_i||_2$, and because $R_{ik} \geq 1 \forall i, k$, we can bound $||\mathbf{R}_i||_2$ to be between 1 and $\frac{1}{k}$.

$k$-spaces clustering fits a model of isotropic Gaussian noise. Our consideration of entries within individual columns shows that the distribution of noise is approximately isotropic. While the M-step only requires that the principal components of variation correspond to the signal from cell proportion variation, assignment of points to spaces in the E-step could be affected if the variation in the scale of noise between individuals, $||\mathbf{R}_i||_2$, becomes extreme and noise is significantly non-isotropic. However, $||\mathbf{R}_i||_2$ is bounded between 1 and $\frac{1}{K}$, and $K$ is small ($K = 5$) in this study. This non-isotropic noise would be properly modeled by a factor model, but as $\alpha$ grows and the imbalance in cell proportions between individuals reduces, noise will be approximately isotropic. Furthermore, it is the degree of imbalances between individuals' cell proportions $\mathbf{R}_i$ that that matters for non-isotropy rather than the absolute differences between individuals' cell proportions, because that is what drives variation in $||\mathbf{R}_i||_2$. In practice, we do not need the unrealistic assumption that cell types are present in equal proportions (as in the symmetric Dirichlet distribution parameterized by $\alpha$)— only that cell proportions are imbalanced to approximately the same extent across individuals. Moving forward, we assume noise is approximately isotropic so that we can more easily study the effect of site-specific variances to produce non-Gaussian noise.

***3.4: Complementary space likelihood of each point $\mathbf{O}_j^{(\mathbf{n})}$ decreases monotonically with residual length and classification is biased depending on distance from origin.*** Under the assumption that noise is isotropic with respect to individuals ($c^2 = ||\mathbf{R}_1||_2 = ... = ||\mathbf{R}_N||_2$), $\mathbf{O}_j^{(\mathbf{n})} \sim \mathcal{N}_N(\mathbf{0}, \sigma_j^2 c^2 \text{I})$. Fig. S10 shows that we can approximate the empirical distribution of $\sigma_j$ fairly well with a Half-normal distribution. Combining equation (8) with a Half-normal prior on $\sigma_j$, we get the following distribution over columns: $f_{\mathbf{O}_j^{(\mathbf{n})}}(r) = \int_0^\infty \frac{1}{(2\pi)^{n/2}\sigma_j^n} e^{-\frac{1}{2}\frac{r^2}{\sigma_j^2}} \cdot \frac{2}{\sigma_* \sqrt{2\pi}} e^{-\frac{1}{2}\sigma_j^2/\sigma_*^2} d\sigma_j$, which monotonically decreases with $r$. Therefore, provided noise is approximately isotropic, it would seem appropriate to use a classification scheme that assigns a point to its nearest subspace. However, there is the caveat that assuming both spaces correctly pass through the origin, any point other than the origin will be closer to the $(k-1)$-dimensional space, so a latent distribution is needed. Thus, we set our noise parameter to be equal between the spaces to exploit the monotonicity rather than a pure, non probabilistic generalization of $k$-means that assigns points to the closest affine subspace.

To understand the impact of the latent distribution in the $(k-1)$-D space, we focus on the vast majority of non DMRs that are located generally close to the origin because they lack variance from cell type proportions. For assignment in $k$-spaces clustering to work on the not-linearly separable DMRs and non DMRs, likelihood for non DMRs given by the $(k-1)$-D space must drop sufficiently due to spreading of probability density over the latent dimensions by the latent space Gaussian versus the tighter isotropic noise Gaussian. This also implies DMRs with very low overall variance between cell types ($M_{j1} \approx M_{jk} \approx ... \approx M_{jK}$ and $\sigma_j$ is small) will be biased towards assignment to the 0-D space. If the EM procedure translates the spaces to be centered at the origin and oriented correctly, and density mixing proportions $\pi_i$ of the spaces are equal, the switch towards misclassification of points should occur at the intersection between the PDF of the $(k-1)$-D Gaussian of the latent space and the PDF for noise along those same dimensions given by the 0-D isotropic Gaussian. In contrast, non DMRs may be misclassified when the realized $\sigma_j$ is large by the same argument: beyond this boundary, the more spread out latent distribution provides a higher density in those dimensions. This could imply a limit on classification accuracy for $k$-spaces clustering as opposed to a specific model designed to detect DMRs.

We hypothesize this balance between spread out latent distribution density of the latent space and tighter isotropic density of the noise space is what produces the observed sudden failure of the model when $\alpha$ becomes large and $\sigma_j$ is bootstrapped from the heavier-tailed observed distribution, as curiously, sensitivity rather than specificity drops to 0 when the model is initialized with approximately correct spaces and allowed to converge to a local minima (Fig. S26b) (For this analysis, the model was initialized by separately fitting each space to the entire dataset, which provides no input information about the partitioning of sites into DMRs and non DMRs but does provide a reasonable starting point for the basis vectors of the $(k-1)$-D space). Of note, this is not the maximum likelihood solution as assigning all points to the $(k-1)$-D space (sensitivity = 1, specificity = 0) is a higher (or equal) likelihood model than assigning all points to the 0-D space. Removing sites with extreme variance (sites with $\sigma_j$ > 0.2, or the top 0.29%) led to the disappearance of this behavior (Fig. S26c).

**4: Note on merged qHCR image hues.** *k*-spaces may naively seem to be similar to counting the number of visually identifiable colors when the 4 channels of the image are overlaid. Indeed, examination of Fig. 4a in (9) shows distinct colors in different regions of the image, where pixel intensities across the 4 channels (*myod1*, *tpm3*, *her1*, *her7*) are mixed together as violet, blue, green, and red, respectively. However, there are two caveats to this. First, and more trivial, is that identifying hues (i.e. by taking a 3 channel qHCR image and assigning the three channels to RGB values) assumes that the lines pass through the origin. In such a case, this could be mathematically represented by hierarchical clustering using the cosine similarity index. However, more critically, even in the 4-channel case, depending on how pixel intensities across channels are mixed to produce colors, different anatomical regions may appear to be of the same hue as one another in one rendering of the data but of different hues in another version of the same image. Thus, even in the 4-dimensional case, our eyes already become unreliable.

**5: Note on overfitting.** A study reexamining ReFACTor's performance on the GALA II dataset pointed out that ReFACTor was trained and tested on the same data and was overfitting (10). We note that just as ReFACTor experienced large drops in $R^2$ when the linear regression was done with train-test partitions, so does *k*-spaces (Fig. S29) (10). However, the eigenvalues (0.266, 0.227, 0.134, 0.120) of *k*-spaces 4-D latent space (i.e. the latent $\sigma_1 ... \sigma_4$) suggest that the observed signal is largely coming from only be three detectable cell types, mirroring the ground truth cell type variation. The ground truth cell composition data in the GALA II dataset indicates that the two cell types demonstrating severe overfitting, monocytes and basophils, have interquartile ranges from 5.7 to 7.9% and 0.4 to 0.5% respectively, near or below the limit of detection in simulation (Fig. 3b, Table S1). In fact, *k*-spaces clustering with only a 2-D latent space rather than a 4-D one performs just as well (Fig. S30).

**6: *k*-spaces clustering identifies mixed cells in single-cell species-mixing experiments.** We also show that *k*-spaces is a useful approach for quantifying the separation in mixtures of cells from different species used to assess the fidelity of single-cell RNA-seq assays. Typically, such mixtures are studied heuristically via a "barnyard plot" comparing the number of unique molecular identifiers (UMIs) that map to the human genome to the number that map to the mouse genome for each barcode. While estimating the rate of doublets or multiplets can be done with manual thresholding, using *k*-spaces, we deconvolve these data by fitting two lines and a point, corresponding to human cells, mouse cells, and mixed cells, which include droplets contaminated by nearby lysed cells as well as doublets. Setting the noise parameter equal for all spaces produces an estimation of a mixed cell rate of 1.1% (mainly doublets), but it underestimates the rate of contamination (Fig. S23a). Allowing each space to have its own noise parameter yields a 6.7% rate of mixed cells, including both doublets and other contaminating processes, and an estimated standard deviation of 32 and 47 cross-species UMIs per droplet for pure human and pure mouse cells, respectively (Fig. S22a).

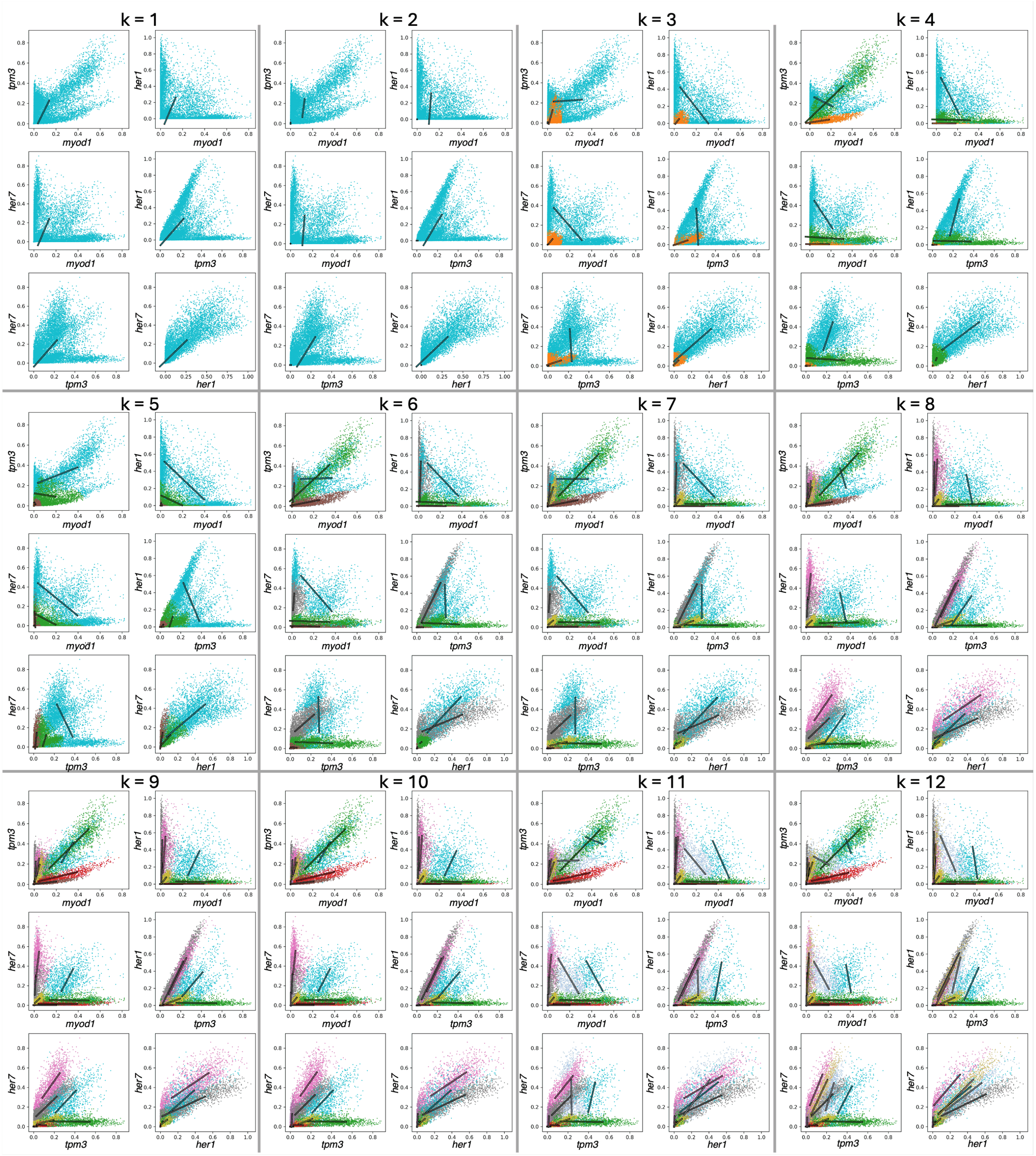


**Fig. S1. Pairwise voxel intensity scatter plots and projections of fitted lines using *k*-spaces with *k* = $\{1, 2, \ldots, 12\}$ on the qHCR zebrafish somitogenesis image of Fig. 2.**

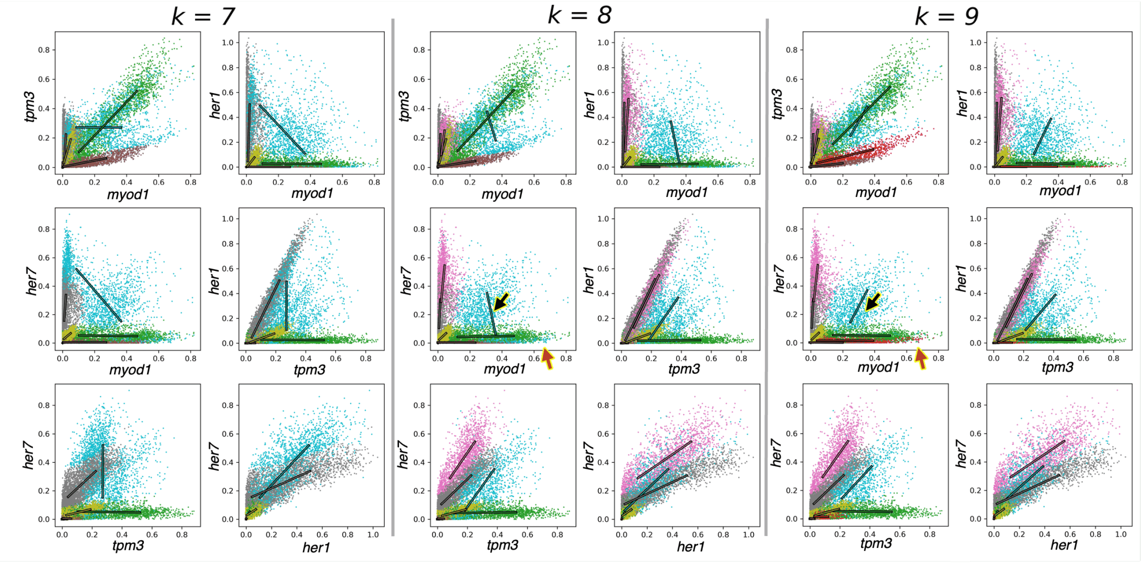


**Fig. S2. Pairwise voxel intensity scatter plots and projections of fitted lines for *k* = $\{7, 8, 9\}$ to aid in model selection.** The six pairwise plots for full data, colored by cluster assignment, along with projections of fitted lines as described in Fig. 2b. Arrowheads indicate evidence of underfitting in $k = 8$.

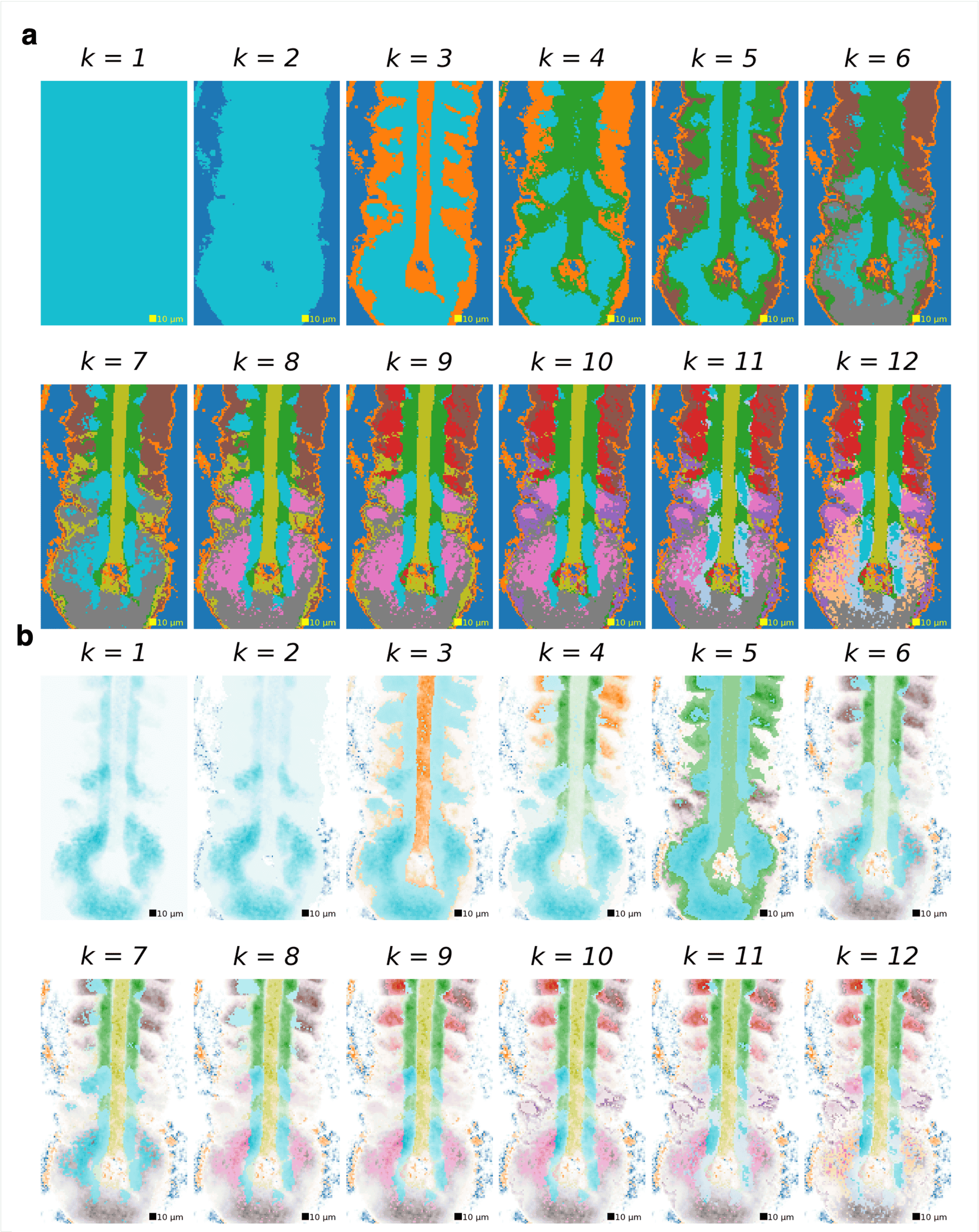


**Fig. S3. Automatic segmentation of the qHCR zebrafish somitogenesis image of Fig. 2 using *k*-spaces with *k* = $\{1, 2, \ldots, 12\}$.** Each panel is the highest likelihood result out of 100 *k*-spaces runs. *k*-spaces runs for a given value of $k$ do not build off of the $k-1$ solution. **a,** Each voxel is shaded by its *k*-spaces cluster index.**b,** *k*-spaces clustering with color saturation proportional to latent variable values as described in Methods.

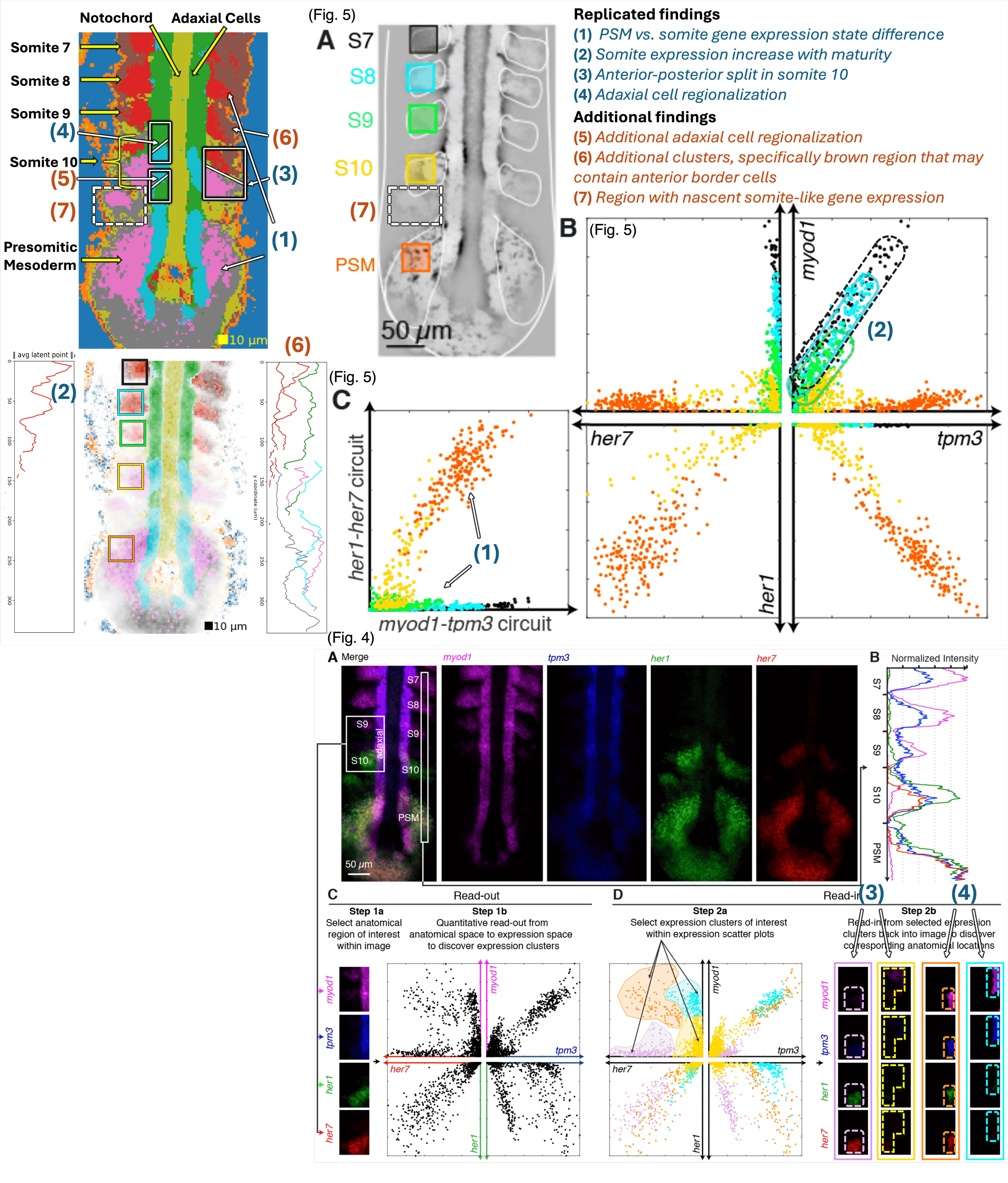


**Fig. S4. qHCR image analysis: annotated comparison to Figs. 4 and 5 of Trivedi et al., 2018 (9).** Fig. 4 of Trivedi et al., 2018 demonstrates read-out/read-in: read-out from one ROI encompassing somites S10 and S9 as well as adjacent adaxial cells to pairwise expression scatter plots (panel C), read-in from manually selected expression clusters back into the anatomical context of the image (panel D). Fig. 5 of Trivedi et al., 2018 demonstrates read-out for five ROIs (PSM, S10, S9, S8, S7) with voxel intensities from panel A displayed in the pairwise expression scatter plots of panel B, and the circuit scatter plot of panel C; note that image cropping differs slightly from the present work. Replicated and additional findings are annotated with colored numbers. **Replicated findings: (1)** *k*-spaces identifies pink and gray programs in the presomitic mesoderm (PSM) vs red and brown programs in the maturing somites (S9, S8, S7); in Fig. 5C, the PSM co-expresses the *her1-her7* circuit and the *myod1-tpm3* circuit while the maturing somites (S9, S8, S7) only express the *myod1-tpm3* circuit. **(2)** *k*-spaces identifies that the red program strength increases as somites mature (S9→S8→S7), as shown by increasing saturation and vertical profile of the red latent variable; in Fig. 5B this can be seen in the increasing amplitude and constant slope of the green, cyan, and black expression clusters.

**Fig. S4 continued.** **(3)** *k*-spaces identifies anterior/posterior regionalization of the youngest somite (S10) with the posterior region featuring the pink program found in the adjacent PSM and the anterior region featuring the red program found in the adjacent maturing somites (S9, S8, S7); Fig. 4D displays similar regionalization of S10 (anterior: yellow program; posterior: lavender program). **(4)** *k*-spaces identifies corresponding regionalization of the adaxial cells adjacent to the nascent somite (anterior: green program; posterior: cyan program); Fig. 4D displays corresponding regionalization (anterior: cyan program; posterior: orange program). **Additional findings:** **(5)** *k*-spaces identifies additional segmentation of the adaxial cells between the nascent somite (S10) and the PSM (alternating between the green and cyan programs); Fig. 4 did not include this region in the ROI so this spatial oscillation of adaxial circuit states was not identified. **(6)** *k*-spaces identifies additional expression clusters corresponding to possible detection of distinct cell types within the maturing somites (red vs brown programs) and the PSM (pink vs gray programs); Fig. 4 did not identify these subdivisions of the maturing somites and the PSM using manual cluster selection. **(7)** *k*-spaces detected a structure between the PSM and the left nascent somite (S10) that deviates from bilateral symmetry and features the same pink program featured in the PSM and the posterior region of S10; Figs. 4 and 5 did not include this region in the manually selected ROIs. Figs. 4 and 5 adapted with permission of N.P., senior author, from Trivedi et al., *Development*, 145, dev156869. Copyright 2018 The Company of Biologists.

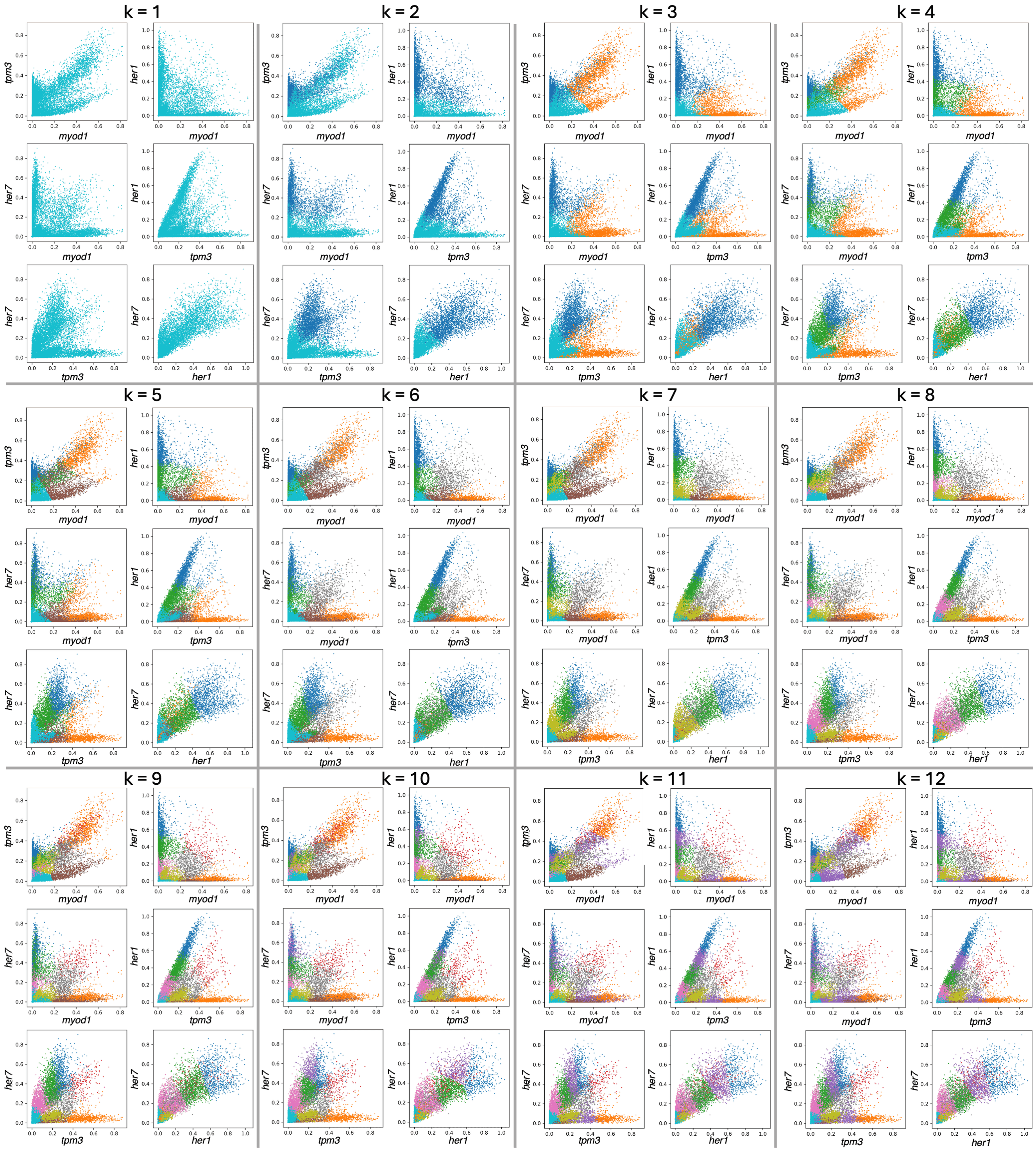


**Fig. S5. Pairwise voxel scatter plots colored by cluster for *k*-means with *k* = $\{1, 2, \ldots, 12\}$.**

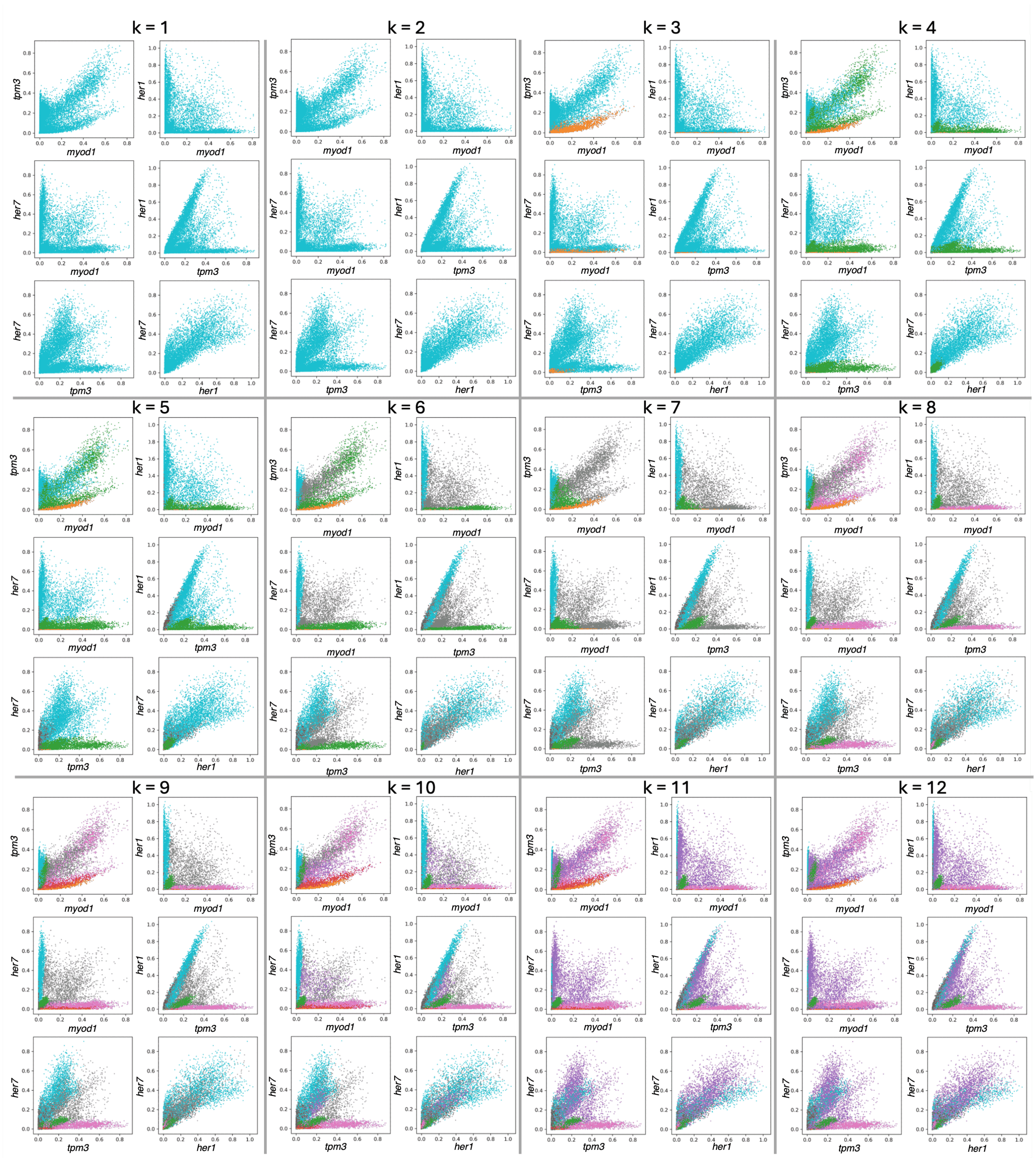


**Fig. S6. Pairwise voxel scatter plots colored by cluster for GMMs with *k* = $\{1, 2, \ldots, 12\}$.**

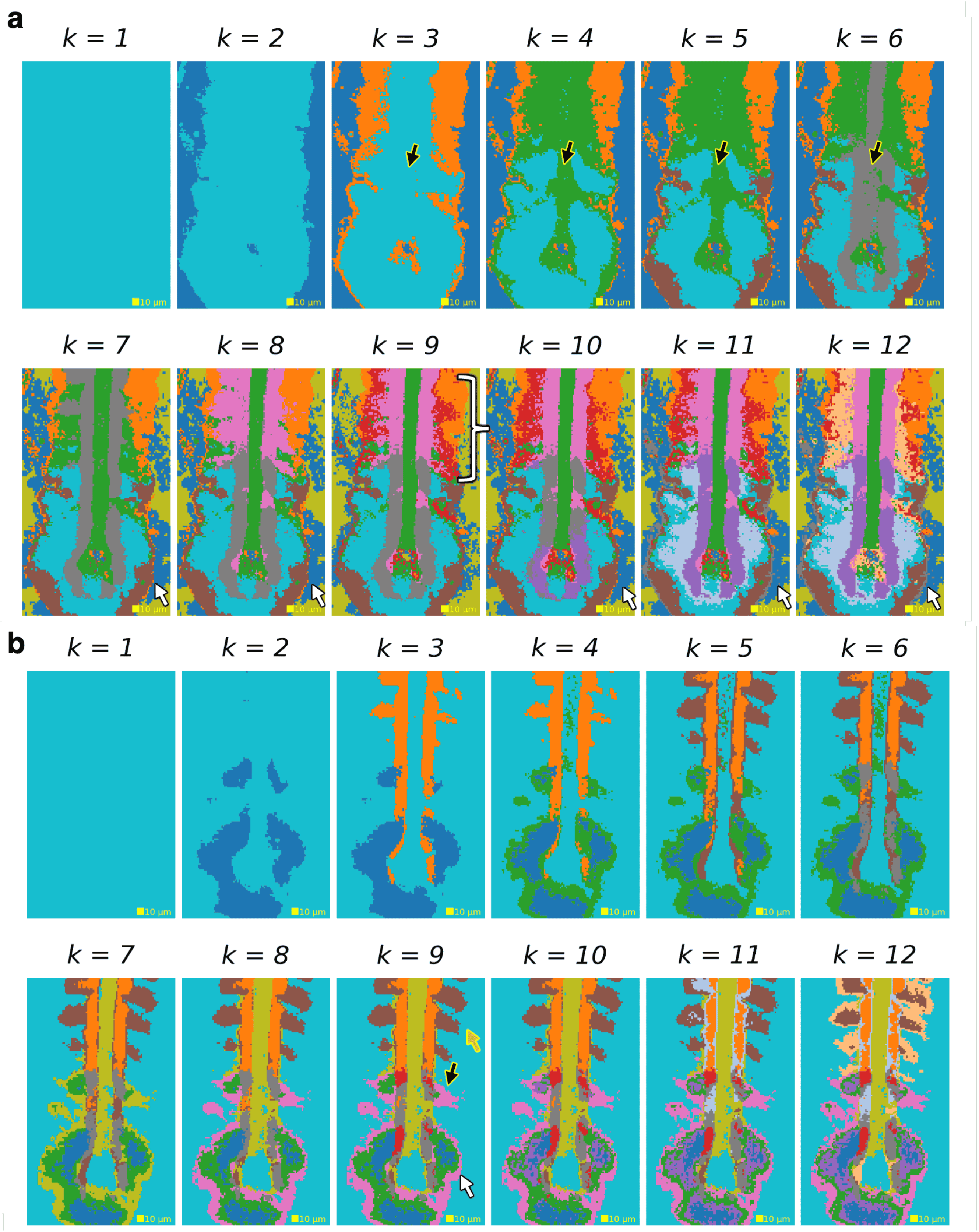


**Fig. S7. Comparison of Gaussian Mixture Models and *k*-means performance on zebrafish embryo data (comparison for Fig. S3a using *k*-spaces).** Each image is the highest likelihood result out of 100 *k*-spaces runs. *k*-spaces runs for a given value of $k$ do not build off of the $k-1$ solution. **a,** GMM clustering. Black arrowheads indicate undetected notochord until $k$ = 7, white arrowheads indicate over-segmentation of background space, and white braces indicate somite region which is poorly segmented across most values of $k$. **b,** *k*-means clustering. Black arrowhead indicates misclassified anterior portion of somite 10 as background, gold arrowhead indicates misclassification of cells on edge of all somites as background, and white arrowhead marks concentric segmentation of presomitic mesoderm likely because the 0-dimensionality of clusters in *k*-means underfits linear clusters.

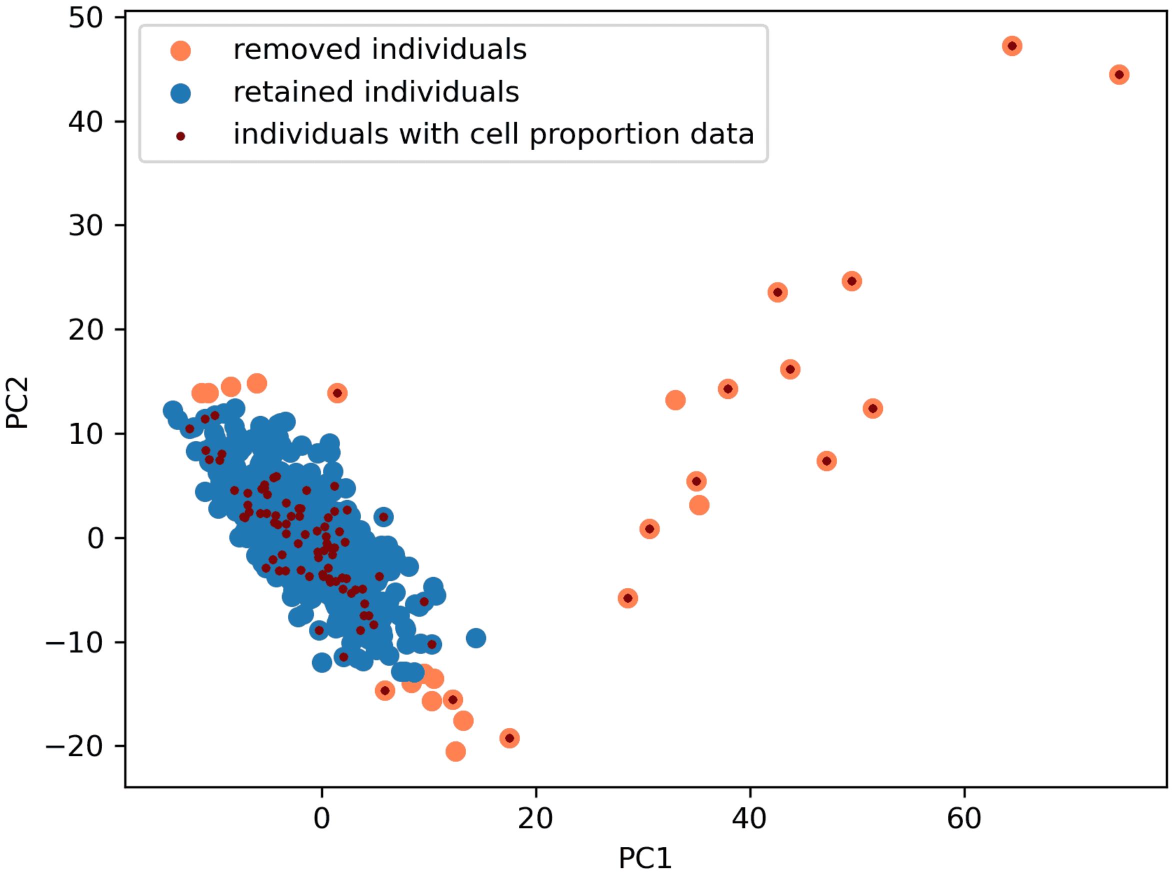


**Fig. S8. Result of preprocessing performed as described in the ReFACTor study (11)** . Individuals more than two standard deviations away from the mean on the first two principal components are shown in orange.

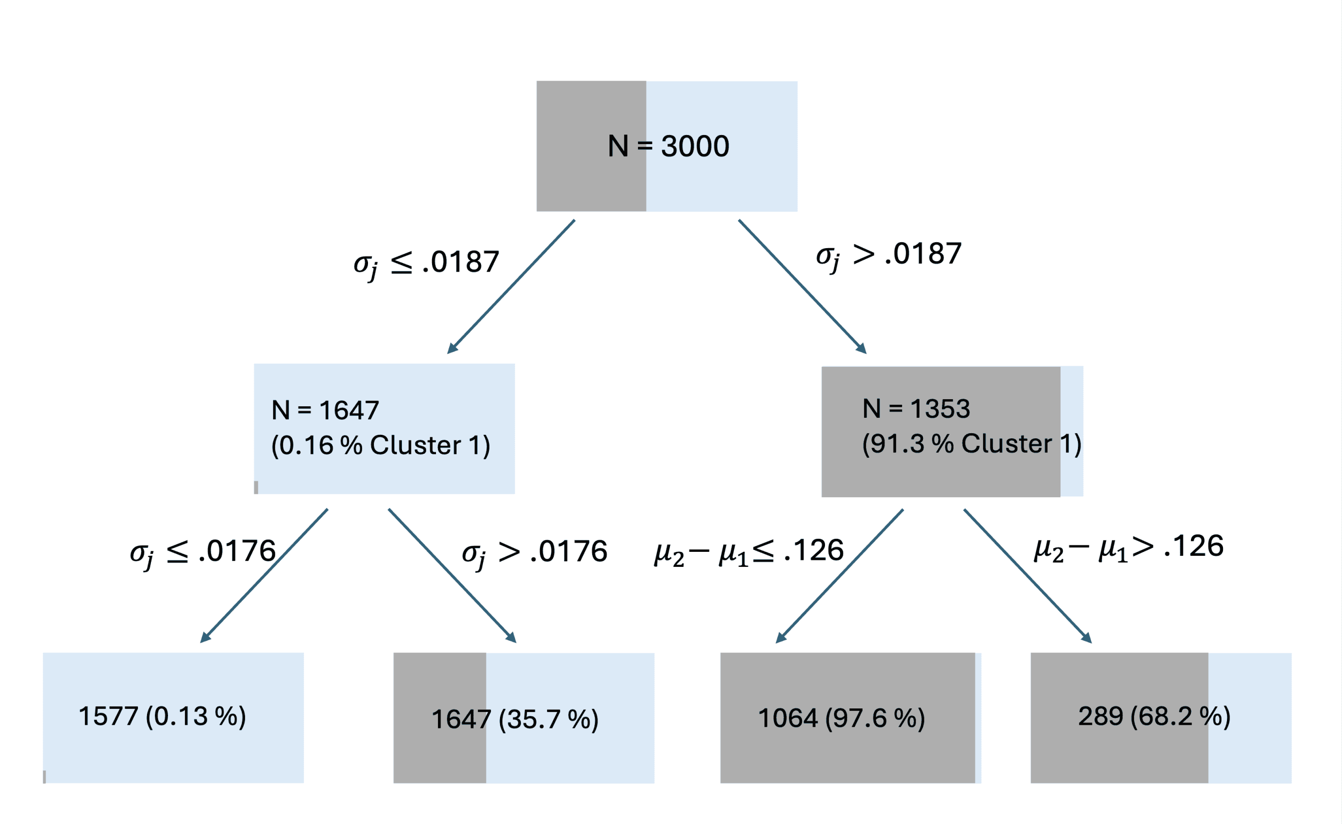


**Fig. S9. Decision tree trained to predict *k*-spaces DMR calls using ground truth difference in $\mu_j$ between cell types and ground truth site specific noise $\sigma_j$ on 3,000 ground truth DMRs in a set of 10,000 total simulated sites.** Simulation was done with the same process described in Methods with only 2 cell types, 500 individuals from a Dirichlet distribution with $\alpha = 1$, 10,000 total sites, 30% DMRs, no added technical noise, and $\sigma_j$ drawn from a half normal with a standard deviation of $0.025$. $\tau = 0.07$ as in the simulation in the main text. Nodes are shaded gray according to the percentage of simulated methylation sites in that node assigned by *k*-spaces to cluster 1 (called DMRs). The high prediction accuracy at the second level of the tree using only $\sigma_j$ reveals that poor performance of *k*-spaces in calling DMRs with unconstrained noise parameters was due to fitting the variation in site-specific variance rather than variation in cell type proportions.

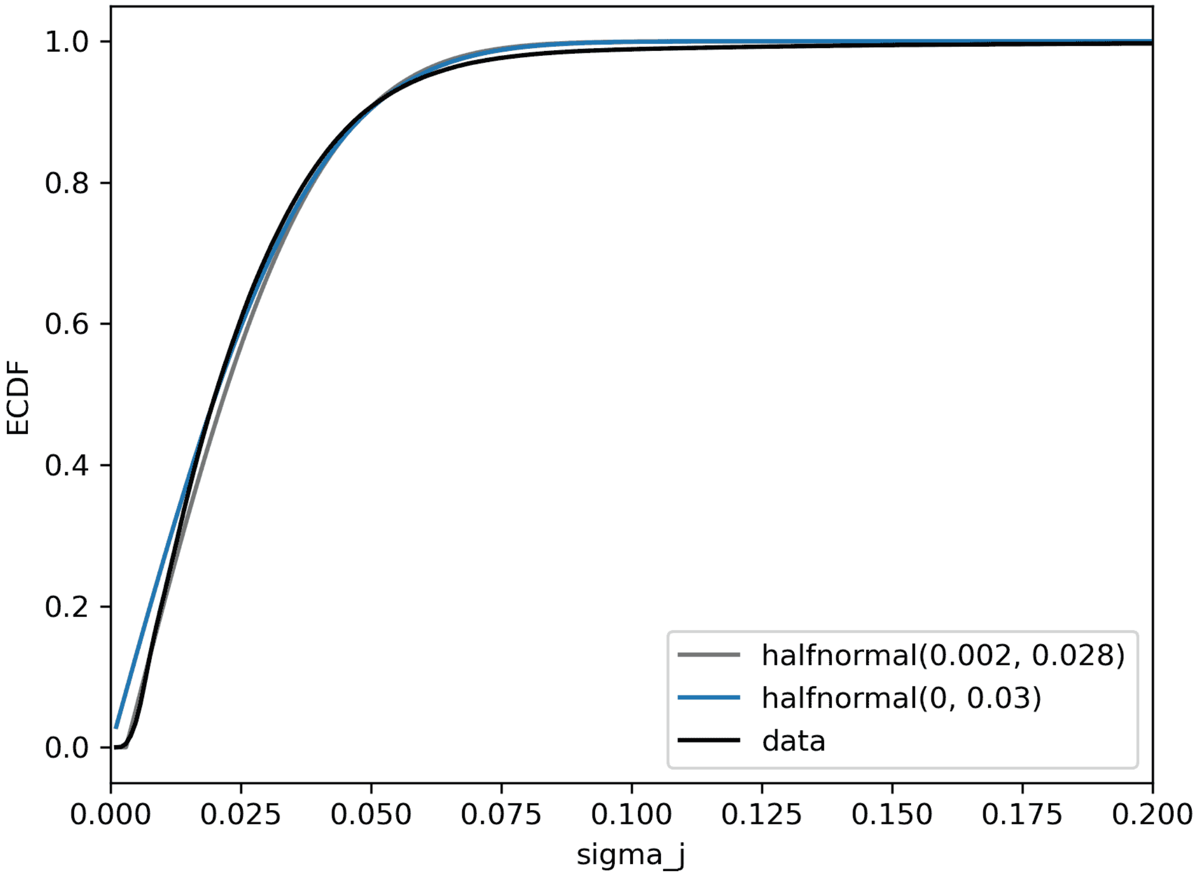


**Fig. S10. The empirical distribution of site-specific variances in (8) is well approximated by a Half-normal distribution**. We use location = 0,$\sigma = 0.03$ in our simulation, as a nonzero lower bound seems unrealistic, but location = $0.002$, $\sigma = 0.0028$ is shown as well. While the Half-normal centered at 0 more noticeably underestimates the empirical distribution of sample variances for the lowest 20% of sites, it is the slight discrepancy at the tail that is responsible for the lower robustness to large $\alpha$ in the simulation when using the empirical distribution of variances instead of a Half-normal. Removing the 0.29% of sites with $\widehat{\sigma_j} > 0.2$ restores the robustness to sudden failure on $\alpha$ up to at least $100$ (Fig. S26).

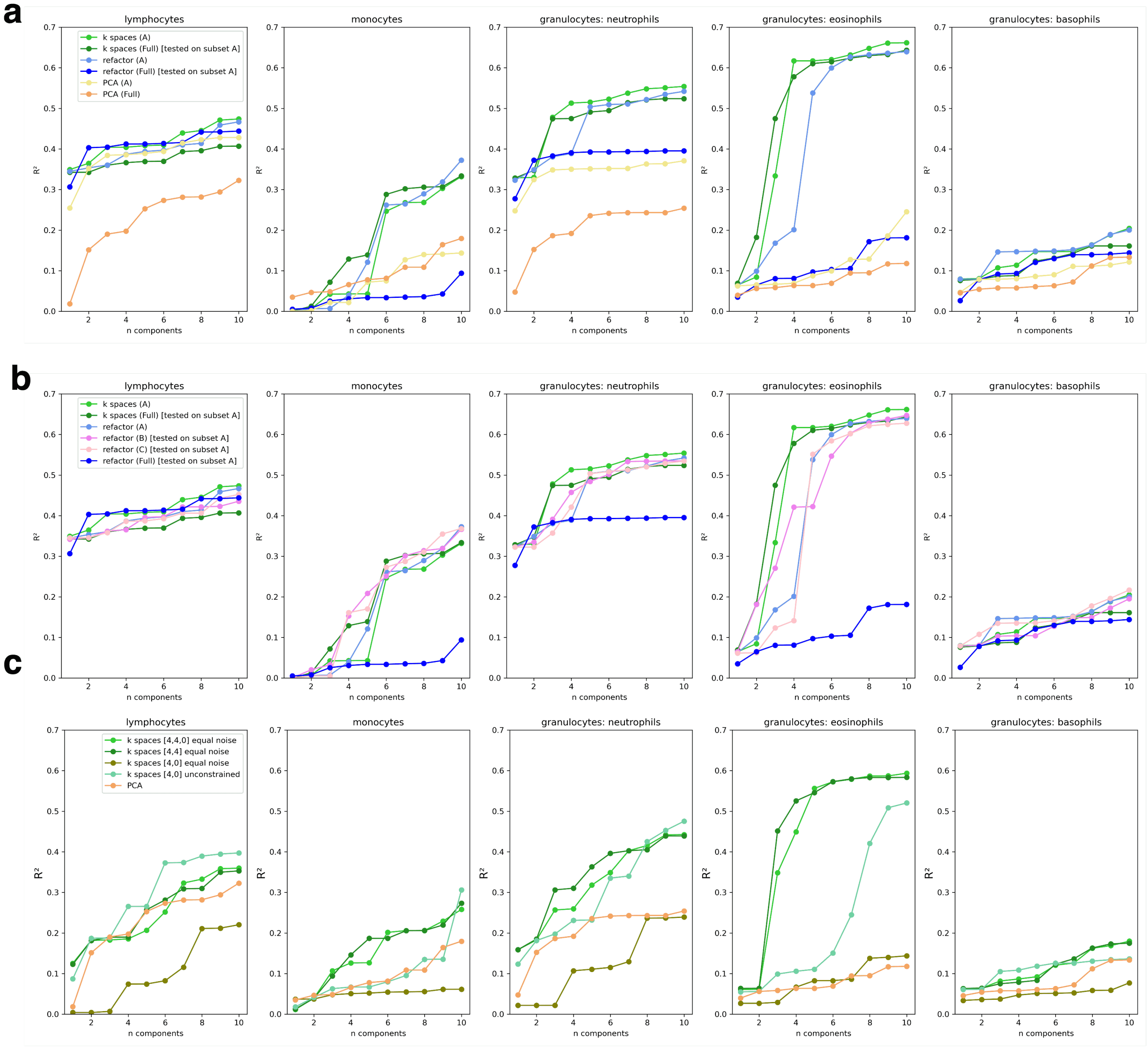


**Fig. S11. *k*-spaces clustering can perform better than ReFACTor on the full GALA II dataset. a,** *k*-spaces retains similar performance between the preprocessed and the full data (the top 500 closest sites by angle are selected for the preprocessed data in addition to thresholding by the noise of the 4-D structured noise space) while ReFACTor does not. Note that while PCA on the full data and PCA on the preprocessed data are quite different in lymphocytes and neutrophils, ReFACTor has the biggest changes in performance on monocytes and eosinophils, where PCA has no performance change. **b,** Eliminating either preprocessing step (B and C) does not result in significant difference, while eliminating both (Full) leads to failure of ReFACTor. **c,** *k*-spaces clustering with only one 4-D and one 0-D space performs poorly as well but can be corrected with the addition of another 4-D space to model structured noise, as shown in (a), or with replacement of the 0-D space with another 4-D space. Incidentally, *k*-spaces clustering with only one 4-D and one 0-D space but without the shared noise parameter performs well.

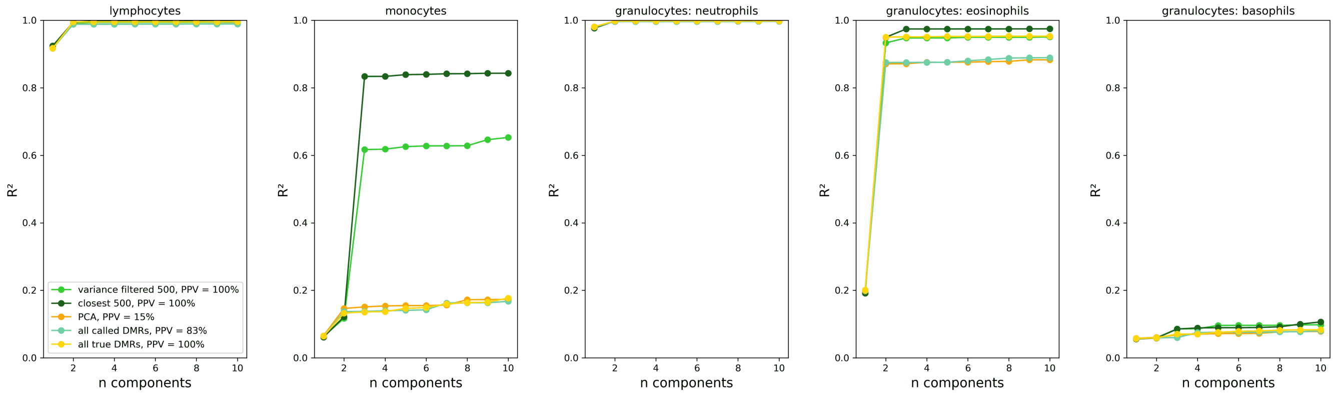


**Fig. S12. Good DMR detection and good cell type proportion are not equivalent in simulation**. Simulated data is the same used to generate 3b and bootstrapped cell type proportions from GALA II (12) and cell type profiles and site specific variances from (8) as described in Methods. PCA and regression was performed as in 3b. Despite good performance in simulation (78% sensitivity, 96% specificity, and 83% positive predictive value (PPV), regression using PCA on all called DMRs fails (teal) and is no better than PCA on all data (orange). Regression analysis on the ground truth DMRs (gold) fails to outperform standard PCA as well. Post-clustering filtering strategies (light green, dark green) improve performance.

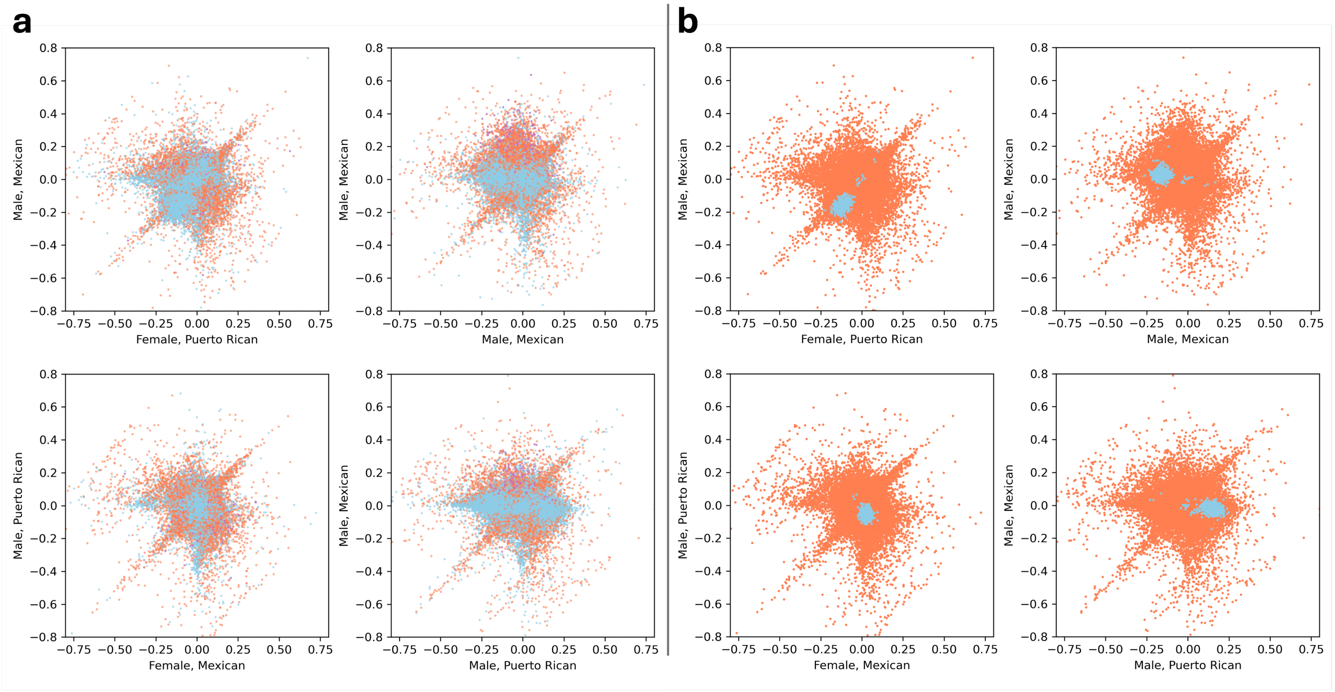


**Fig. S13. Called DMRs and selected informative DMRs shown in four pairs of individuals from the full GALA II dataset. a,** All sites assigned to the first 4-D latent space (capturing structured noise with respect to the cell proportion problem) are shown in orange. All sites assigned to the second 4-D latent space (containing DMRs) are shown in blue. Remaining sites clustered onto 0-D space are purple. Noise is evidently not isotropic, and a streak persists independent of apparent cell proportion-driven covariation in DMRs. **b,** 500 DMRs (blue) selected from the 4-D DMR space on the basis of having a variance greater than that of the structured noise 4-D space capturing sites perturbed by random noise. All other sites are colored orange.

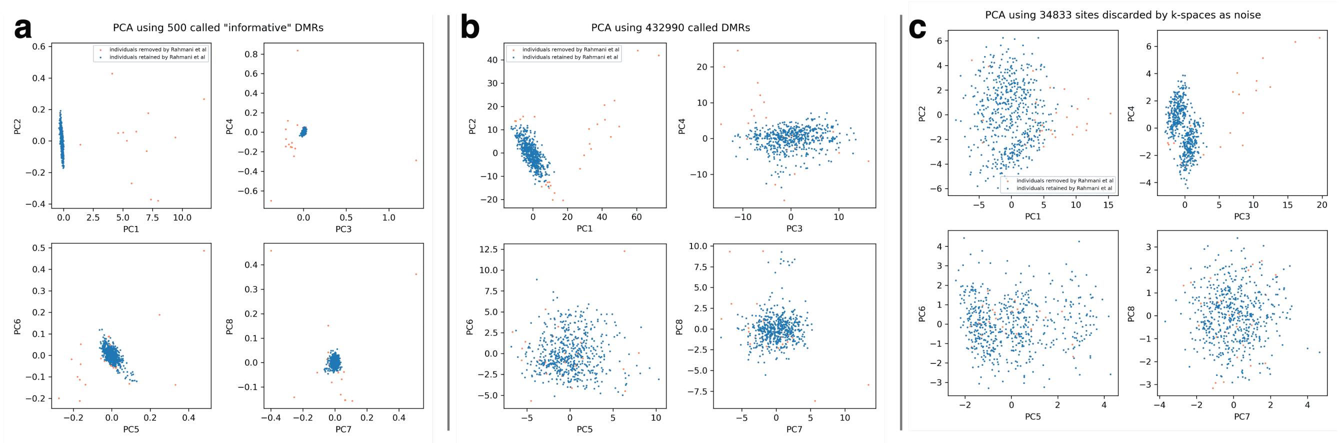


**Fig. S14. Structure of 'signal' and 'structured noise' discarded by *k*-spaces [4,4,0] on the GALA II data.** Orange points are those removed in (11) prior to applying ReFACTor. **a,** Top 8 PCs of the data using the selected "informative" DMRs. **b,** Top 8 PCs of the data using all sites captured by the first 4-D subspace (all called DMRs). **c,** Top 8 PCs of the data using all sites captured by the second 4-D subspace intended to capture and discard structured noise.

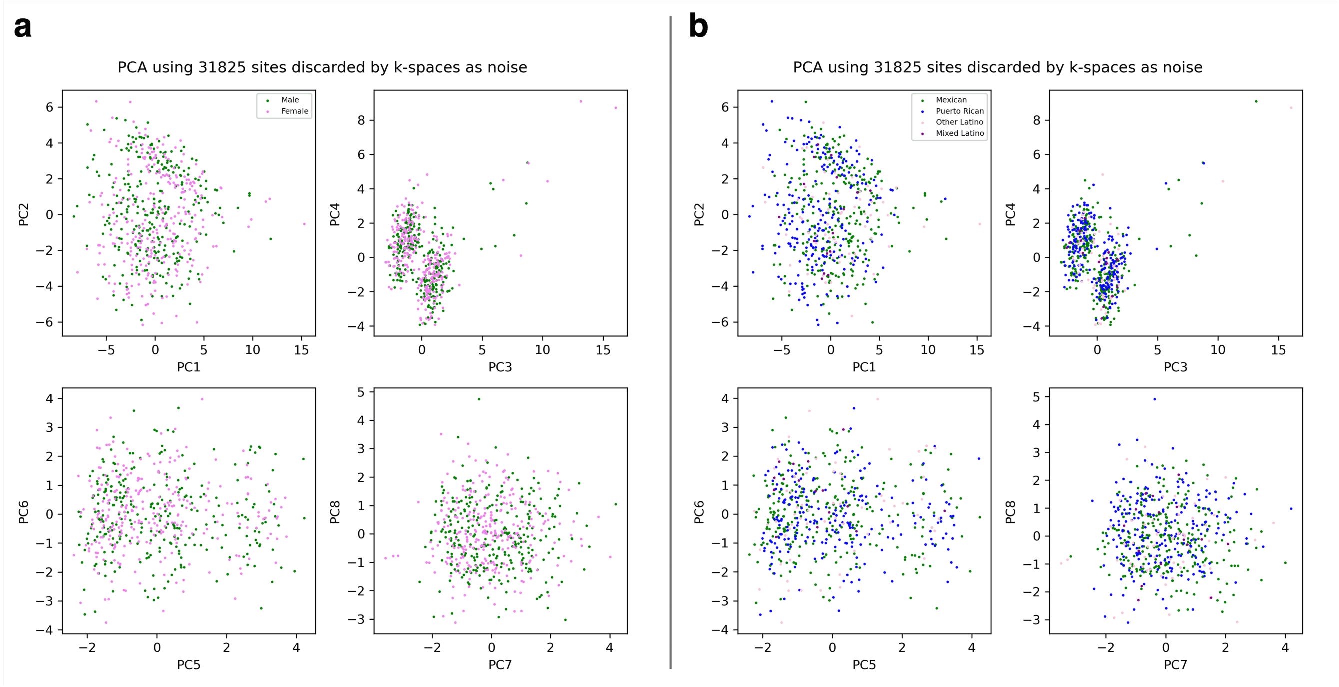


**Fig. S15. 'Structured noise' discarded by *k*-spaces in GALA II does not appear to correlate with available metadata.** PCA of samples using the 31825 sites assigned to the 4-D 'noise' subspace when clustering with two 4-D subspaces and a 0-D subspace.

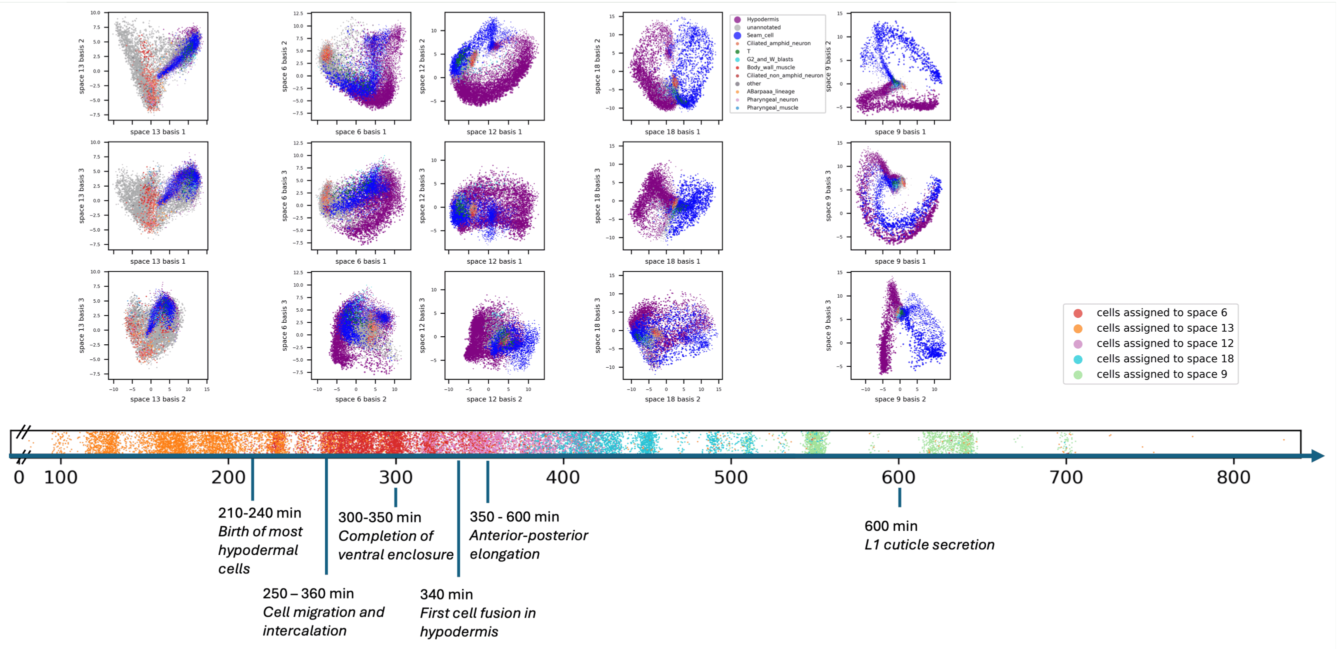


**Fig. S16. Clusters 6,9,12,13, and 18 in *C. elegans* embryogenesis (13) visualized with cell type annotations.** Neurons, hypodermis, and seam cells are all ectoderm derived lineages. Space 13 shows the divergence from neurons early in development, which is is depicted as a break in the orange UMAP cluster in Fig. 4b.

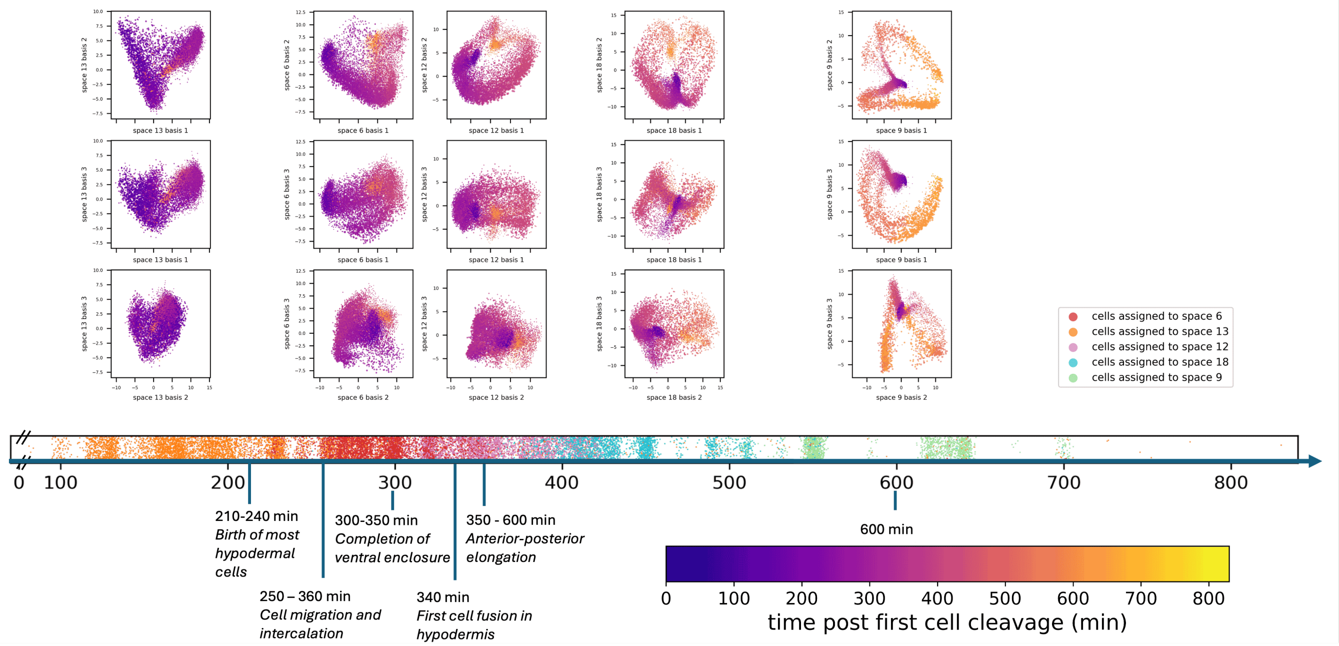


**Fig. S17. Clusters 6,9,12,13, and 18 in *C. elegans* embryogenesis (13) visualized with embryo time.** Early and late cells can be compared within the subspaces to determine if cells return to the same state as their progenitors with respect to the gene expression programs represented in a given axis or if there is continuous progression with maturity.

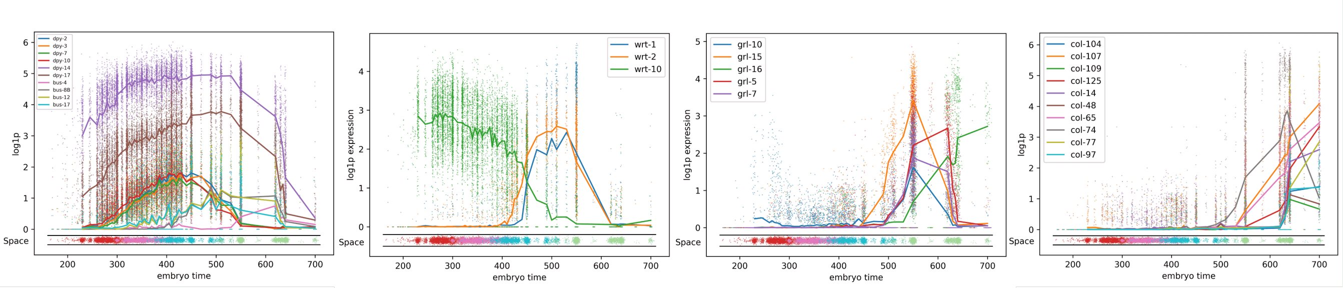


**Fig. S18. Selected gene families represented in *k*-spaces basis vectors**. While these genes typically strongly influenced the basis vectors (i.e., appeared among the top ranked genes by coefficients) in only one to two spaces, expression levels of the genes can subsequently be tracked across all cells to show when they are expressed. Cells from the orange cluster (cluster 13) are not included as that cluster includes neuronal progenitors.

| | Embryonic time (min) $10^{th}$-$90^{th}$ percentile | Enrichment analysis interpretation | GO Biological Process | GO Molecular Function | GO Cellular Component | Notes |
|---|---|---|---|---|---|---|
| 3-Space 13 | 130 – 260 | Events: end of rapid proliferation and birth of most hypodermal cells | | | | |
| Basis 1 | | | none | none | none | |
| Basis 2 | | | • Negative regulation of DNA recombination<br>• Nucleosome assembly<br>• Protein refolding<br>• Chromosome condensation<br>• Translation elongation | • Nucleosomal DNA binding<br>• Translation elongation factor activity<br>• Structural constituent of chromatin | • nucleosome | |
| 3-Space 6 | 260 - 330 | Events: cell migration and intercalation | | | | |
| Basis 1 | | Remodeling of ECM (lipocalin family) | • Tetrahydrobiopterin biosynthetic process<br>• cuticle development involved in collagen and cuticulin based molting cycle<br>• post-embryonic body morphogenesis | • structural constituent of collagen and cuticulin-based cuticle<br>• structural constituent of chromatin | • nucleosome<br>• gap junction<br>• collagen trimer<br>• extracellular region | Lots of dpy genes |
| 3-Space 12 | 320 – 410 | Evens: end of intercalation, completion of ventral closure, first cell fusion in hypodermis, elongation | | | | |
| Basis 1 | | molt | • notch signaling pathway<br>• molting cycle, collagen and cuticulin-based cuticle | • notch binding | • ER lumen<br>• extracellular region | Warthog, some dpy genes, nhr |
| Basis 2 | | | • lysine biosynthetic process via aminoadipic acid<br>• tetrahydrobiopterin biosynthetic process<br>• lipid transport<br>• carboxylic acid catabolic process | none | none | Nhr-127, nhr-270, lipocalin family |
| 3-Space 18 | 400 - 500 | Events: cell fusions in hypodermis, elongation | | | | |
| Basis 1 | | some cuticle genes | none | none | • extracellular matrix/external encapsulating structure | Cut-2, cut-5, nhr-270 |
| Basis 2 | | some cuticle genes | none | none | • cytosolic large ribosomal subunit | |
| 3-Space 9 | 550 – 640 | Events: cuticle secretion | | | | |
| Basis 1 | | Cuticle synthesis | • Unclassified<br>• *depleted for "cellular process" and "regulation of cellular process"* | • Structural component of cuticle<br>• *Depleted for "binding"* | • collagen trimer<br>• extracellular matrix<br>• Unclassified<br>• *depleted for "cytoplasm" and "nucleus"* | Grl family, Wrt family, lots of collagen |
| Basis 2 | | Cuticle synthesis | none | • structural component of cuticle<br>• glycosyltransferase activity<br>• *depleted for "binding"* | • extracellular matrix<br>• collagen trimer<br>• extracellular region | Turns off in seam cell arm |
| Basis 3 | | Cuticle synthesis | • *depleted for "cellular process"* | • structural component of cuticle,<br>• *depleted for "binding"* | • collagen containing ECM<br>• collagen trimer<br>• extracellular region<br>• *depleted for "cytoplasm" and "nucleus"* | on/off in both arms<br>Grl inverted sign with respect to pc1 |

**Fig. S19. Enrichment analysis results on genes with coefficients greater than $0.05$ or less than $-0.05$ in basis vectors from *k*-spaces.**

| 3-Space 13 B1 | 3-Space 13 B2 | 3-Space 6 B1 | 3-Space 12 B1 | 3-Space 12 B2 | 3-Space 18 B1 | 3-Space 18 B2 | 3-Space 9 B1 | 3-Space 9 B2 | 3-Space 9 B3 |
|---|---|---|---|---|---|---|---|---|---|
| 130-260 min | 130-260 min | 260-330 min | 320-410 min | 320-410 min | 400- 500 min | 400- 500 min | 550 – 640 min | 550 – 640 min | 550 – 640 min |
| | | | interpretation | | | | cuticle secretion | cuticle secretion | cuticle secretion |
| May be a bifurcation. Increases rapidly but in two directions | Decreases with maturation | Increases with maturation | Off/on for hypodermis, increases slightly for seam | Increases for hypodermis | Bifurcation, increases in seam cells vs hypodermis | Increases | Increase with maturation | Turning off in seam cells | On/off |
| Overshoots in one arm and then decreases in cells on loop. | | Decreases in late stage | | | | Decreases at later time points | decrease in early time points | Turned on from 400-500 min | Final state is same as early cells |
| - B0281.5<br>- C08F1.6<br>- C50F4.6<br>- F14B6.3<br>- F39F10.4<br>F53B3.5<br>F54D5.7<br>- F59C12.4<br>- K02B12.2<br>- T05D4.2<br>- T25E12.6<br>W04A8.4<br>Y110A2AL.4<br>Y65A5A.1<br>Y7A9D.1<br>act-4<br>- aldo-2<br>- ant-1.1<br>- cec-8<br>cey-1<br>cht-1<br>clec-196<br>cpg-20<br>cpg-24<br>crt-1<br>- cyb-3<br>dpy-14<br>- dsl-2<br>dsl-3<br>- eef-1A.1<br>eef-1A.2<br>- eef-1B.1<br>- eef-1G<br>enpl-1<br>erd-2.1<br>fasn-1<br>- fbxb-37<br>- fbxb-41<br>- fbxb-84<br>- fbxb-91<br>gmap-1<br>gst-20<br>- hil-3<br>- hil-4<br>idh-1<br>igcm-3<br>lbp-1<br>lbp-3<br>lpr-3<br>- mcm-7<br>mps-2<br>noah-1<br>- pcn-1<br>pdi-2<br>- plag-15<br>pmt-2<br>- rbm-3.1<br>ret-1<br>rml-3<br>- rsp-1<br>sec-61.A<br>- skr-7<br>sqt-3<br>srlf-1<br>sto-1<br>sym-1<br>- tba-1<br>- tbb-2<br>- top-2<br>trap-2<br>txdc-12.1<br>wrt-10 | B0281.5<br>C01G6.3<br>- C08F1.10<br>C35E7.5<br>F14B6.3<br>F39F10.4<br>- F44E5.4<br>- F44E5.5<br>F59C12.4<br>R06C1.4<br>R11A5.3<br>T05D4.2<br>- W04A8.4<br>- Y71F9AL.7<br>- Y75B8A.18<br>Y82E9BR.3<br>- ZK973.11<br>ahcy-1<br>aldo-2<br>- asp-4<br>atp-6<br>btb-6<br>cht-1<br>clec-196<br>- cnt-2<br>ctc-3<br>cyb-3<br>dpy-14<br>dsl-2<br>dsl-3<br>eef-1A.1<br>eef-1B.1<br>eef-1G<br>epg-2<br>- fbxb-66<br>gmap-1<br>- hil-2<br>- hil-3<br>hil-4<br>- his-24<br>- hmg-1.1<br>- hsp-16.2<br>- hsp-16.41<br>- hsp-16.48<br>hsp-3<br>- knl-2<br>lbp-3<br>- mix-1<br>- nurf-1<br>pab-1<br>pdi-2<br>phdh-1<br>plag-15<br>pmt-2<br>rbm-3.1<br>skr-7<br>sqt-3<br>srlf-1<br>- sto-1<br>sym-1<br>tba-1<br>tbb-2<br>tct-1<br>- tir-1<br>ttr-50<br>wrt-10 | - C01G6.3<br>C26B9.3<br>C29E4.15<br>- C35E7.5<br>F48D6.4<br>F57B9.1<br>K02E10.4<br>K09G1.1<br>- T04D3.5<br>- T05H4.11<br>- W04A8.4<br>- Y37E3.30<br>Y41D4B.6<br>- Y65A5A.1<br>- Y7A9D.1<br>- act-4<br>- ant-1.1<br>aqp-2<br>- atz-1<br>calu-1<br>- cav-1<br>ceeh-1<br>- chdp-1<br>- cht-1<br>- clec-266<br>col-121<br>col-76<br>cpg-20<br>- cpi-2<br>- crt-1<br>cutl-2<br>cyn-6<br>dao-2<br>dpy-10<br>dpy-14<br>dpy-17<br>dpy-2<br>dpy-3<br>dpy-7<br>dsl-6<br>- enpl-1<br>- epg-2<br>fkb-4<br>fkb-7<br>hch-1<br>- hil-2<br>- his-24<br>- his-72<br>- hmg-1.1<br>- hrpa-1<br>- htz-1<br>inx-12<br>inx-13<br>- inx-3<br>lam-3<br>- lbp-1<br>- lbp-3<br>- lgg-1<br>- lpr-3<br>lron-8<br>mlt-11<br>mlt-8<br>- nbet-1<br>nlp-31<br>noah-1<br>noah-2<br>nsy-4<br>- pab-1<br>- pab-2<br>pcbd-1<br>pck-2<br>pgph-3<br>- pmt-2<br>qdpr-1<br>- rbm-3.2<br>sqt-3<br>- sto-1<br>- ttr-50<br>- ubq-1<br>ugt-60 | B0205.4<br>B0365.9<br>B0393.5<br>C06C3.4<br>C06E7.88<br>C26B9.3<br>C26F1.1<br>- C47E12.13<br>C48E7.1<br>C52G5.2<br>C53A5.2<br>F11E6.9<br>F15B9.8<br>F26E4.3<br>F31C3.6<br>- F33H2.8<br>F44E2.4<br>F46F11.7<br>F46G11.6<br>- F54D5.7<br>- F55C9.3<br>- F55C9.5<br>F57B9.1<br>K02E10.4<br>K08B12.1<br>M03F4.6<br>T01D1.8<br>- T04G9.4<br>T05H10.3<br>T19C4.1<br>- W04H10.6<br>Y102A11A.5<br>Y110A2AL.4<br>Y23H5B.8<br>- Y37E3.30<br>Y41D4B.6<br>Y45F10B.59<br>Y58A7A.2<br>- Y65A5A.1<br>ZC449.2<br>acn-1<br>aldo-2<br>- asns-2<br>atf-8<br>best-14<br>- clec-266<br>clec-78<br>col-121<br>- cpg-20<br>- cpg-24<br>- crt-1<br>- cutl-2<br>dao-2<br>dpy-2<br>dpy-3<br>dpy-7<br>- dsl-3<br>dsl-6<br>- enpl-1<br>far-3<br>- fasn-1<br>- fbn-1<br>- gln-3<br>- gst-20<br>- hch-1<br>hgo-1<br>hhat-1<br>hil-7<br>hsp-25<br>- hsp-3<br>- idh-1<br>ifa-3<br>- lbp-1<br>lpr-1<br>lpr-3<br>lpr-4<br>lpr-5<br>lpr-6<br>lpr-7<br>nab-1<br>- nhr-127<br>nlp-33<br>pcbd-1<br>- pdi-6<br>pgph-3<br>- pmt-2<br>ptr-25<br>- sams-1<br>spp-13<br>srm-1<br>- sto-1<br>- sym-1<br>tag-273<br>ttr-2<br>ttr-3<br>- txdc-12.1<br>unc-52<br>- wrt-10 | C35A5.11<br>- D1005.2<br>F18A11.2<br>F33H2.8<br>- F48D6.4<br>F54B11.10<br>- F57B9.1<br>- F58H10.1<br>- H40L08.2<br>- K01A2.4<br>- R148.5<br>T03G6.1<br>T04G9.4<br>T19B10.2<br>- W04A8.4<br>- Y110A2AL.4<br>Y23H5B.8<br>- Y41D4B.6<br>- Y43F8B.2<br>Y6D1A.2<br>- aass-1<br>atf-8<br>- atp-6<br>bus-12<br>bus-17<br>- calu-1<br>- ceeh-1<br>- cey-1<br>- clec-196<br>- cpg-20<br>- cpg-8<br>cpi-2<br>- ctc-1<br>- cutl-2<br>- cyn-6<br>- dao-2<br>dhs-5<br>- dsl-3<br>- dsl-6<br>- eef-1A.2<br>- elt-3<br>- fkb-4<br>- gmap-1<br>- hacl-1<br>- lam-3<br>lpr-1<br>lpr-3<br>lpr-4<br>lpr-5<br>lpr-6<br>- lron-8<br>- mab-21<br>- maoc-1<br>mlt-11<br>- mps-2<br>- ndk-1<br>nhr-127<br>nhr-270<br>- nlp-31<br>nmy-1<br>- pah-1<br>- pcbd-1<br>- pck-2<br>- pgph-3<br>- qdpr-1<br>- rabr-1<br>- ret-1<br>- rmd-2<br>rml-3<br>spp-13<br>- srlf-1<br>sym-2<br>- ttr-20<br>- ugt-60<br>wrt-10 | - B0365.9<br>- B0393.5<br>C02E7.6<br>- C06C3.4<br>- C06E7.88<br>C29A12.6<br>C35A5.11<br>- C48E7.1<br>- C53A5.2<br>C54C8.4<br>D1014.5<br>- F11E6.9<br>F13A7.12<br>- F15B9.8<br>F15H10.8<br>F17B5.6<br>F18A11.2<br>- F26E4.3<br>- F31C3.6<br>F33H2.8<br>F35G2.5<br>F37A4.3<br>- F46E10.2<br>- F46F11.7<br>- F46G11.6<br>F49D11.6<br>F54B11.10<br>F58E6.13<br>- F58H1.2<br>K01D12.1<br>- K02E10.4<br>- K08B12.1<br>- T01D1.8<br>T03G6.1<br>- T05H10.3<br>T06A1.1<br>- T19C4.1<br>- Y102A11A.5<br>- Y41D4B.6<br>- ZC449.2<br>abu-13<br>act-4<br>- aqp-2<br>arf-5<br>asns-2<br>- best-14<br>bgnt-1.2<br>bre-1<br>bus-12<br>bus-17<br>bus-8B<br>- calu-1<br>- clec-78<br>cope-1<br>- cpg-8<br>ctg-1<br>cut-2<br>cut-5<br>cyp-42A1<br>- dao-2<br>dex-1<br>dhs-5<br>- dsl-6<br>dtmk-1<br>eef-1A.1<br>eef-2<br>- far-3<br>glf-1<br>nbet-1<br>nhr-127<br>nhr-270<br>- nlp-33<br>nmy-1<br>nstp-3<br>oac-51<br>pst-2<br>- ptr-25<br>rla-1<br>rml-3<br>rocf-1<br>rpl-12<br>rpl-7A<br>samt-1<br>- srlf-1<br>- sym-1<br>sym-2<br>- tag-297<br>- ttr-2<br>- ttr-59<br>- unc-52<br>- vha-2<br>wrt-2 | C01G6.9<br>C02E7.6<br>C02F4.4<br>C14A4.9<br>- C29E4.15<br>C35A5.11<br>C52G5.2<br>F01D5.6<br>F09F9.2<br>F15H10.8<br>F18C5.5<br>F41F3.3<br>F49D11.6<br>- F53B3.5<br>F58H1.2<br>K01D12.1<br>K08B12.1<br>M03B6.3<br>- R06C1.4<br>R07E3.6<br>T03D8.6<br>T05H10.3<br>T19A5.3<br>Y102A11A.5<br>- Y110A2AL.4<br>Y37A1B.7<br>Y54G2A.76<br>- Y58A7A.2<br>Y65B4BL.1<br>Y73E7A.8<br>ZC449.1<br>ZK154.1<br>abu-13<br>- act-4<br>- ant-1.1<br>- asns-2<br>- atp-6<br>bus-8B<br>cnc-3<br>- cpg-24<br>- ctc-3<br>- ctg-1<br>cut-2<br>dpy-17<br>- dsl-6<br>- eef-1A.1<br>- eef-1B.1<br>- eef-2<br>glf-1<br>grl-15<br>grl-5<br>grl-7<br>- hch-1<br>- inx-12<br>- inx-13<br>mab-7<br>- mlt-11<br>- mlt-8<br>mlt-9<br>- mps-2<br>- ndk-1<br>- nduo-6<br>- nmy-1<br>- noah-1<br>- noah-2<br>nstp-3<br>osm-8<br>- pab-1<br>- pcbd-1<br>- pdi-2<br>- pdi-6<br>- rla-1<br>- rml-3<br>- rpl-12<br>- rpl-27<br>- rpl-7A<br>- rps-16<br>- sma-1<br>spp-13<br>sups-1<br>- sym-1<br>- sym-2<br>- tag-273<br>- tct-1<br>txdc-12.1<br>wrt-1<br>- wrt-10<br>wrt-2 | - C01G6.9<br>C02E7.7<br>- C02F4.4<br>C10F3.7<br>- C14A4.9<br>- C26B9.3<br>- C35A5.11<br>- C52G5.2<br>- F09F9.2<br>F10D11.6<br>- F15B9.8<br>- F18C5.5<br>- F30A10.2<br>- F33D4.6<br>F41F3.3<br>F53F1.4<br>- F58H1.2<br>- H03E18.1<br>H10E21.4<br>K01D12.9<br>- K08B12.1<br>- M03B6.3<br>- R07E3.6<br>R07E5.4<br>R12E2.14<br>R12E2.15<br>R12E2.7<br>R74.2<br>- T19B10.2<br>- T26C5.2<br>- Y102A11A.5<br>- Y11D7A.5<br>- Y37A1B.7<br>Y47D7A.13<br>- Y54G2A.76<br>- Y65B4BL.1<br>- Y71G12B.25<br>- Y73E7A.8<br>- ZC123.1<br>- ZC449.1<br>ZK180.5<br>- abu-13<br>- aqp-2<br>- cbn-1<br>cht-5<br>- cnc-3<br>col-104<br>col-107<br>col-109<br>col-125<br>col-14<br>col-48<br>col-65<br>col-74<br>col-77<br>col-97<br>cut-3<br>- cut-6<br>cutl-14<br>- dpy-14<br>- dpy-17<br>- glf-1<br>- grl-10<br>- grl-15<br>grl-16<br>- grl-5<br>- grl-7<br>- hhat-1<br>- lpr-3<br>- mab-7<br>- mlt-9<br>- osm-8<br>ram-2<br>- rml-3<br>- spp-13<br>- sqt-3<br>- sups-1<br>- txdc-12.1<br>- wrt-1<br>- wrt-2 | - C01G6.9<br>C02E7.6<br>C10F3.7<br>C15C6.1<br>C29A12.6<br>C35A5.11<br>- C38C6.3<br>- C53A5.2<br>D1014.5<br>- F09F9.2<br>F11A10.6<br>F11A5.5<br>F13A7.12<br>F15H10.8<br>F23H12.7<br>- F26E4.3<br>- F30A10.2<br>F35G2.5<br>F36F12.1<br>F37A4.3<br>F47B8.5<br>F49D11.6<br>F58E6.13<br>- F58H1.2<br>H23N18.5<br>K01D12.1<br>- K01D12.9<br>- K08B12.1<br>R05G9R.1<br>- R12E2.15<br>- R74.2<br>T01D1.4<br>T06A1.1<br>T08G5.3<br>W08G11.1<br>- Y102A11A.5<br>- Y37A1B.7<br>Y47D7A.13<br>- Y64H9A.2<br>Y75B12B.11<br>ZK381.2<br>ZK829.3<br>abu-13<br>arf-1<br>bgnt-1.2<br>bgnt-1.3<br>bre-1<br>bus-12<br>bus-17<br>bus-4<br>bus-8B<br>- clec-230<br>cnc-3<br>- col-165<br>col-65<br>- col-74<br>cpg-23<br>cpg-7<br>cut-2<br>- cut-3<br>cut-5<br>dex-1<br>dpy-14<br>dpy-17<br>dtmk-1<br>- grd-5<br>grl-16<br>lips-3<br>- lpr-7<br>mlt-9<br>- nlp-31<br>- nlp-33<br>nstp-3<br>ntp-1<br>oac-51<br>pst-2<br>rocf-1<br>samt-1<br>- sqt-3<br>swt-5<br>- unc-52<br>- wrt-1 | - B0365.9<br>- C06E7.88<br>- C10F3.7<br>- C18E9.2<br>C38C6.3<br>- C48E7.1<br>C50D2.1<br>- F11E6.9<br>- F26E4.3<br>F26G1.5<br>F30A10.2<br>- F31D4.5<br>F33A8.7<br>F41F3.3<br>- F46G11.6<br>- F52A8.1<br>F56D3.1<br>- F58H1.2<br>- F59B10.5<br>K01D12.1<br>- K08B12.1<br>- M03B6.3<br>- R07E3.6<br>R74.2<br>T05H10.3<br>- Y102A11A.5<br>- Y54F10AM.6<br>- Y54G2A.76<br>Y64H9A.2<br>- Y73E7A.8<br>ZC123.1<br>- ZC449.2<br>abu-13<br>- cav-1<br>- clec-180<br>clec-230<br>- col-125<br>- col-14<br>col-165<br>- col-65<br>col-74<br>- col-77<br>- ctc-3<br>cut-2<br>cut-3<br>- dao-2<br>dpy-14<br>- gln-3<br>gmap-1<br>grl-10<br>grl-15<br>- grl-16<br>grl-5<br>grl-7<br>- hmg-1.1<br>lgc-22<br>lips-3<br>lon-8<br>- lpr-1<br>- lpr-3<br>- lpr-4<br>- lpr-5<br>- lpr-6<br>lpr-7<br>mlt-9<br>pqn-32<br>- spp-13<br>sqt-3<br>- tag-297<br>- ttr-59<br>- wrt-1<br>wrt-10<br>- wrt-2 |

**Fig. S20. Gene lists from selected basis vectors of 3-D spaces 6, 9, 12, 13, and 18.** Genes with negative loadings are indicated with a minus sign. Time ranges are the 10th to 90th percentiles of time points represented by cells assigned to each space. Genes involved in morphology, molting, and the cuticle are highlighted in yellow (based on expert knowledge of PWS verified by annotations in WormBase (14) and a table of molt-related genes in (15)), and genes from the nuclear hormone receptor, warthog, and groundhog-like families are highlighted in blue.

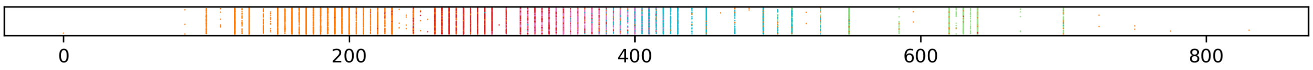


**Fig. S21. Strip plot of embryo time colored by cluster assignment.** The strip plot of embryo time shown in Fig. 4 was randomly perturbed by a Gaussian noise with a standard deviation of 3 minutes for visibility of overlapping points. The unmodified plot is shown here.

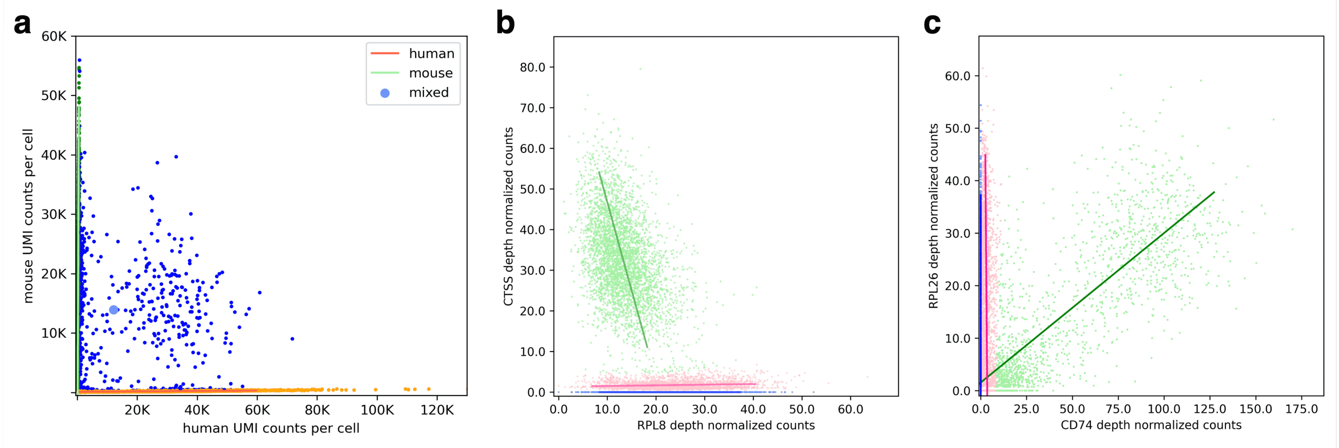


**Fig. S22. Further applications in RNA sequencing. a,** Analysis of a human-mouse single-cell mixture experiment (16) with two lines and a point. **b,** Scatterplot of RPL8 and CTSS expression in (17). $R^2 = 0.37$ overall. **c,** Scatterplot of CD74 and RPL26 expression in (18). $R^2 = 0.113$ overall, implying a little to no linear relationship measured by simple correlation despite clear structure in the data.

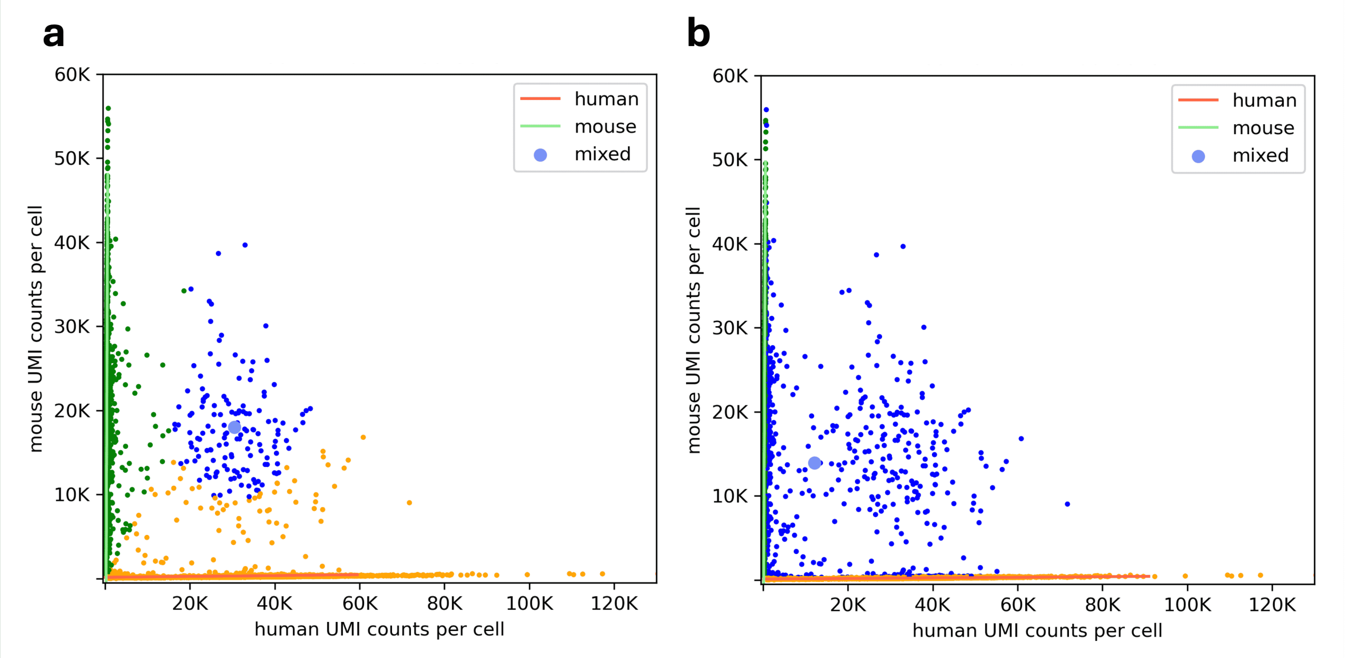


**Fig. S23. Closest and probabilistic assignment yield different outcomes in a barnyard plot.** Data is from (16). **a,** Setting the noise parameter equal for all spaces yields an estimate of 1.1% mixed cells **b,** Probabilistic assignment yields 6.7% mixed cells (reproduced from Fig. S22a).

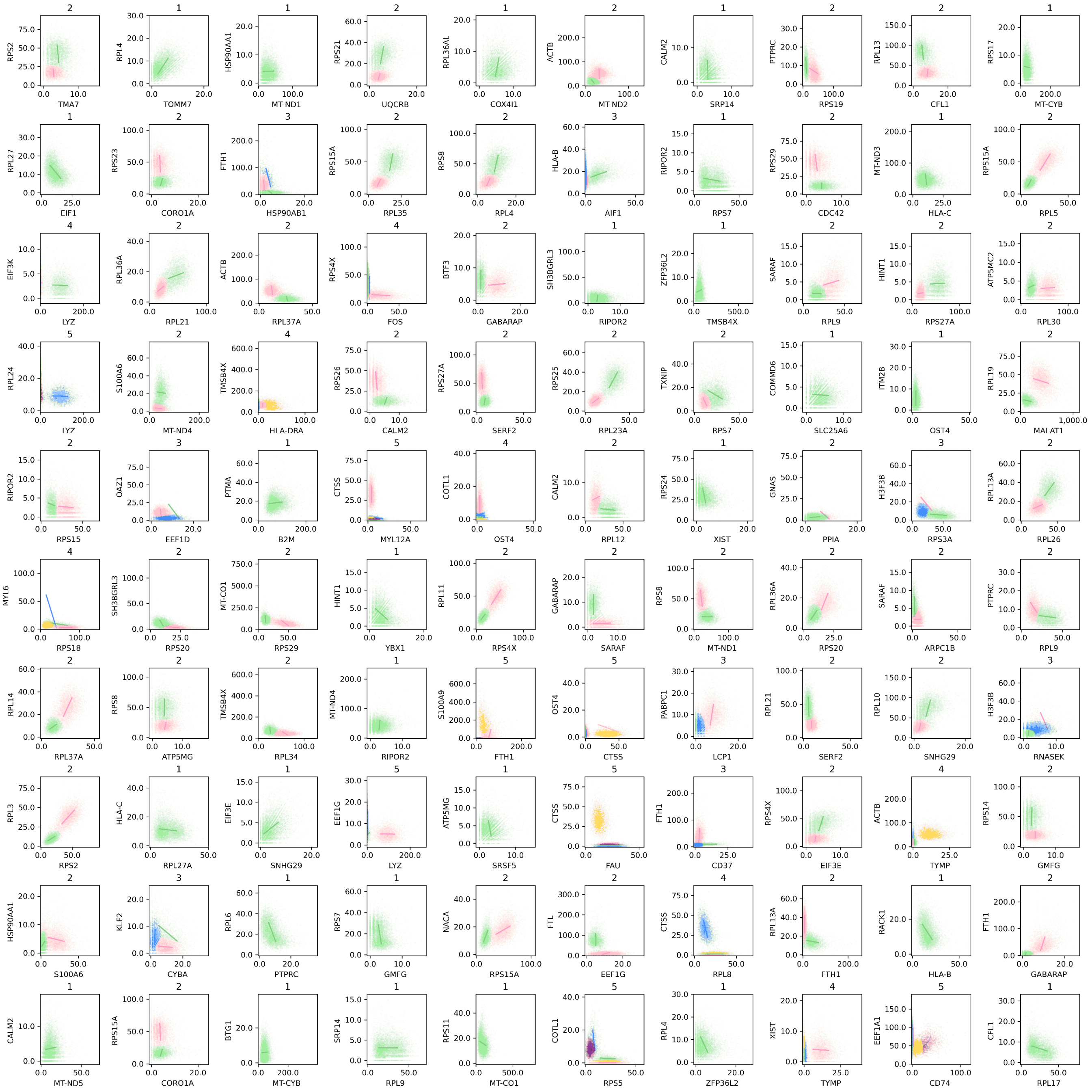


**Fig. S24. 100 random examples of clustering with selection of $k$ via the ICL.** Starting from $k$ = 1, $k$ was incremented if the ICL improved, with a maximum of $k$ = 5. The subplot title for each gene pair indicates the selected value of $k$.

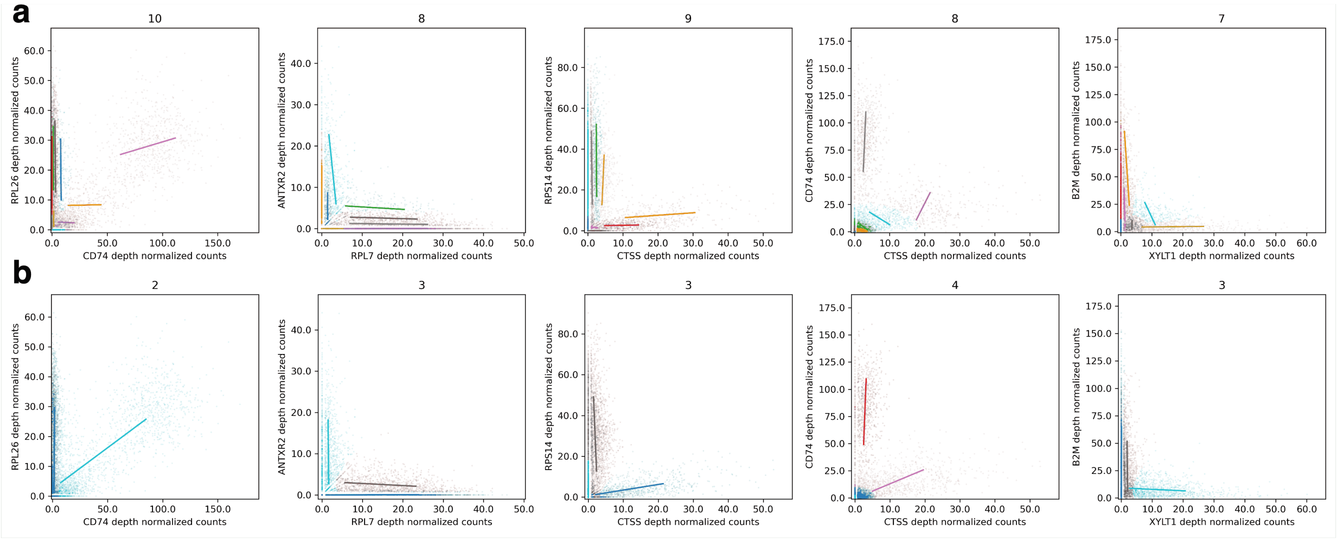


**Fig. S25. Use of ICL on 5 selected gene pairs from (18).** **a,** ICL still leads to some overclustering. Starting from $k$ = 1, $k$ was incremented if the ICL improved, with a maximum of $k$ = 20. The subplot title for each gene pair indicates the selected value of $k$. **b,** Automated elbow detection using the kneedle algorithm (19) on the ICL values from (a) provides better model selection. Subplot title again indicates the selected value of $k$. Remaining overclustering is due to fitting dropout.

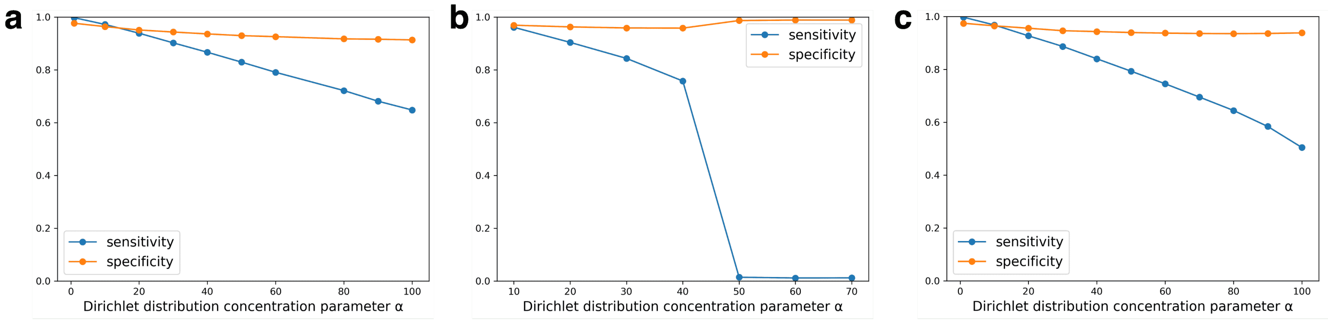


**Fig. S26. Robustness to variation of Dirichlet concentration parameter.** **a,** $\sigma_j \sim \mathrm{Halfnorm}(0, 0.03^2)$. **b,** Empirical distribution of site-specific variances (reproduced from Fig. 3 **b,** Empirical distribution of site specific variances with the top 0.29% of variances ($\sigma_j > 0.2$) removed.

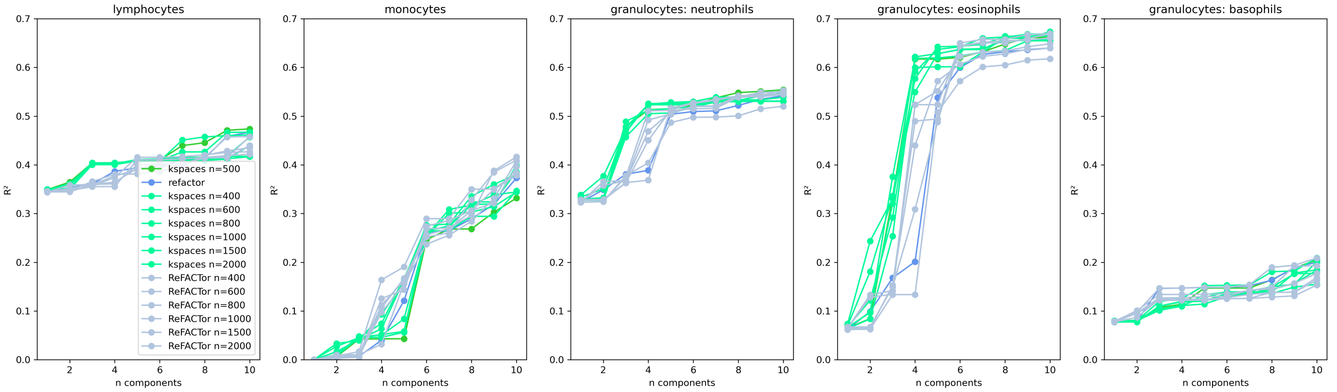


**Fig. S27. Varying n, the number of selected sites, on the GALA II data.** The threshold for selecting informative sites is the standard deviation of noise, 0.053.

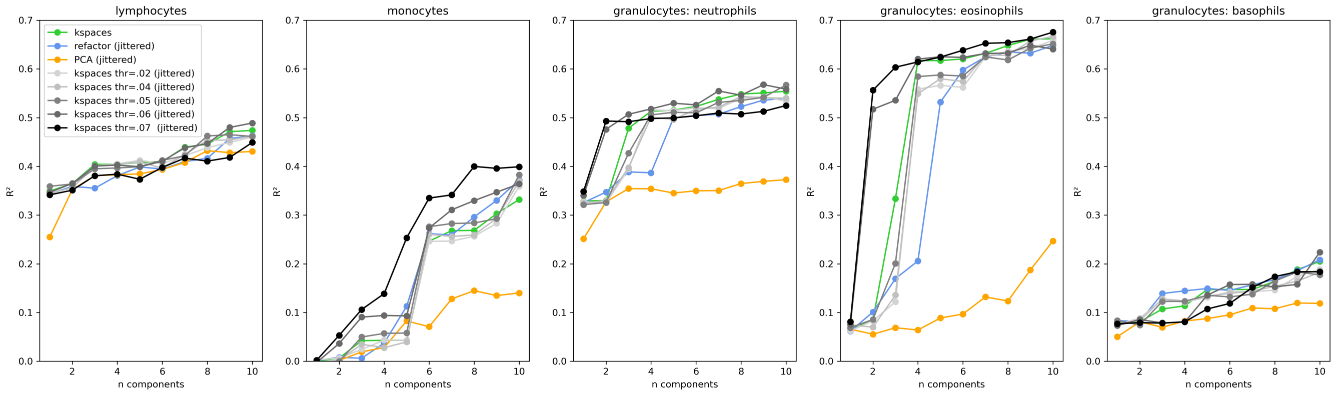


**Fig. S28. Varying the minimum standard deviation threshold for site selection in *k*-spaces.** $n$ is held constant at 500 sites. Our default choice is the standard deviation of noise, which is 0.053 here.

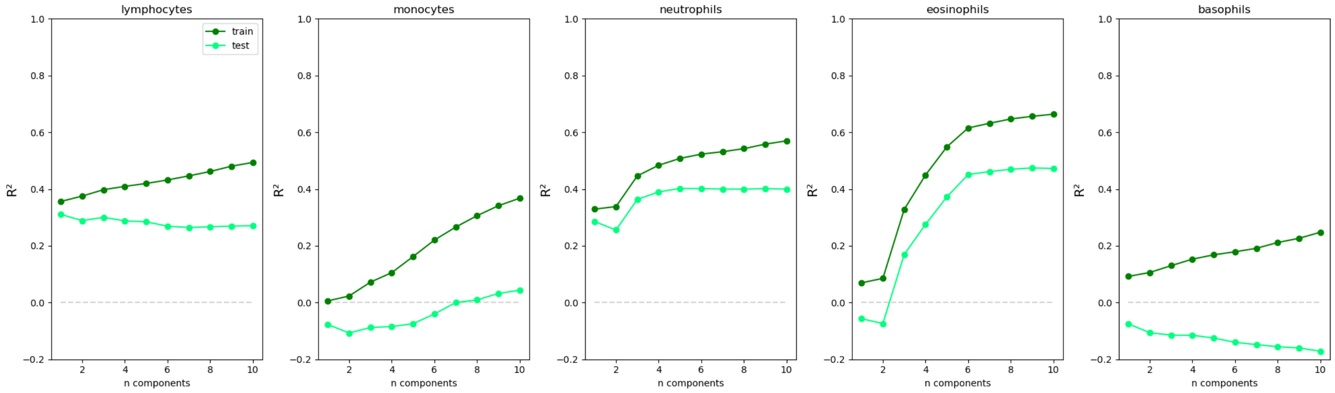


**Fig. S29. Cross validation as in (10).** $R^2$ values from linear regression are reported as a function of the number of principal components used, averaged over 50 runs. For each run, the subset of the GALA II data with cell proportion measurements passing the preprocessing filters in (11) was randomly split into training (n = 50) and test data (n = 30). *k*-spaces clustering with a 4-D and 0-D space with equal noise parameters was run on the training data to obtain informative sites as well as the principal components of those sites in the training data. For the test sets, the same sites and the principal components obtained from the training data were used for regression.

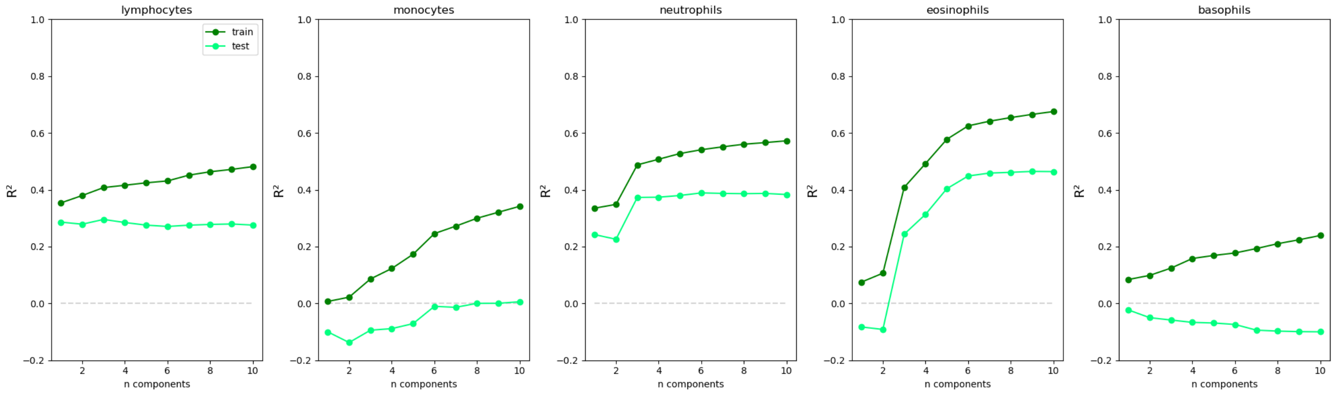


**Fig. S30. Cross validation as in (10) with a 2-D space for DMRs**. The same procedure was used as in Fig. S29 but *k*-spaces clustering with a 2-D space was used instead of 4-D.

| Cell Type | 10th percentile | 25th | 50th | 75th | 90th |
|---|---|---|---|---|---|
| Lymphocytes | 21.8 | 27.8 | 32.7 | 40.2 | 43.9 |
| Monocytes | 4.7 | 5.7 | 6.6 | 7.9 | 9.5 |
| Neutrophils | 40.0 | 47.4 | 54.8 | 61.3 | 68.5 |
| Eosinophils | 1.2 | 1.8 | 4.0 | 6.5 | 9.9 |
| Basophils | 0.3 | 0.4 | 0.5 | 0.5 | 0.7 |

**Table S1. GALA II cell type variation n = 80.**

### References


1. Carl Eckart and Gale Young. The approximation of one matrix by another of lower rank. *Psychometrika*, 1:211–218, September 1939. doi: 10.1007/BF02288367.
2. Tara Chari and Lior Pachter. The specious art of single-cell genomics. *PLOS Computational Biology*, 19(8):e1011288, August 2023. ISSN 1553-7358. doi: 10.1371/journal.pcbi.1011288.
3. Laurens van der Maaten and Geoffrey Hinton. Visualizing Data using t-SNE. *Journal of Machine Learning Research*, 9:2579–2605, 2008.
4. Leland McInnes, John Healy, and James Melville. UMAP: Uniform Manifold Approximation and Projection for Dimension Reduction, September 2020. arXiv:1802.03426 [stat].
5. Michael Collins, Sanjoy Dasgupta, and Robert E. Schapire. A Generalization of Principal Component Analysis to the Exponential Family. In Thomas G. Dietterich, Suzanna Becker, and Zoubin Ghahramani, editors, *Advances in Neural Information Processing Systems 14*, pages 617–624. The MIT Press, November 2002. ISBN 978-0-262-27173-8. doi: 10.7551/mitpress/1120.003.0084.
6. Michael E Tipping and Christopher M Bishop. Mixtures of Probabilistic Principal Component Analyzers. *Neural Computation*, 11(2):443–482, February 1999. doi: 10.1162/089976699300016728.
7. Andrey N. Kolmogorov. Erkältung eines Borelschen Paradoxons. In *Grundbegriffe der Wahrscheinlichkeitsrechnung*, pages 44–45. Julius Springer, Berlin, 1933.
8. Lovisa E. Reinius, Nathalie Acevedo, Maaike Joerink, Göran Pershagen, Sven-Erik Dahlén, Dario Greco, Cilla Söderhäll, Annika Scheynius, and Juha Kere. Differential DNA Methylation in Purified Human Blood Cells: Implications for Cell Lineage and Studies on Disease Susceptibility. *PLoS ONE*, 7(7):e41361, July 2012. ISSN 1932-6203. doi: 10.1371/journal.pone.0041361.
9. Vikas Trivedi, Harry M. T. Choi, Scott E. Fraser, and Niles A. Pierce. Multidimensional quantitative analysis of mRNA expression within intact vertebrate embryos. *Development*, 145(1):dev156869, January 2018. ISSN 1477-9129, 0950-1991. doi: 10.1242/dev.156869.
10. Shijie C Zheng, Stephan Beck, Andrew E Jaffe, Devin C Koestler, Kasper D Hansen, Andres E Houseman, Rafael A Irizarry, and Andrew E Teschendorff. Correcting for cell-type heterogeneity in epigenome-wide association studies: revisiting previous analyses. *Nature Methods*, 14(3):216–217, March 2017. ISSN 1548-7091, 1548-7105. doi: 10.1038/nmeth.4187.
11. Elior Rahmani, Noah Zaitlen, Yael Baran, Celeste Eng, Donglei Hu, Joshua Galanter, Sam Oh, Esteban G Burchard, Eleazar Eskin, James Zou, and Eran Halperin. Sparse PCA corrects for cell type heterogeneity in epigenome-wide association studies. *Nature Methods*, 13(5):443–445, May 2016. ISSN 1548-7091, 1548-7105. doi: 10.1038/nmeth.3809.
12. Esteban Burchard. Differential dna methylation in latino population. https://www.ncbi.nlm.nih.gov/geo/query/acc.cgi?acc=GSE77716, 2 2016.
13. Jonathan S. Packer, Qin Zhu, Chau Huynh, Priya Sivaramakrishnan, Elicia Preston, Hannah Dueck, Derek Stefanik, Kai Tan, Cole Trapnell, Junhyong Kim, Robert H. Waterston, and John I. Murray. A lineage-resolved molecular atlas of *C. elegans* embryogenesis at single-cell resolution. *Science*, 365(6459):eaax1971, September 2019. ISSN 0036-8075, 1095-9203. doi: 10.1126/science.aax1971.
14. Paul W Sternberg, Kimberly Van Auken, Qinghua Wang, Adam Wright, Karen Yook, Magdalena Zarowiecki, Valerio Arnaboldi, Andrés Becerra, Stephanie Brown, Scott Cain, Juancarlos Chan, Wen J Chen, Jaehyoung Cho, Paul Davis, Stavros Diamantakis, Sarah Dyer, Dionysis Grigoriadis, Christian A Grove, Todd Harris, Kevin Howe, Ranjana Kishore, Raymond Lee, Ian Longden, Manuel Luypaert, Hans-Michael Müller, Paulo Nuin, Mark Quinton-Tulloch, Daniela Raciti, Tim Schedl, Gary Schindelman, and Lincoln Stein. WormBase 2024: status and transitioning to Alliance infrastructure. *GENETICS*, 227(1):iyae050, May 2024. ISSN 1943-2631. doi: 10.1093/genetics/iyae050.
15. Vladimir Lažetić and David S. Fay. Molting in *C. elegans*. *Worm*, 6(1):e1330246, January 2017. ISSN 2162-4054. doi: 10.1080/21624054.2017.1330246.
16. 10x Genomics. 10k 1:1 mixture of human hek293t and mouse nih3t3 cells, chromium gem-x single cell 3', single gene expression dataset analyzed using cell ranger v8.0.0. https://www.10xgenomics.com/datasets/10k-hgmm-3p-gemx, 4 2024. Accessed on October 2, 2024.
17. 10x Genomics. 10k human pbmcs, 3' v3.1 chromium x (without intronic reads), single gene expression dataset analyzed using cell ranger v6.1.2. https://www.10xgenomics.com/datasets/10k-human-pbmcs-3-v3-1-chromium-x-without-introns-3-1-high, 3 2022. Accessed on October 2, 2024.
18. 10x Genomics. 5k human pbmcs (donor 4), universal 3' gene expression dataset analyzed using cell ranger 9.0.0. https://www.10xgenomics.com/datasets/5k_Human_Donor4_PBMC_3p_gem-x, 11 2024. Accessed on June 23, 2024.
19. Ville Satopaa, Jeannie Albrecht, David Irwin, and Barath Raghavan. Finding a "Kneedle" in a Haystack: Detecting Knee Points in System Behavior. In *ICDCSW '11: Proceedings of the 2011 31st International Conference on Distributed Computing Systems Workshops*, pages 166–171, Minneapolis, MN, USA, June 2011. doi: 10.1109/ICDCSW.2011.20.
